# Structure-based Evolution of G protein-biased μ-opioid Receptor Agonists

**DOI:** 10.1101/2022.03.22.485330

**Authors:** Haoqing Wang, Florian Hetzer, Weijiao Huang, Qianhui Qu, Justin Meyerowitz, Jonas Kaindl, Harald Hübner, Georgios Skiniotis, Brian K. Kobilka, Peter Gmeiner

## Abstract

The μ-opioid receptor (μOR) is the major target for opioid analgesics. Activation of μOR initiates signaling through G protein pathways as well as through β-arrestin recruitment. μOR agonists that are biased towards G protein signaling pathways demonstrate diminished side effects. PZM21, discovered by computational docking, is a G protein biased μOR agonist. Here we report the cryoEM structure of PZM21 bound μOR in complex with G_i_ protein. Structure-based evolution led to multiple PZM21 analogs with more pronounced G_i_ protein bias and increased lipophilicity to improve CNS penetration. Among them, FH210 shows extremely low potency and efficacy for arrestin recruitment. We further determined the cryoEM structure of FH210 bound to μOR in complex with G_i_ protein and confirmed its expected binding pose. The structural and pharmacological studies reveal a potential mechanism to reduce β-arrestin recruitment by the μOR, and hold promise for developing next-generation analgesics with fewer adverse effects.

**Table of Contents Graphical Abstract:** 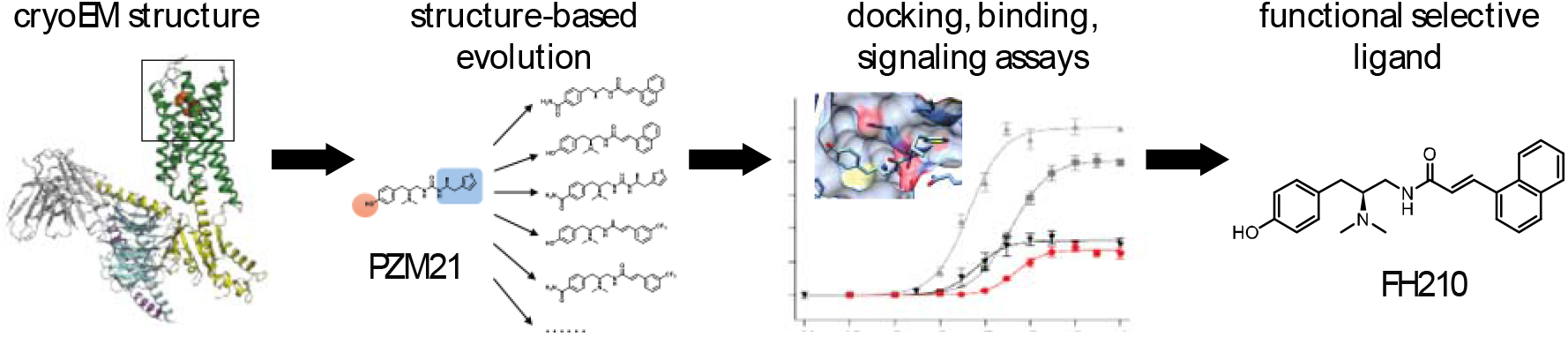

We obtained cryoEM structures of the μ-opioid receptor (μOR) bound to the lead compound PZM21 and the newly developed agonist FH210 to understand the mechanism of their biased signaling and to guide the evolution of next-generation analgesics with fewer adverse effects.

---

Addiction to opioid analgesics has led to the opioid crisis, causing more than 70,000 death per year.^[1]^ Consequently, there has been a search for new alternative analgesic drugs with lower addictive potential and reduced side effects^[2]^. Computational docking followed by structure-based optimization has led to the discovery of PZM21, a G protein biased μOR partial agonist producing analgesia with diminished side effects in mice.^[3]^ *In vivo* experiments with PZM21 showed no rewarding or reinforcing effects^[4]^ and less respiratory depression compared to morphine, creating a wider therapeutic window.^[5]^ Nevertheless, the blood-brain ratio of PZM21 in mice indicates only moderate receptor occupancy in the central nervous system (CNS) after systemic administration, likely due to its relatively high polarity (log P: 2.9).^[3]^ Moreover, PZM21 still evokes tolerance and withdrawal symptoms.^[4]^ As the recruitment of β-arrestin-2 to the μOR has been shown to correlate with the development of tolerance *in vivo*^[6]^, we seek derivatives of PZM21 featuring more pronounced G protein bias and increased lipophilicity^[7]^.

Interestingly, for several family A GPCRs, ligands addressing the classical orthosteric pocket and extending into the extracellular vestibule generate functional selectivity. It appears that additional interactions with the extracellular part of the transmembrane helices and loop regions^[8]^ confer a conformational restriction of the receptor which impacts the stability of different receptor-transducer complexes. Compared to classical opioids, the new G protein-biased ligands PZM21 and the clinically approved drug oliceridine (TRV130) may also interact with amino acids in the extracellular vestibule.^[9]^

In this study, we obtained a high-resolution structure of the μOR bound to PZM21 to understand the mechanism of its biased signaling and used structure-based evolution to develop analogs with enhanced G protein bias. We also obtained a high-resolution structure of one of these analogs (FH210) having a further increase in G protein bias relative to PZM21, largely due to a decrease in potency and efficacy in arrestin recruitment. The structure reveals differences between the binding of PZM21 and FH210 that may be responsible for the reduced arrestin recruitment.

To enable a structure-based design, we first determined a high-resolution structure of PZM21 bound to μOR in complex with G_i_ protein. Using single-particle cryoEM, we obtained the PZM21-μOR-G_i_ protein complex structure at 2.9Å resolution (Figure 1A, Figure S1A, Figure S2 and Table S1). The map reveals well-defined densities for the amino acids forming the orthosteric pocket as well as for the ligand PZM21 (Figure 1B, Figure S2D). The structure revealed two common features that were previously observed in the μOR in complex with DAMGO^[10]^, BU72^[11]^ and β-FNA^[12]^: a salt bridge between the basic amine of the drug and D147^3.32^ and a position of the phenol hydroxy group in close proximity to H297^6.52^ (Figure 1B). In the structures of BU72 and β-FNA bound μOR, two water molecules could be resolved, mediating a hydrogen bond network between the phenol hydroxyl group and H297^6.52^.^[11-12]^ In our structure, the phenol moiety of PZM21 showed an analogous spatial arrangement. However, mediating water molecules could not be resolved. Hence, we performed molecular dynamics (MD) simulations to assess the stability of the PZM21 pose and the contribution of a water network. During the simulations, PZM21 remained close to its initially modeled pose and stable water-mediated interactions between the phenol of PZM21, H297^6.52^ and the backbone carbonyl of K233^5.39^ could be observed (Figure S3). This polar network was further confirmed by chemical synthesis and functional characterization of a PZM21 carboxamide analog bioisosterically replacing the phenol hydroxy group and the mediating water (FH310, Table S2). This structural congener showed an almost five-fold increase in binding affinity determined by radioligand binding studies (FH310, Table S3).

**Figure 1:**
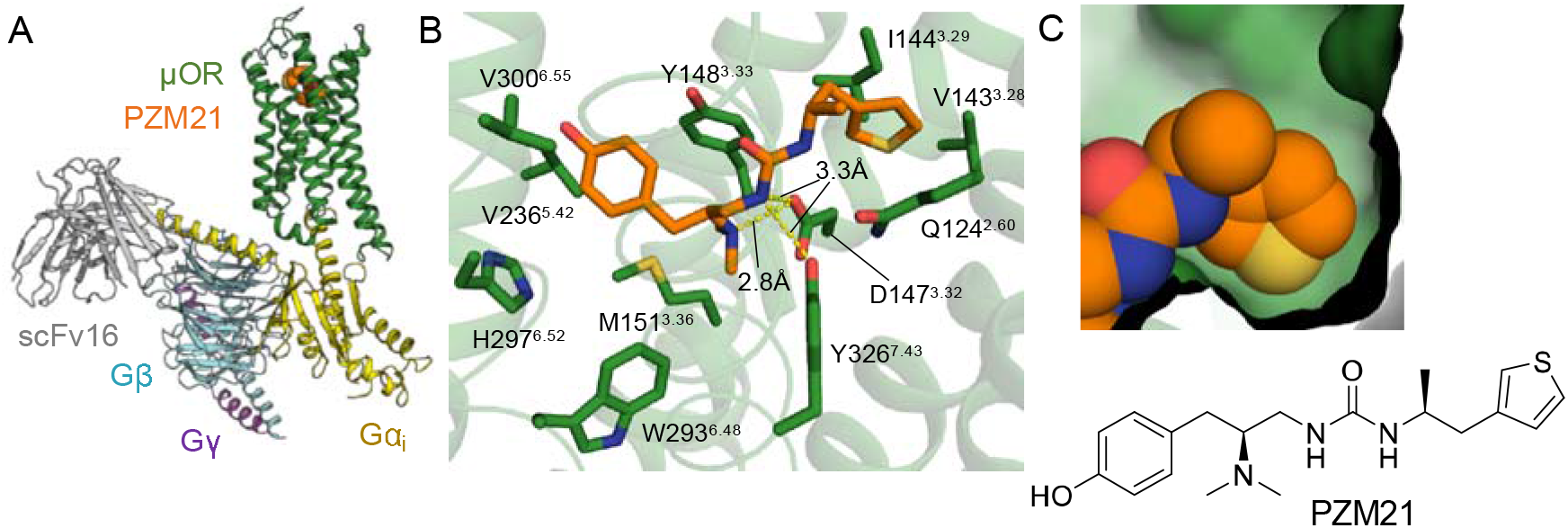
A) Structure of the μOR–G_i_ complex bound to PZM21 colored by subunit. Green, μOR; orange, PZM21; gold, Gα_i_; cyan, Gβ; purple Gγ; gray scFv16. B) View of PZM21 in the binding pocket. PZM21 forms polar contacts to D147^3.32^ and Y326^7.43^ and the phenol group is near H297^6.52^ suggesting that PZM21 forms a water-mediated interaction with H297^6.52^, as observed in previous structures of the μOR. C) Surface presentation of the thiophene group of PZM21 in the lipophilic vestibule of the μOR.

The cryoEM structure of PZM21 bound to the μOR shows that the thiophenylalkyl moiety interacts with a lipophilic vestibule formed by the extracellular ends of TM2 and TM3 and ECL1 (Figure 1C, Figure S4A). We reasoned that PZM21 analogs showing higher complementarity with the extracellular vestibule, more specifically to V143^3.28^, I144^3.29^, W133^ECL1^ and N127^2.63^, may confer an increase of G protein bias compared with unbiased agonists such as DAMGO and BU72 (Figure S4A). Following this hypothesis, we designed 19 test compounds guided by the PZM21-μOR cryoEM structure exchanging the urea unit and the thiophenylalkyl substituent by manually selected bioisosteric fragments. After molecular docking and careful inspection of the complexes, we selected those derivatives that had a reasonable binding pose, a promising docking score and an increase of lipophilicity for synthesis (Table S2). The structure-guided design led us to acrylamide analogs of PZM21. The compounds lack one of the urea NH groups and, hence, have only two instead of three hydrogen bond donors, which may increase CNS penetration.

To obtain candidate PZM21 analogs, enantiopure acrylic and acetylenic amides were synthesized starting from the amino acid derived precursor *L*-tyrosine amide (**1**). A reductive dimethylation with formaldehyde and sodium triacetoxyborohydride followed by borane reduction of the amide functionality led to the building block **2**. Acrylic and acetylenic amides were synthesized via BOP- or PyBOP-promoted amide coupling reaction of the primary amine **2** and commercially available acrylic and acetylenic acids (compounds of type **3**, Scheme 1, Scheme S1).

**Scheme 1.**
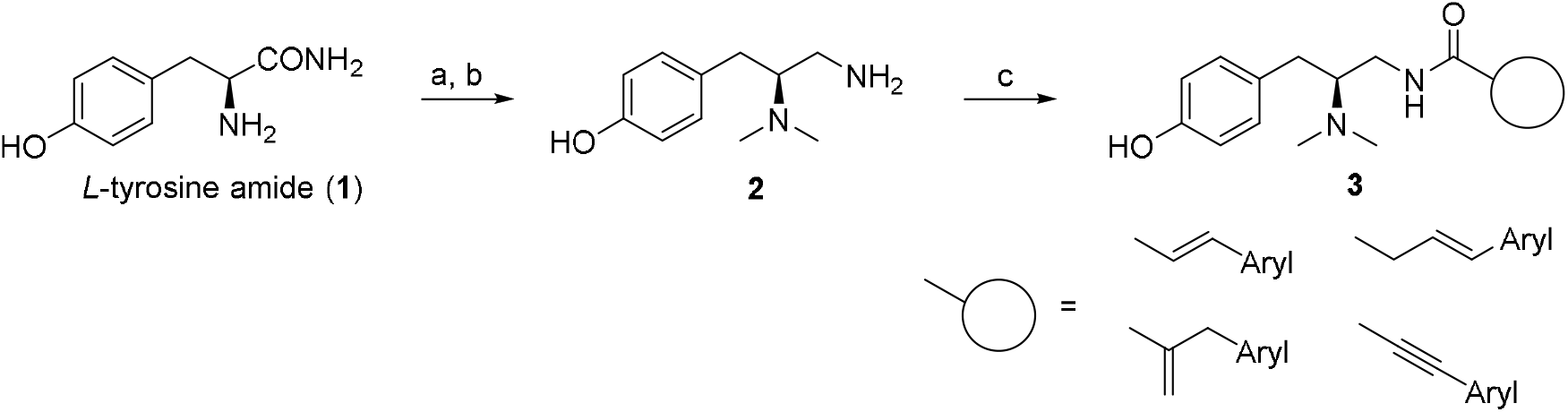
Synthesis of PZM21 analogs. Reagents and conditions: a) formaldehyde, sodium triacetoxyborohydride, water, acetonitrile, −10°C, 5 min (> 99 %); b) borane-THF complex, THF, 0°C to reflux, 6 h (43 %), c) carboxylic acid, BOP or PyBOP, triethylamine, DMF, r.t., 0.5-24 h (56-95 %).

Seeking compounds with low *K_i_*-values similar to PZM21 (*K_i_*(μOR)=31 nM), we assessed receptor affinity of these PZM21 analogs for the μOR in a radioligand-binding assay and compared them with the reference drugs fentanyl, morphine and oliceridine (Table S3). Ten out of the eleven novel analogs surpassed the μOR affinity of PZM21. Notably, these PZM21 analogs were also selective for the μOR over δOR and κOR (Table S3). In contrast to PZM21, which binds to the μOR (*K_i_*= 31 nM) and κOR (*K_i_*= 24 nM) with similar affinities, most analogs are selective for the μOR (*K_i_*= 6.6-41 nM) over both δOR (*K_i_*=25-530 nM) and κOR (*K_i_*=58-600 nM) (Table S3).

We then investigated these potent analogs for their signaling profiles. Functional assays were performed to identify compounds capable of activating the μOR-dependent G protein pathway with an E_max_ of at least 90% of fentanyl and high bias over β-arrestin recruitment. Whereas G protein activation was investigated by monitoring IP_1_ accumulation in presence of a co-transfected G_qi_-alpha subunit, β-arrestin-2 recruitment was determined using a fragment complementation assay (DiscoverX PathHunter) (Figure 2A-C, Table S4). Four agonists showed efficacy greater than 90% of fentanyl for G protein activation. Interestingly, for three of these agonists, β-arrestin recruitment was below a detection threshold of 5% compared to fentanyl. To improve assay sensitivity for arrestin and to better discriminate between ligands with low arrestin efficacy, we monitored β-arrestin-2 recruitment in cells coexpressing G protein receptor kinase 2 (GRK2), an intracellular kinase that phosphorylates the C-terminus of the μOR, thereby stabilizing interactions with β-arrestin-2.^[13]^ Following these conditions, we were able to obtain dose-response curves allowing a more precise evaluation of functional bias.

**Figure 2:**
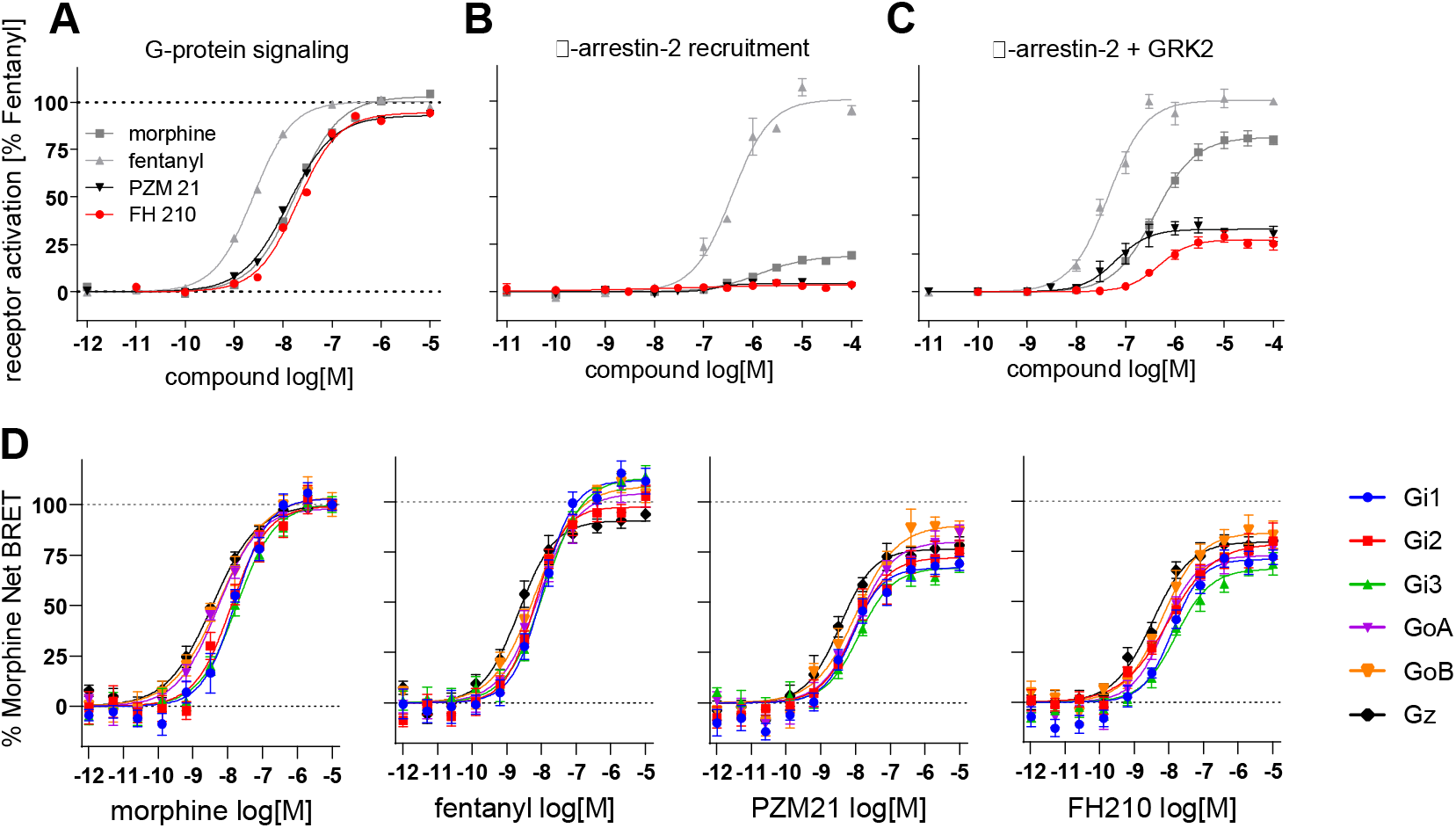
Functional activity of morphine, fentanyl, PZM21 and FH210 at the μOR. A) G-protein signaling measured by IP accumulation assay in cells transfected with the chimeric G protein G_qi_. B,C) β-arrestin-2 recruitment measured by PathHunter assay, with (C) and without (B) GRK2 co-transfection. D) G_i/o/z_ subtypes activation measured by BRET-based assay^[14]^.

Based on the functional characterization of these PZM21 analogs, we identified several candidates with potential clinical utility. In particular, FH210 (log P: 3.9), a naphthyl-substituted acryl amide, showed high complementarity and increased Van der Waals interactions within the lipophilic vestibule while maintaining an overall PZM21-like binding mode, based on our docking results (Figure S4B, Table S2). FH210 is more biased towards the G protein pathway than PZM21. While FH210 stimulates G protein activation to a similar extent as does PZM21, it shows attenuated β-arrestin-2 recruitment in the presence of GRK2 (Figure 2A,C, Table S4): compared to PZM21, the β-arrestin-2 dose-response curve of FH210 in the presence of GRK2 is shifted about one order of magnitude rightwards (EC_50_ = 94 nM and 860 nM respectively) and a decreased efficacy was observed (E_max_ = 35 % and 28 %, respectively) (Figure 2C). Off-target effects of FH210 at 20 other GPCRs were also studied in radioligand binding experiments (Table S5). We detected binding affinities in the micromolar range at these targets, except for the 5-HT_2A_ (*K_i_*=190 nM), α_1A_ (*K_i_*=490 nM), α_2B_ (*K_i_*=610 nM) and D_4.4_ receptor (*K_i_*=940 nM).

GPCR agonists also differed in the efficacy and order of potency for G_α_-subtypes activation. μOR primarily signals through the G_i/o/z_ family of G proteins that include G_i1_, G_i2_, G_i3_, G_oA_, G_oB_ and G_z_. *In vivo* studies have shown that specific G_α_-subtype pathways differentially contribute to μOR-dependent behavioral responses, including tolerance and antinociception^[15]^. We therefore characterized the G_α_-subtypes signaling properties of PZM21 and FH210, using BRET TRUPATH biosensors^[14]^ (Figure 2D, Table S6). For reference compounds we included morphine and fentanyl. In these experiments we observed similar potency and efficacy profiles for PZM21 and FH210. Both are partial agonist at all G_i/o/z_ subtypes with the E_max_ ranging from 67% to 89% of morphine. For PZM21, the EC_50_ ranges from 3.7nM for G_z_ to 14nM for G_i3_. For FH210, the EC_50_ ranges from 3.8nM for G_z_ to 19nM for G_i3_ (Table S6).

To understand the structural basis for the very weak efficacy and potency of FH210 in arrestin recruitment, we obtained the cryoEM structure of FH210 bound to the μOR-G_i_ protein complex. Single-particle cryoEM was performed to obtain a threedimensional map of the FH210-μOR-Gi protein complex at a global resolution of 3.0Å (Figure 3A, Figure S1B, Figure S2 and Table S1). Our cryoEM map includes well-defined densities for the amino acids forming the orthosteric pocket as well as for the ligand FH210 (Figure 3B, Figure S2D). The resolved binding conformation of FH210 closely resembles the pose we obtained from molecular docking experiments using the PZM21-μOR-G_i_ protein complex (Figure 3B and Figure S4B). Overall, we found a striking resemblance between the ligand binding sites of the two resolved structures. The common features include an ionic interaction between the ammonium group of the ligand and D147^3.32^, a hydrogen bond interaction with Y326^7.43^ via the carboxamide NH of FH210 and the proximity of the phenol hydroxy group and H297^6.52^ (Figure 1B and Figure 3B). FH210 also interacts with the lipophilic vestibule like PZM21 (Figure 1C and Figure 3C). MD-simulations show a water-mediated network between the phenol of FH210, H297^6.52^ and K233^5.39^ stabilizing the binding pose (Figure S3).

**Figure 3:**
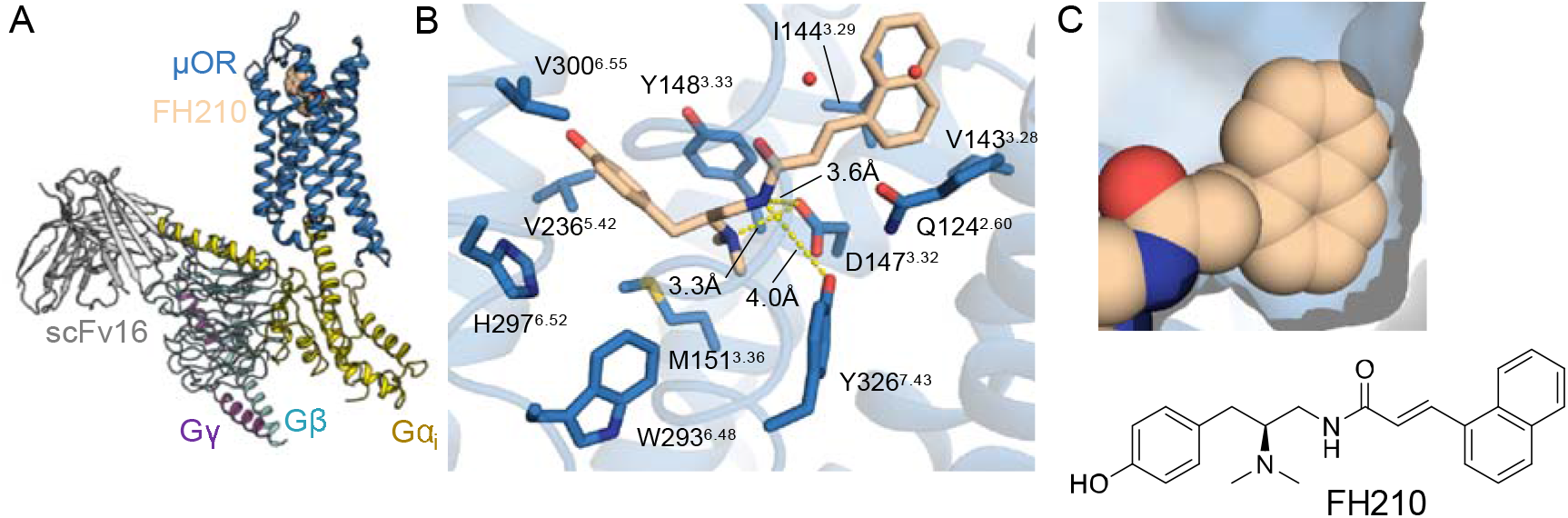
A) Structure of the μOR–G_i_ complex bound to FH210 colored by subunit. Blue, μOR; beige, FH210; gold, Gα_i_; cyan, Gβ; purple Gγ; gray scFv16. B) View of FH210 in the binding pocket. Blue, μOR; beige, FH210. FH210 forms polar contacts with D147^3.32^ and Y326^7.43^ and shows proximity of the phenol group to H297^6.52^ suggesting that the ligand forms a water mediated interaction to H297^6.52^. C) Surface representation of the naphthyl group of FH210 in the lipophilic vestibule of the μOR.

The three-dimensional map of the cryoEM complex revealed densities for two water molecules (Figure S2D). These two water molecules do not have strong interaction with either FH210 or any μOR residues. In MD simulations they did not maintain the position and interactions observed in the cryoEM structure, suggesting that they do not form a distinct water network by themselves.

Besides similarities, we identified the structural features that are unique to the FH210-bound μOR-G_i_ protein complex. We found substantially greater Van der Waals interactions between the naphthyl substituent of FH210 and the lipophilic vestibule formed by TM2, TM3, ECL1 and ECL2. Calculation of the contact surface area between the lipophilic vestibule and the thiophenyl- (PZM21, 124 Å^2^) or the naphthyl-moiety (FH210, 155 Å^2^) revealed a 31 Å^2^ higher contact area between the naphthyl group of FH210 and the receptor (Figure 4A, B). This is particularly attributed to additional contacts with D216^ECL2^, C217^ECL2^, W133^ECL1^ and N127^2.63^ (Figure 4B). In agreement with this, MD simulations showed that the lipophilic vestibule of FH210 bound μOR is more compact compared with PZM21 bound μOR, suggesting that the naphthyl moiety of FH210 is involved in stronger hydrophobic interaction with nearby residues (Figure 4C). The structural differences between the two complexes are primarily found at the extracellular part of the transmembrane helices and loop regions supporting the initial hypothesis that ligand interactions with the extracellular vestibule influence functional bias.

**Figure 4:**
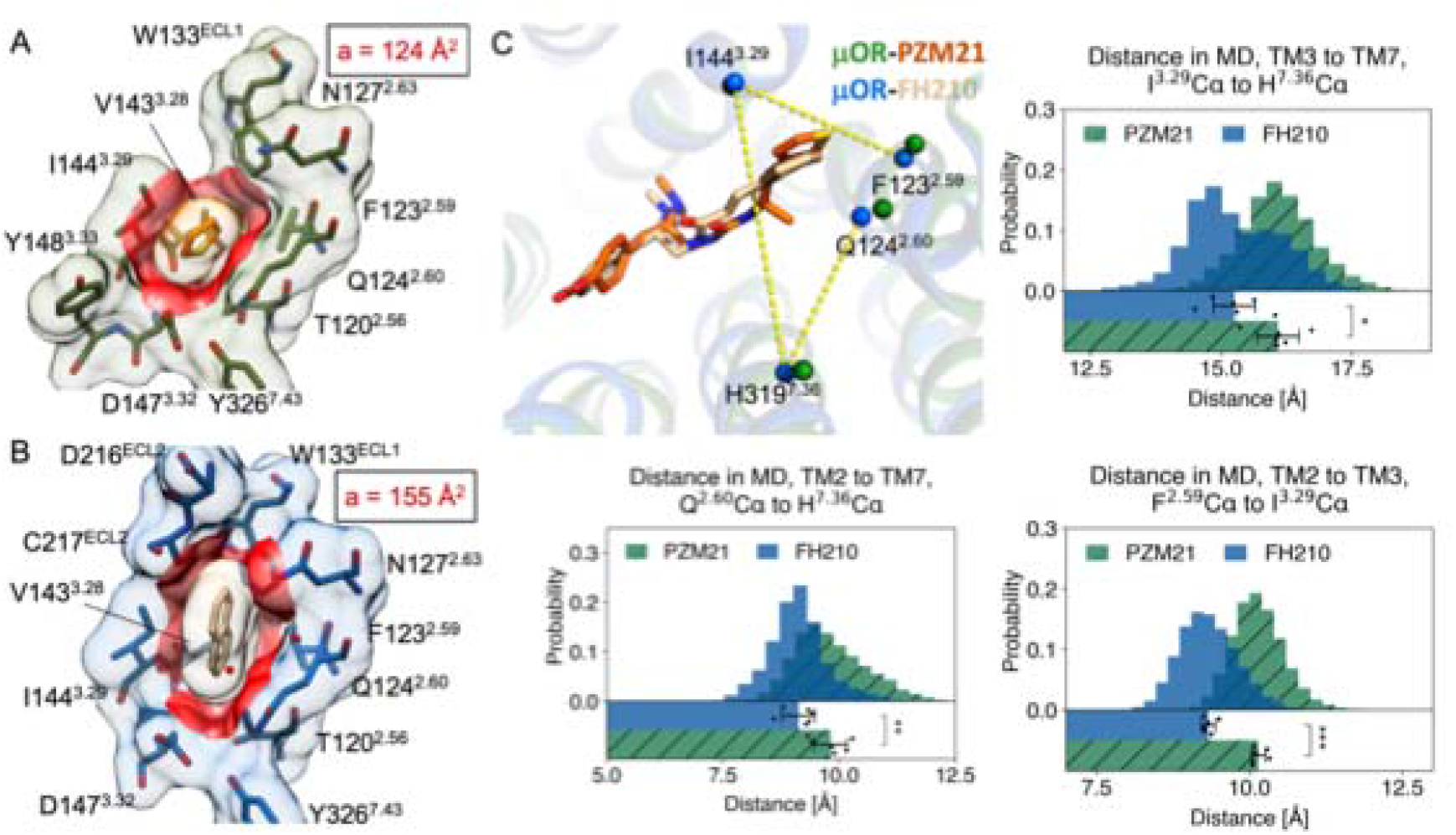
Comparison of μOR bound to PZM21 and FH210. A,B) Contact surface area (red) between the thiophene of PZM21 (orange) and the PZM21-μOR cryoEM complex (green), a = 124 Å^2^; and between the naphthyl of FH210 (beige) and the FH210-μOR cryoEM complex (blue), a = 155 Å^2^. Contact surface area was calculated with a cut-off value of 2Å using UCSF Chimera^[16]^. C) Representative structures extracted from MD simulations of PZM21-μOR and FH210-μOR. The simulations show a more compact lipophilic vestibule for FH210 bound μOR. The distances are measured between the Cα atoms of the labeled residues. The histogram plots on the right shows the distance distribution over all simulations. The bar chart at the bottom of each plot shows the mean and SEM, with the dots representing the individual values. These statistics are based on 6 individual simulations and the first 500ns of each simulation was not included in these analyses.

Recent studies using NMR spectroscopy and MD simulations have suggested a potential mechanism for the μOR biased signaling^[17]^. NMR studies by Cong et al. revealed that G protein biased agonists including PZM21 trigger conformational changes in TM7, ICL1 and H8. MD simulations showed a primary binding pose for PZM21 that penetrates deeper into the binding pocket wedged between TM2 and W293^6.48^, the conserved “rotamer toggle switch”, as well as a transient pose that appears to be similar to the one observed in our structure. In our studies neither PZM21 nor FH210 interact with W293^6.48^, although we cannot exclude the possibility that the alternate binding pose observed by Cong et al. plays a role in stabilizing the μOR in the early stages of complex formation.

The most striking difference between the PZM21 and FH210 binding poses in our study is in the more extensive interactions of the naphthyl moiety and the pocket formed by residues in TM2, TM3 and ECL1 (Figure 4A, B). This may stabilize the μOR in a conformation that is less compatible for interacting with arrestins, however, given that the cytoplasmic surface of the μOR in our structures is stabilized by interactions with nucleotide-free Gi, we are unable to observe possible differences in conformations stabilized by PZM21 and FH210.

Kelly et al. proposed a pharmacophore model in which strong Y326^7.43^ and D147^3.32^ interaction will increase arrestin recruitment but stronger Y148^3.33^ and weaker D147^3.32^ engagement will lower arrestin interaction. In agreement with their model, FH210’s strong G protein bias could be explained by its weaker polar contacts with Y326^7.43^ and D147^3.32^ compared with PZM21 and DAMGO (Figure S5A). During our MD simulations, we also observed that FH210 spent more time engaged in a water-mediated hydrogen bond to Y148^3.33^ and less time forming a hydrogen bond to Y326^7.43^, in comparison to PZM21 (Figure S5B, C). However, this model cannot explain the biased behavior of PZM21, as its interaction with Y326^7.43^ and D147^3.32^ are very similar to unbiased full agonists DAMGO (Figure S5A). It is worth noting that the dynamics observed in Cong et al. and Kelly et al. are based on the inherent dynamics of agonist bound μOR in the absence of G protein, while in our study the conformation of μOR and the binding pose of the ligands are further affected by the stable coupling to a nucleotide-free G protein.

In summary, this study provides a structure-based evolution strategy for μOR drug discovery. Based on the cryoEM structure of PZM21 bound μOR [18], we have explored the chemical space of PZM21 analogs and identified agonists with improved selectivity and functionality. These novel ligands can be potential therapeutic leads with attenuated side effects. Moreover, we have also determined the cryoEM structure of μOR in complex with FH210 [18], a newly developed naphthyl-substituted acryl amide analog of PZM21. The cryoEM structures and the pharmacological data, suggest that G protein biased signaling of μOR can be achieved by targeting an extended pocket formed by TM2, TM3 and ECL1. These results, together with other recent advances in the field, provide valuable molecular templates for the design of safer analgesics and a deeper understanding of biased agonism at this important pharmaceutical target.

## Acknowledgements

This work was supported by the American Heart Association Postdoctoral Fellowship (H.W.), the National Institutes of Health grants R37DA036246 (B.K.K), T32GM089626 (J.G.M.), and the DFG grant GRK 1910 (F.H., J.K., H.H., P.G.). B.K.K. is a Chan Zuckerberg Biohub Investigator. CryoEM data collection was performed at the Stanford CryoEM Center (cEMc).. The authors thank Elizabeth Montabana and Chensong Zhang for their support on CryoEM data collection. CryoEM data processing for this project was performed on the Sherlock cluster from the Stanford Research Computing Center.

## Competing interests

B.K.K. and P.G. are co-founders of Epiodyne. B.K.K is a co-founder of and consultant for ConformetRx.

