## Supporting Information for "Structure-based Evolution of G protein-biased μ-opioid Receptor Agonists"

### Table of Contents

***Supporting Figures*** **2**

***Supporting Tables*** **7**

***Material and Methods*** **16**

***Model building and refinement*** **17**

***Supporting Scheme*** **22**

***Chemical synthesis material and methods*** **23**

***References*** **45**

***$^{1}$H-NMR and $^{13}$C-NMR spectra*** **48**

**Supporting Figures**

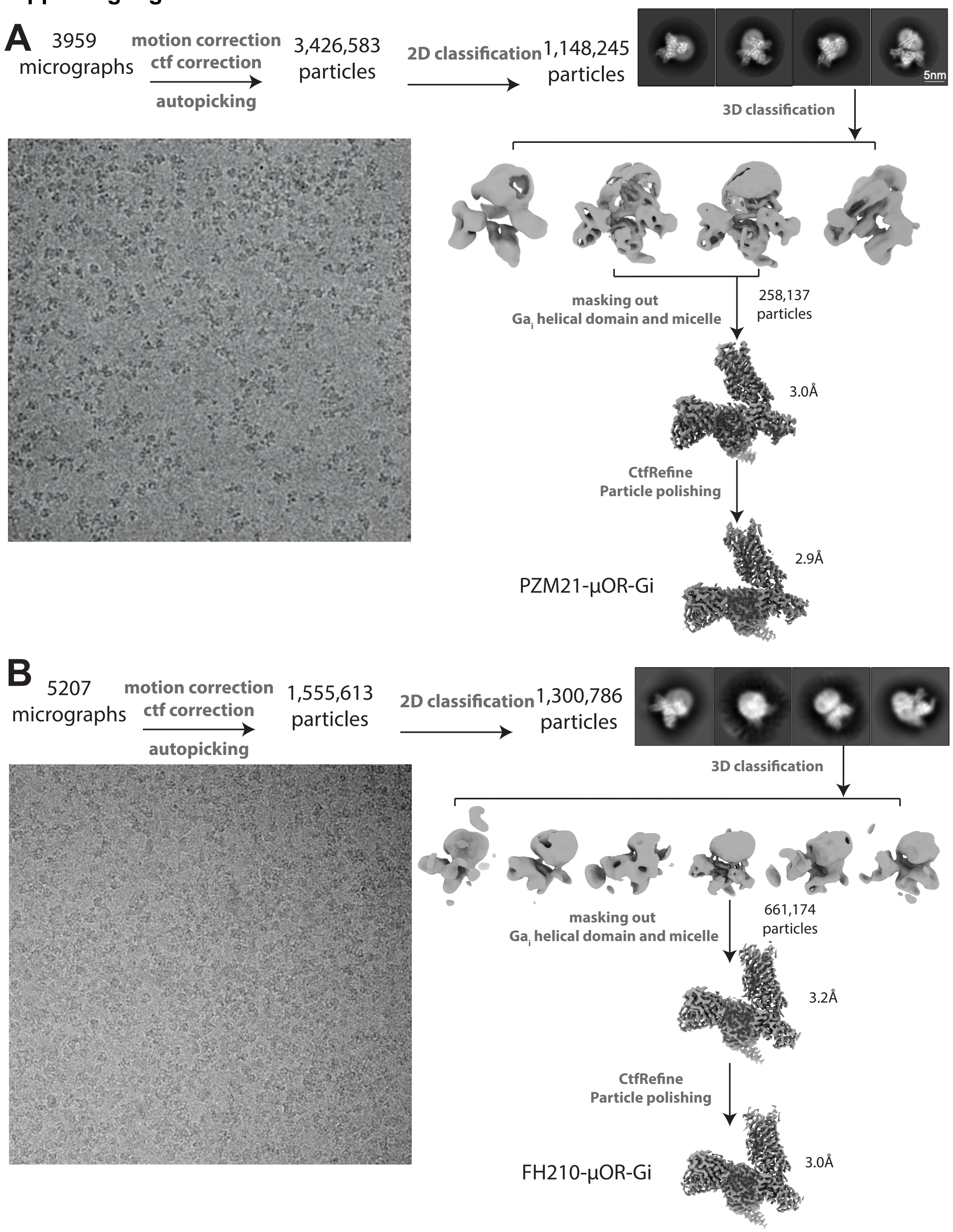


**Figure S1.** Work-flows of cryo-EM data processing procedures for PZM21 (**A**) and FH210 (**B**) bound μOR–G$_i$-scFv16 complex.

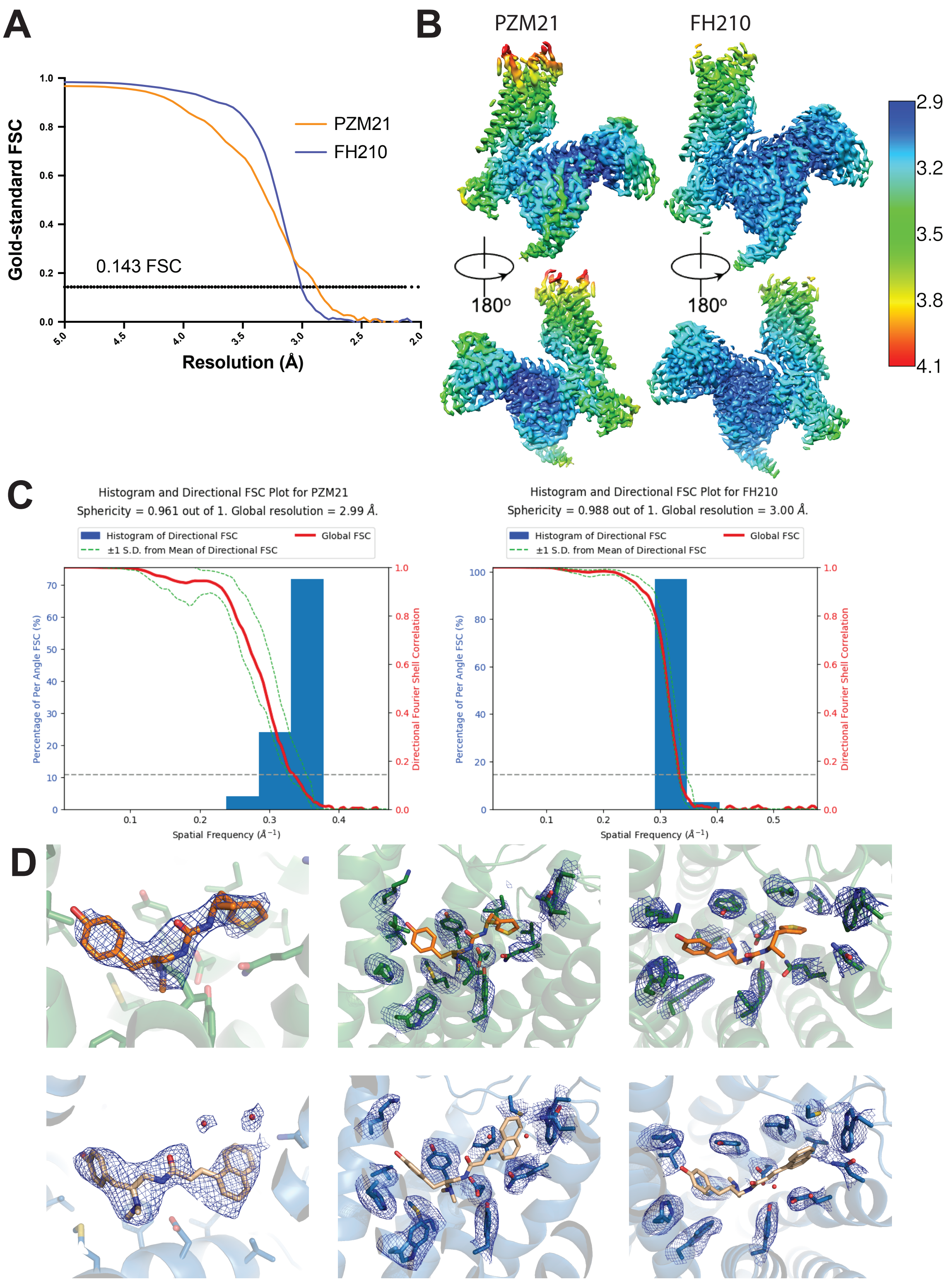


**Figure S2.** Resolution assessment of cryo-EM maps. Gold-standard FSC curves (**A**), local resolution (**B**), and 3DFSC plots (**C**) for PZM21 and FH210 bound μOR–$G_i$-scFv16 structures. Overall resolution is 2.9 Å for PZM21 bound μOR–$G_i$-scFv16 and 3.0 Å for FH210 bound μOR–$G_i$-scFv16 using the gold Standard FSC = 0.143 criterion. **D**) Zoom in view of the ligands and orthosteric site residues densities.

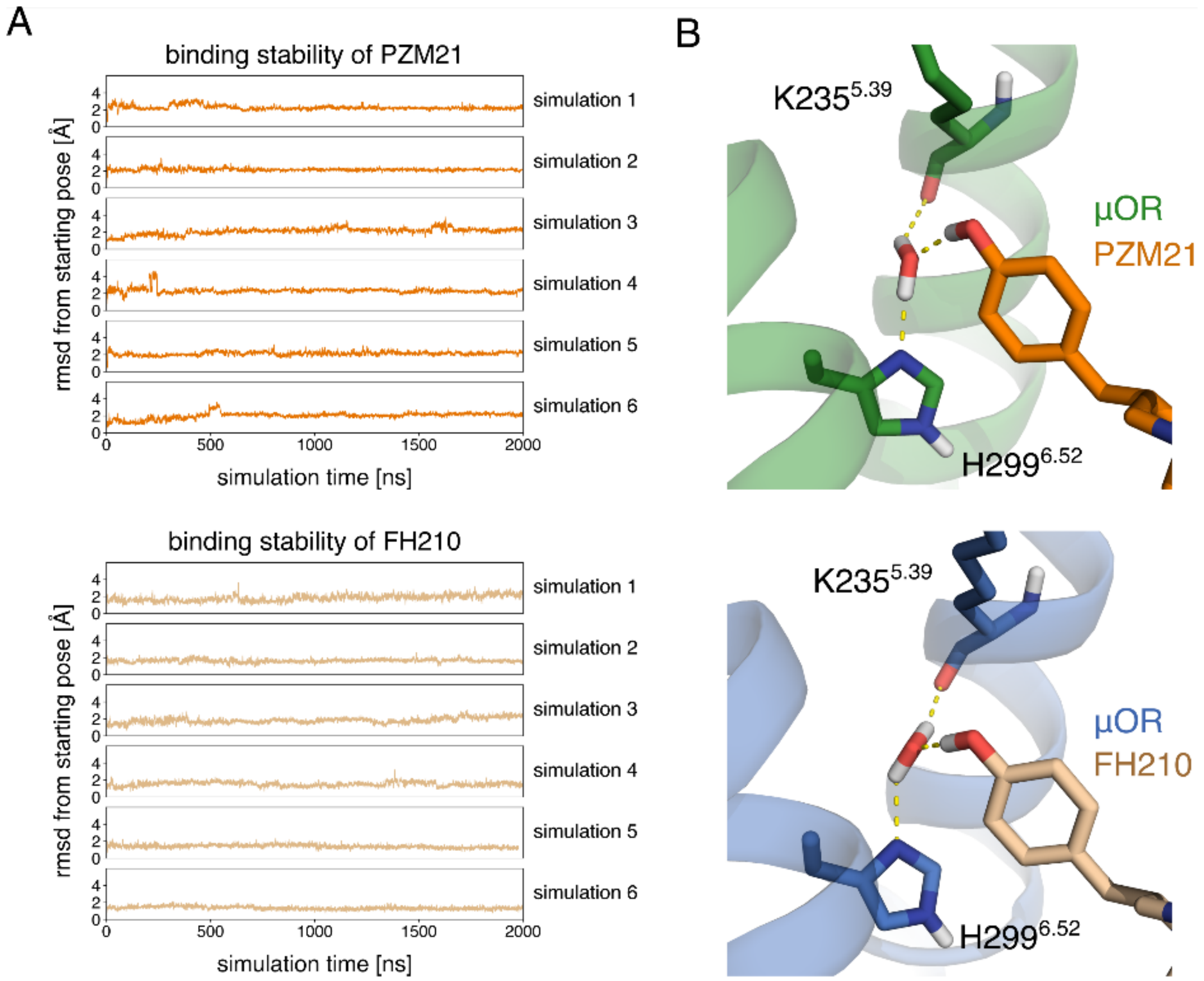


**Figure S3:** Stability of ligand binding to μOR in MD simulations and presence of water mediated interaction of the phenol groups of PZM21 and FH210. (**A**) The root-mean-square deviation (rmsd) for PZM21 (orange) and FH210 (brown), relative to the starting conformation, is displayed. The rmsd was calculated for all non-hydrogen atoms. PZM21 and FH210 remain close to the initial starting structure during productive MD simulations. (**B**) A representative frame of the two simulation conditions is displayed. For the two systems, a water mediated interaction is observed between the phenol of PZM21 or FH210 and L235$^{5.39}$ and H299$^{6.52}$.

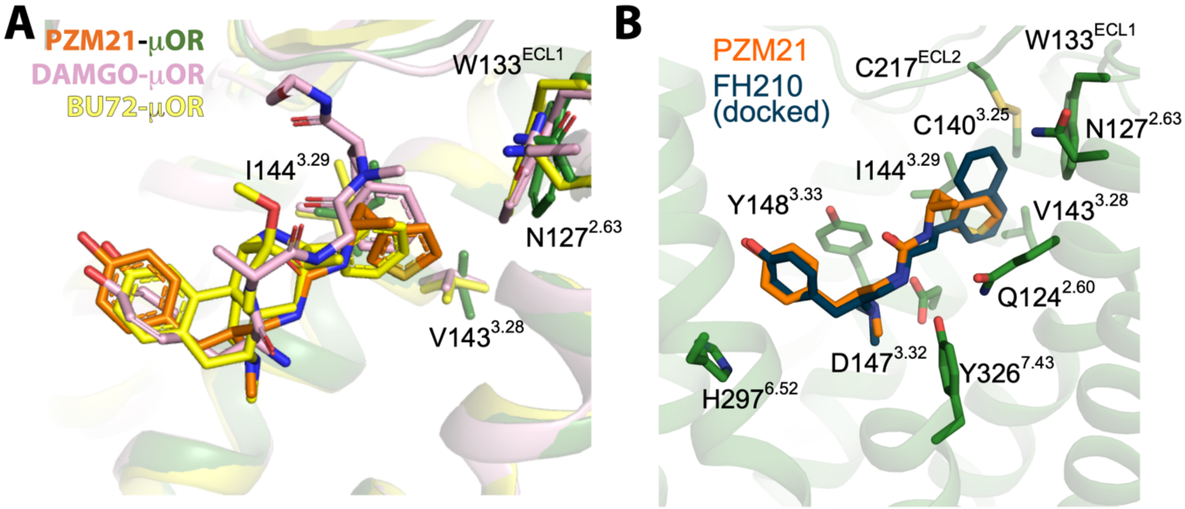


**Figure S4**: (**A**) Overlap of PZM21 (orange), DAMGO (pink), and BU72 (yellow) binding poses. The thiophenylalkyl moiety of PZM21 extend further into the extracellular vestibule compared with DAMGO and BU72. (**B**) Overlap of FH210 (dark blue) docked into the orthosteric pocket of the μOR–$G_i$ complex with PZM21 (orange). The docking pose of FH210 shows a comparable position of the phenol-group, the ammonium cation and the amide nitrogen.

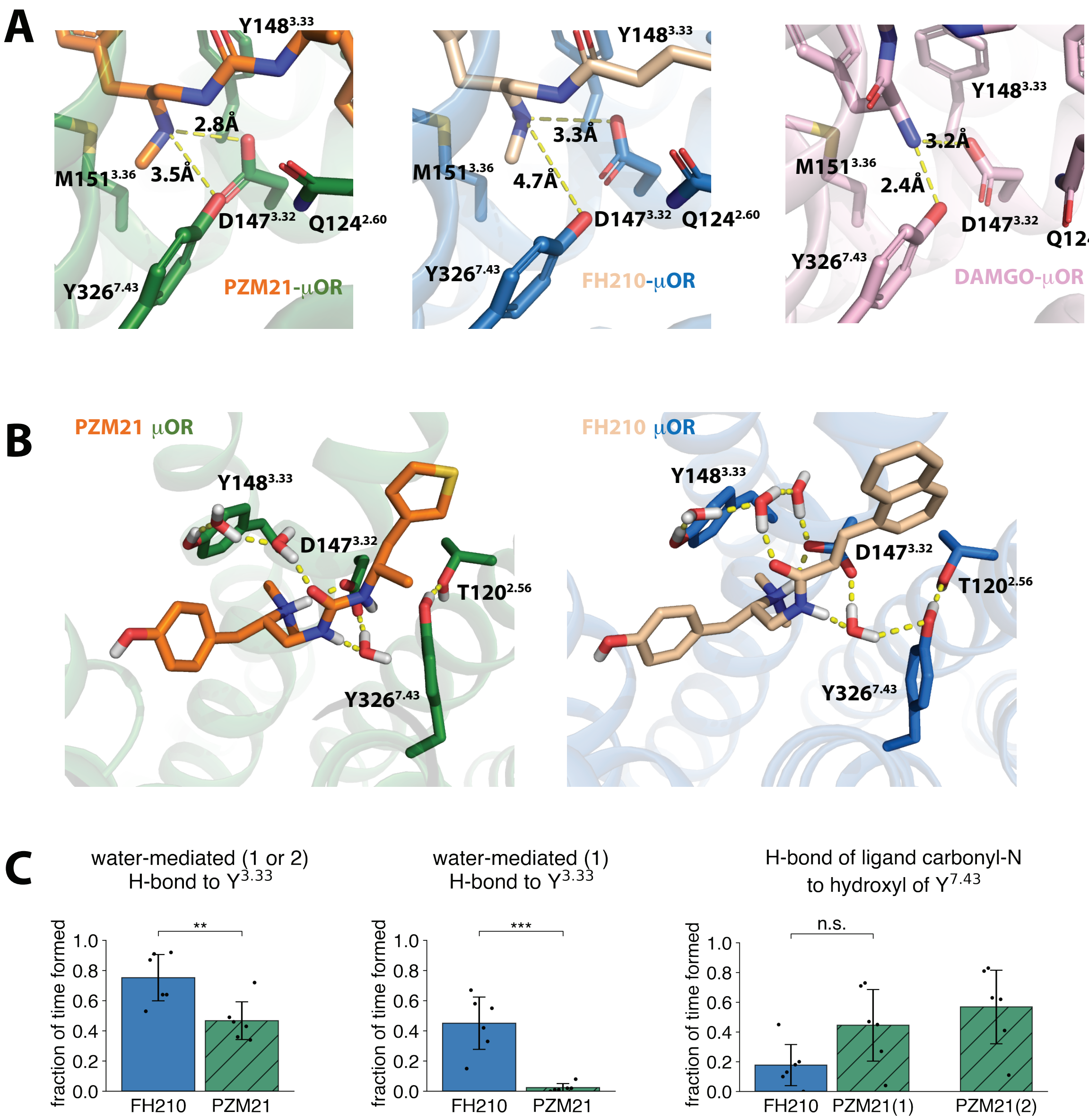


**Figure S5:** A) The polar contacts between PZM21, FH210, and DAMGO and D147$^{3.32}$ and Y326$^{7.43}$. B) The representative structures from MD simulations of PZM21-μOR and FH210-μOR. Water-mediated H-bonds to D147$^{3.32}$, Y148$^{3.33}$, and Y326$^{7.43}$ are shown in yellow dash lines. C) The fraction of time of FH210/PZM21 engaged in (water-mediated) hydrogen bonds with Y148$^{3.33}$ and Y326$^{7.43}$. The bar charts show the mean and SEM, with the dots showing the individual values. The statistics are based on 6 individual simulations. The first 500ns of every simulation were not included in these analyses.

**Supporting Tables**
**Table S1.** Data collection, refinement, and model statistics.

| | **PZM21-μOR-Gi-scFv16** | **FH210-μOR-Gi-scFv16** |
|---|---|---|
| **EM Data collection and processing** | | |
| Voltage (kV) | 300 | 300 |
| Electron exposure (e-/Å$^2$) | 66.75 | 68.5 |
| Pixel size (Å) | 1.06 | 0.8676 |
| Symmetry imposed | C1 | C1 |
| Final particle images (no.) | 258,137 | 661,174 |
| Map resolution(Å) | 2.9 | 3 |
| final b_iso obtained (Å2) | 100.58 | 104.5 |
| Initial model used | PDB: 6DDE | PDB: 6DDE |
| **Model vs. Data** | | |
| CC (mask) | 0.82 | 0.88 |
| FSC (model) = 0/0.143/0.5 (Å) | 2.8/2.9/3.0 | 2.9/3.0/3.1 |
| **Composition** | | |
| Non-hydrogen atoms | 8529 | 8539 |
| Protein residues | 1119 | 1121 |
| Water | 0 | 2 |
| Ligands | 1 | 1 |
| **Bonds (RMSD)** | | |
| Bond lengths (Å) | 0.004 | 0.004 |
| Bond angles (˚) | 0.654 | 0.615 |
| **Validation** | | |
| MolProbity | 1.7 | 1.80 |
| EMRinger score | 3.12 | 4.53 |
| Clashscore | 8.22 | 9.34 |
| rotamer outlier (%) | 0.11 | 0 |
| Cb outliers (%) | 0 | 0 |
| **Ramachandran plot** | | |
| Favored (%) | 96.28 | 95.57 |
| Allowed (%) | 3.54 | 4.43 |
| Outliers (%) | 0.18 | 0 |
| **Deposit** | | |
| EMDB | 24978 | 25034 |
| PDB | 7SBF | 7SCG |

**Table S2**: Molecular docking experiments performed with novel PZM21 analogs at the PZM21-cryo-EM structure using AutoDock Vina[1]. In each experiment, ten docking poses were obtained and filtered for those forming the canonical salt bridge with D149$^{3.32}$, a hydrogen bond to Y328$^{7.43}$ and an adequate superimposition with the native PZM21 binding pose. Among the ones matching these criteria, the top-scoring docking pose is reported below.

| Cmpd. | Structure | Docking Pose | Score [kcal/mol] | logP a) |
|---|---|---|---|---|
| PZM21 | 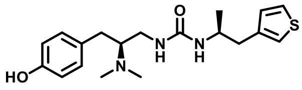 | 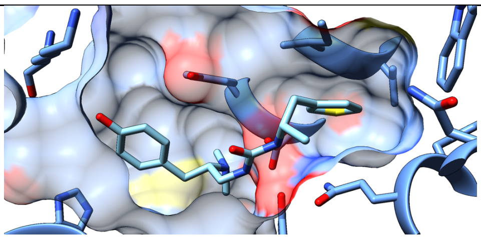 (PZM21-cryo-EM complex) | - | 2.9 |
| FH310 | 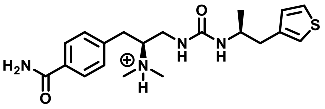 | 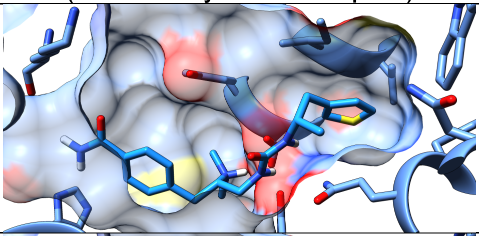 | -8.6 | 2.3 |
| FH321 | 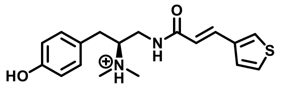 | 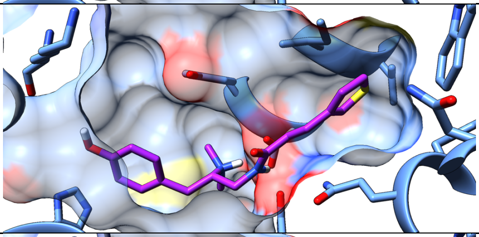 | -6.8 | 2.8 |
| FH163 | 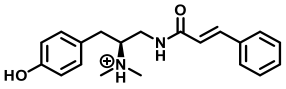 | 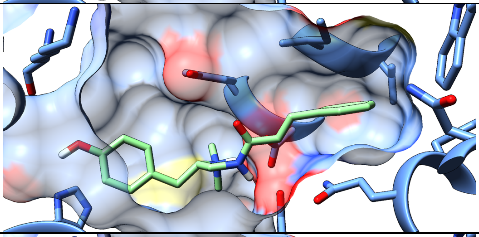 | -8.3 | 2.7 |
| FH172 | 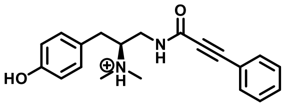 | 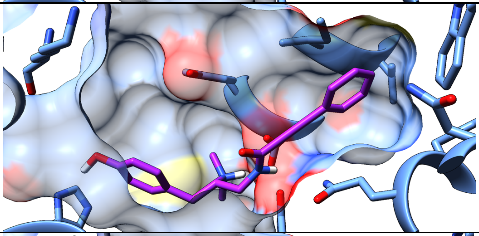 | -8.1 | 2.0 |
| FH210 | 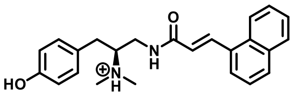 | 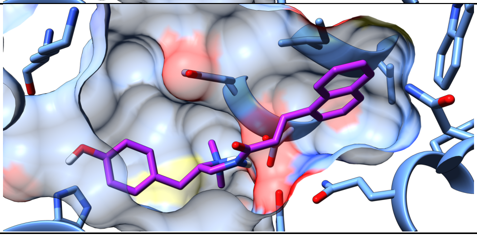 | -9.5 | 3.9 |

| | | | | |
|---|---|---|---|---|
| FH217 | 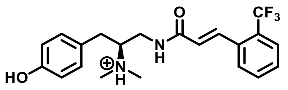 | 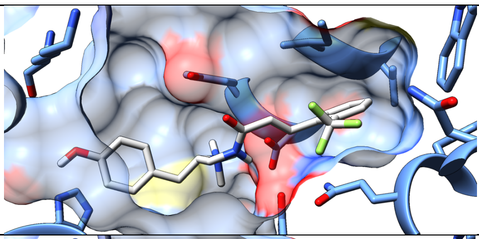 | -9.0 | 3.7 |
| FH218 | 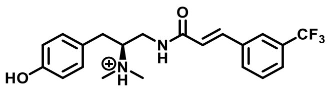 | 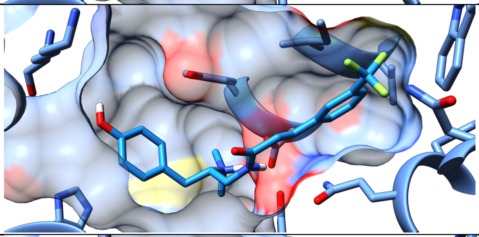 | -9.2 | 3.7 |
| FH219 | 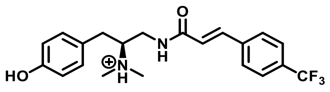 | 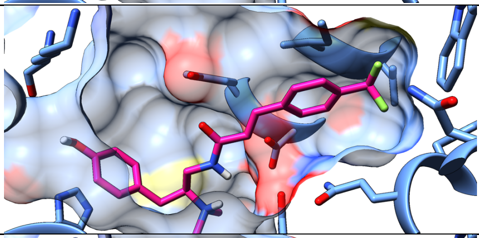 | -8.5 | 3.7 |
| FH299 | 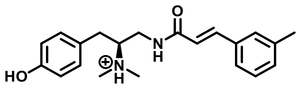 | 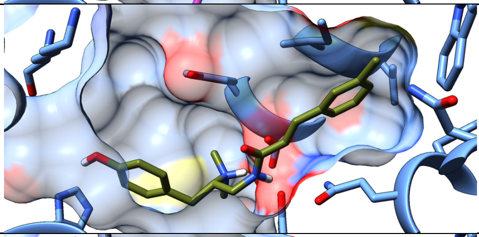 | -8.9 | 3.0 |
| FH315 | 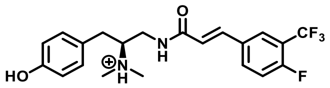 | 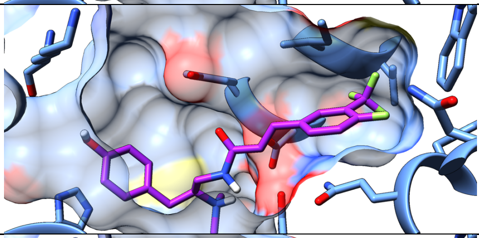 | -9.0 | 3.9 |
| FH_D112 | 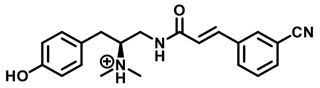 | 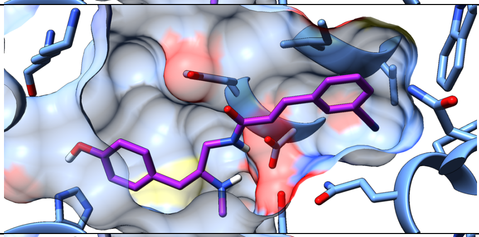 | -8.3 | 2.6 |
| FH_D113 | 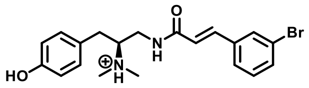 | 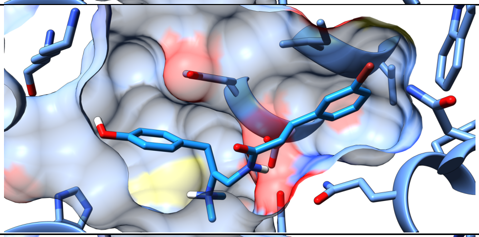 | -8.2 | 3.6 |
| FH_D114 | 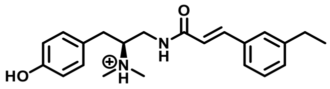 | 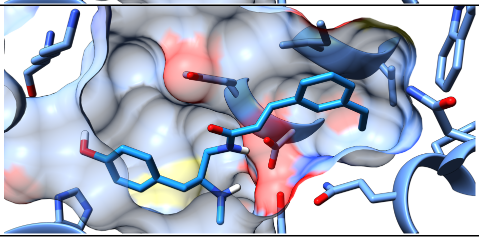 | -8.2 | 3.3 |

| | | | | |
|---|---|---|---|---|
| FH223 | 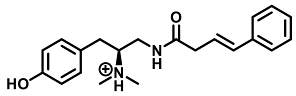 | 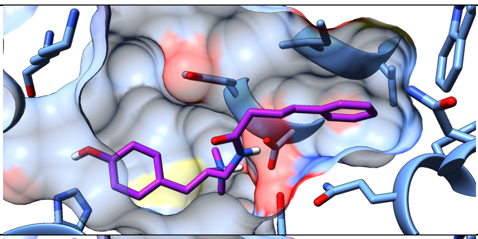 | -8.2 | 3.1 |
| FH273 | 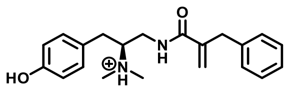 | 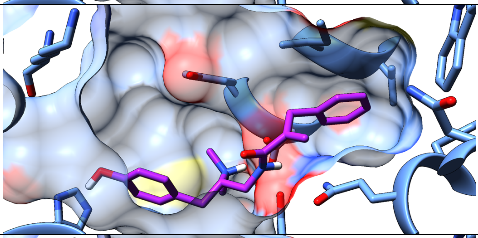 | -7.7 | 2.8 |
| FH_D74 | 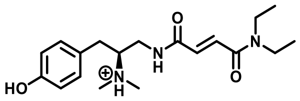 | 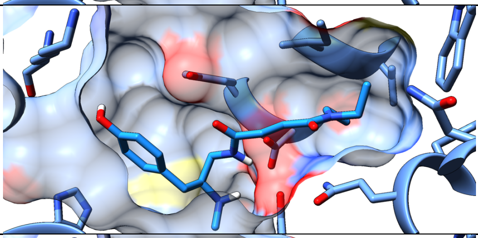 | -6.9 | 1.4 |
| FH_D102 | 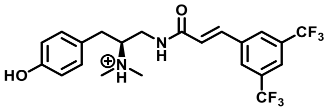 | 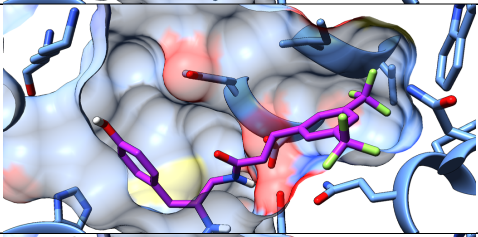 | -9.8 | 4.7 |
| FH_D103 | 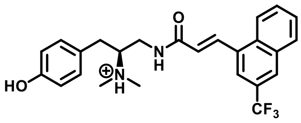 | 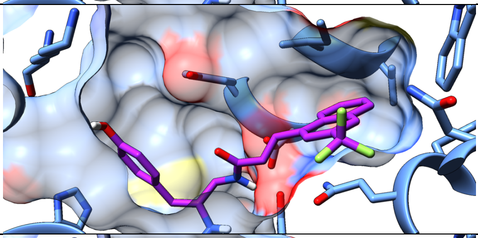 | -10.0 | 4.9 |
| FH_D104 | 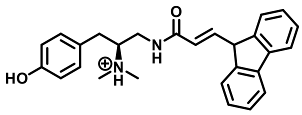 | 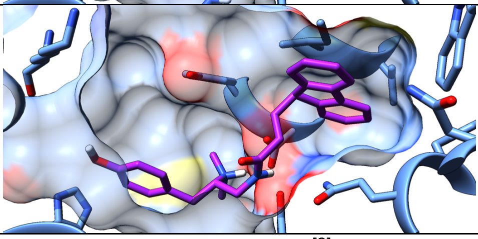 | -10.7 | 4.4 |

a) logP calculated for the neutral species; method by Crippen et al.[2]

**Table S3**: Binding affinities and subtype selectivity of novel PZM21 analogs derived from radioligand binding experiments with human δOR, κOR and μOR subtypes[a]

| compound | structure | $K_i$ value in [nM ± SEM][b] | | | ratio[c] of | |
|---|---|---|---|---|---|---|
| | | δOR | κOR | μOR | δ/μOR | κ/μOR |

| | | | | | | |
|---|---|---|---|---|---|---|
| morphine | 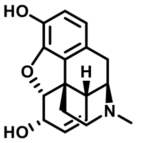 | 1500 ± 620 | 1200 ± 310 | 49 ± 5 | 31 | 24 |
| oliceridine | 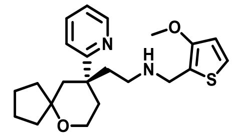 | 1400 ± 270 | 2700 ± 300 | 24 ± 4 | 58 | 112 |
| fentanyl | 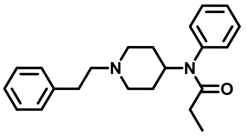 | 210 ± 56 | 510 ± 51 | 11 ± 2 | 19 | 46 |
| PZM21 | 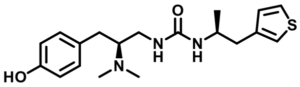 | 540 ± 96 | 24 ± 3 | 31 ± 4 | 17 | 0.77 |
| FH310 | 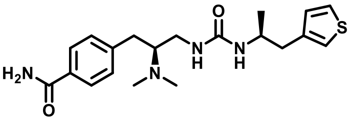 | 160 ± 22 | 58 ± 15 | 6.6 ± 1 | 24 | 8.8 |
| FH321 | 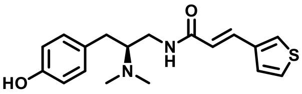 | 59 ± 8 | 130 ± 15 | 8.5 ± 1 | 6.9 | 15 |
| FH163 | 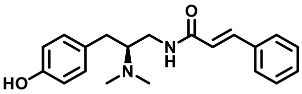 | 25 ± 2 | 120 ±14 | 13 ± 2 | 1.9 | 9.2 |
| FH172 | 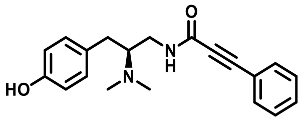 | 220 ± 26 | 400 ± 42 | 22 ± 2 | 7.9 | 18 |
| FH210 | 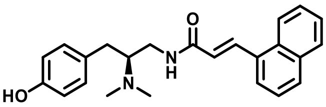 | 220 ± 30 | 190 ± 32 | 18 ± 1 | 12 | 11 |
| FH217 | 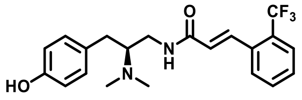 | 180 ± 21 | 140 ± 6 | 28 ± 2 | 6.4 | 5.0 |
| FH218 | 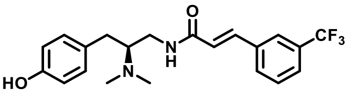 | 120 ± 10 | 590 ± 54 | 30 ± 5 | 4.0 | 20 |
| FH299 | 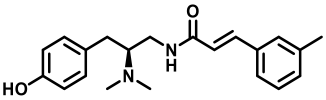 | 300 ± 40 | 500 ± 32 | 24 ± 3 | 13 | 21 |
| FH315 | 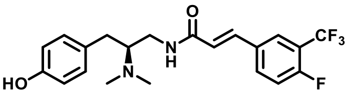 | 250 ± 19 | 600 ± 71 | 23 ± 3 | 11 | 26 |
| FH223 | 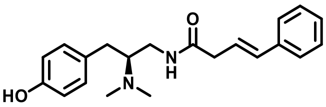 | 530 ± 130 | 390 ± 37 | 15 ± 4 | 35 | 26 |
| FH273 | 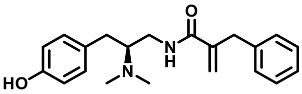 | 250 ± 23 | 210 ± 30 | 41 ± 7 | 6.1 | 5.1 |

[a] Determined by competition binding with membranes from HEK293T cells transiently expressing the human μOR, δOR, and κOR, respectively and the radioligand [$^{3}$H]dynorphine. [b] $K_i$ values are the means ±SEM of 4-15 individual experiments each done in triplicate. [c] Subtype selectivity expressed as ratio of the $K_i$ values of δOR or κOR divided by the $K_i$ value of μOR.

**Table S4:** μOR stimulated G protein signaling and β-arrestin-2 recruitment of PZM21 analogs and references.

| compound | G protein signaling[a] | | β-arrestin-2[b] | | β-arrestin-2 + GRK2[c] | |
|---|---|---|---|---|---|---|
| | $E_{max}$ [%][d] | $EC_{50}$ [nM][e] | $E_{max}$ [%][f] | $EC_{50}$ [nM][e] | $E_{max}$ [%][g] | $EC_{50}$ [nM][e] |

| | | | | | | |
|---|---|---|---|---|---|---|
| fentanyl | 100 | 2.6 ± 0.2 | 100 | 410 ± 72 | 100 | 55 ± 6.7 |
| morphine | 103 ± 1 | 21 ± 3 | 19 ± 1 | 2600 ± 1300 | 80 ± 3 | 390 ± 17 |
| oliceridine | 95 ± 3 | 19 ± 2 | < 5 | no curve | 42 ± 2 | 99 ± 12 |
| PZM21 | 94 ± 3 | 15 ±3 | < 5 | no curve | 35 ± 4 | 94 ± 17 |
| FH310 | 101 ± 1 | 3.3 ± 0.6 | 15 ± 2 | 87 ± 43 | 76 ± 3 | 44 ± 4 |
| FH321 | 55 ± 4 | 7.9 ± 1.3 | < 5 | no curve | < 5 | no curve |
| FH163 | 65 ± 3 | 7.6 ± 1.1 | < 5 | no curve | < 5 | no curve |
| FH172 | 44 ± 8 | 24 ± 3 | < 5 | no curve | < 5 | no curve |
| FH210 | 92 ± 4 | 18 ± 2 | < 5 | no curve | 28 ± 2 | 860 ± 340 |
| FH217 | 45 ± 4 | 14 ± 2 | < 5 | no curve | < 5 | no curve |
| FH218 | 104 ± 2 | 17 ± 3 | < 5 | no curve | 54 ± 2 | 350 ± 50 |
| FH299 | 78 ± 2 | 45 ± 13 | < 5 | no curve | 13 ± 1 | 220 ± 25 |
| FH315 | 104 ± 5 | 39 ± 5 | < 5 | no curve | 48 ±1 | 740 ± 97 |
| FH223 | < 5 | no curve | < 5 | no curve | < 5 | no curve |
| FH273 | 21 ± 3 | 82 ± 12 | < 5 | no curve | < 5 | no curve |

[a] Measurement of G-protein signaling was performed applying the IP-One assay® (Cisbio) in HEK293T cells transiently co-transfected with the human µOR and the hybrid G protein $G\alpha_{qi}$. [b] β-Arrestin-2 recruitment was determined with the PathHunter assay (DiscoverX) in HEK293T cells stably expressing an enzyme acceptor (EA) tagged β-arrestin-2 and the ProLink-PK1 tagged µOR. [c] β-Arrestin-2 recruitment in presence of co-transfected G protein-coupled receptor kinase 2. [d] Efficacy of G protein signaling [% ± SEM] relative to the maximum effect of fentanyl derived from three to twelve individual experiments each done in duplicates.[e] Potency displayed in nM ± SEM. [f] Efficacy of arrestin recruitment [% ± SEM] relative to fentanyl was derived from three to ten individual experiments each done in duplicates. [g] Efficacy of arrestin recruitment in presence of GRK2 [% ± SEM] relative to fentanyl derived from three to eight repeats each done in duplicates. No curve: no $EC_{50}$ value could be determined due to low efficacy less than 5 %.

**Table S5**: Selectivity profile of binding affinity for PZM21, FH210, FH218 and FH310 to a set of class A GPCRs.[a]

| GPCR | radioligand | $K_i$ [nM ± SD][b] | | | |
|---|---|---|---|---|---|
| | | PZM21 | FH210 | FH218 | FH310 |

| | | | | | |
|---|---|---|---|---|---|
| adrenoceptor | | | | | |
| $\alpha_{1A}$ | [$^3$H]prazosine | 6000 ± 2700 | 490 ± 99 | 740 ± 110 | 41000 ± 7800 |
| $\alpha_{2A}$ | [$^3$H]RX821002 | 47000 ± 27000 | 1800 ± 420 | 4800 ± 1600 | >50000 |
| $\alpha_{2B}$ | [$^3$H]RX821002 | 7600 ± 1300 | 610 ± 110 | 1700 ± 350 | 11000 ± 4000 |
| $\beta_1$ | [$^3$H]CGP12177 | >50000 | 2300 ± 780 | 20000 ± 2100 | >50000 |
| $\beta_2$ | [$^3$H]CGP12177 | >50000 | 6700 ± 2300 | 12000 ± 2100 | >50000 |
| dopamine | | | | | |
| $D_1$ | [$^3$H]SCH23390 | 32000 ± 6300 | 1200 ± 280 | 1200 ± 320 | >50000 |
| $D_{2long}$[c] | [$^3$H]spiperone | 25000 ± 3500 | 2200 ± 490 | 4000 ± 1100 | >50000 |
| $D_3$[d] | [$^3$H]spiperone | >50000 | 6000 ± 0 | 4100 ± 990 | >50000 |
| $D_{4.4}$[e] | [$^3$H]spiperone | 20000 ± 4100 | 940 ± 78 | 1200 ± 190 | >50000 |
| $D_5$ | [$^3$H]SCH23390 | >50000 | 1700 ± 780 | 1100 ± 400 | >50000 |
| serotonin | | | | | |
| 5-$HT_{1A}$ | [$^3$H]WAY100635 | >100000 | 10000 ± 3600 | 23000 ± 1400 | >50000 |
| 5-$HT_{2A}$ | [$^3$H]ketanserin | 580 ± 35 | 190 ± 160 | 370 ± 8 | 47000 ± 21000 |
| 5-$HT_6$ | [$^3$H]N-methyl-LSD | 28000 ± 17000 | 8800 ± 3200 | 17000 ± 6400 | >50000 |
| muscarinic | | | | | |
| $M_1$ | [$^3$H]NMS | 3400 ± 140 | 11000 ± 7400 | 8400 ± 2100 | >50000 |
| $M_2$ | [$^3$H]NMS | 4400 ± 1600 | 6500 ± 1300 | 29000 ± 5700 | >50000 |
| $M_3$ | [$^3$H]NMS | 3900 ± 1600 | 15000 ± 14000 | 27000 ± 710 | >50000 |
| neurotensin | | | | | |
| $NTS_1$ | [$^3$H]NT(8-13) | >50000 | >50000 | >50000 | >50000 |
| $NTS_2$ | [$^3$H]NT(8-13) | >50000 | >50000 | 45000 ± 22000 | >50000 |
| orexin | | | | | |
| $OX_1$ | [$^3$H]SB674042 | >50000 | >50000 | >50000 | >50000 |
| $OX_2$ | [$^3$H]EMPA | >50000 | >50000 | >50000 | >50000 |

[a] Radioligand displacement experiments with membranes from HEK 293T cells transiently transfected with the appropriate receptor. [b] Binding affinity displayed as mean $K_i$ value in nM ± SD from two to four individual experiments each done in triplicate. [c] Dopamine $D_{2long}$ receptor was stably expressed in CHO cells. [d] Dopamine $D_3$ receptor was stably expressed in dhfr$^-$ CHO cells. [e] The receptor isoform $D_{4.4}$ was stably expressed in CHO cells.

**Table S6**: G protein subtypes profile of PZM21, FH210, FH218 and FH310 measured by BRET TRUPATH.

| compound | Gi1 | | Gi2 | | Gi3 | | GoA | | GoB | | Gz | |
|---|---|---|---|---|---|---|---|---|---|---|---|---|
| | $E_{max}$[%] | $EC_{50}$[nM] | $E_{max}$[%] | $EC_{50}$[nM] | $E_{max}$[%] | $EC_{50}$[nM] | $E_{max}$[%] | $EC_{50}$[nM] | $E_{max}$[%] | $EC_{50}$[nM] | $E_{max}$[%] | $EC_{50}$[nM] |
| morphine | 104±4 | 17±4 | 101±4 | 14±1 | 101±1 | 20±4 | 99±0 | 5.4±1.2 | 105±3 | 6±0.7 | 101±2 | 3.9±0.6 |
| fentanyl | 110±3 | 11±2 | 98±4 | 6.3±1.2 | 115±6 | 14±1 | 105±10 | 8.7±2.9 | 107±4 | 6.8±1.9 | 91±1 | 2.9±1.4 |
| PZM21 | 68±3 | 10±2 | 73±13 | 9.2±1.7 | 68±7 | 14±1 | 80±7 | 11±1 | 89±5 | 12±3 | 77±3 | 3.7±1 |
| FH210 | 72±2 | 14±1 | 80±10 | 10±1 | 67±7 | 19±6 | 73±4 | 8.5±3.6 | 85±7 | 7.3±1.7 | 80±3 | 3.8±1.3 |

Efficacy of [% ± SEM] relative to the maximum effect of morphine derived from three individual experiments each done in duplicates. Potency displayed in nM ± SEM, derived from three individual experiments each done in duplicates.

### Material and Methods

#### Expression and purification of μOR

For the PZM21-bound μOR complex, we used a modified *M. musculus* μOR construct with removable N-terminal Flag-tag and C-terminal histidine tag. N-terminal residues (1-63) of μOR were replaced with the thermostabilized apocytochrome $b_{562}$RIL from Escherichia coli (M7W, H102I and R106L) (BRIL) protein and a linker of GSPGARSAS. N-terminal Flag-tag and C-terminal histidine tag were removable with rhinovirus 3C protease. For the FH210-bound μOR complex, we used a wild type full-length *M. musculus* μOR construct with N-terminal hemagglutinin (HA) signal sequence and FLAG tag, and C-terminal histidine tag.

Basically, μOR constructs were expressed in *Spodoptera frugiperda* Sf9 insect cells using the baculovirus method (Expression Systems). The receptors were solubilized from cell membranes with 1% *n*-dodecyl-β-D-maltoside (DDM, Anatrace)/0.1% cholesterol hemisuccinate (CHS), and purified by nickel-chelating sepharose chromatography. The Ni-NTA eluate was then incubated with 1% lauryl maltose neopentyl glycol (L-MNG)/0.1% CHS for 1 hour on ice. After detergent exchange, 2mM $CaCl_2$ was added and the sample was loaded onto M1 anti-Flag resin and washed with progressively lower concentrations of salt. The μOR was then eluted from M1 resin in a buffer consisting of 20 mM Hepes pH 7.5, 100 mM NaCl, 0.003% L-MNG/0.0003% CHS supplemented with 1 uM naloxone, Flag peptide and 5 mM EDTA. The M1 elute was further purified by size exclusion chromatography on the Superdex 200 10/300 gel filtration column (GE Healthcare) in 20 mM HEPES pH 7.5, 100 mM NaCl, 0.001% L-MNG/0.0001% CHS. The monomeric fractions were pooled, concentrated, and flash frozen in liquid nitrogen.

#### Expression and purification of heterotrimeric $G_i$

Heterotrimeric $G_i$ was expressed and purified as previously described[3]. Basically, *Trichuplusia ni* Hi5 insect cells were co-infected with two viruses, one encoding the wild-type human Gαi subunit and another encoding the wild-type human β1γ2 subunits with an histidine tag inserted at the N-terminus of the β1 subunit. After 48 hours, cells were harvested and lysed in hypotonic buffer. The heterotrimeric Gi was extracted in a buffer containing 1% sodium cholate and 0.05% DDM. The soluble fraction was purified using Ni-NTA chromatography, and the detergent was exchanged from cholate/DDM to DDM on column. After elution, human rhinovirus 3C protease (3C protease) was added and the histidine tag was cleaved overnight at 4°C during dialysis. Then the heterotrimeric Gi without tag will be further purified through reverse Ni-NTA chromatography. Finally, the flow through of the reverse Ni-NTA step will be purified by the Mono Q 5/50 GL column (GE Healthcare) to get rid of the 3C protease. The purified heterotrimeric $G_i$ will be concentrated, stored in 20 mM HEPES pH 7.5, 100 mM NaCl, 0.05% DDM, 100 uM TCEP, 10 uM GDP before complexing.

#### Expression and Purification of scFv16

scFv16 was developed and purified as previously described[4]. Basically, scFv with C terminal His tag was expressed in Trichuplusia ni Hi5 insect cells. After infection and expression, the insect cell supernatant was loaded onto Ni-NTA resin and the scFv was eluted in 20 mM HEPES pH 7.5, 500 mM NaCl, and 250 mM imidazole. The eluate was incubated with 3C protease overnight to cleave the C-terminal His tag. After dialysis into the buffer consisting of 20mM HEPES pH 7.5 and 100 mM NaCl, scFv16 was further purified by reverse Ni-NTA chromatography. The flow-through was collected and applied over a Superdex 200 16/60 column (GE Healthcare). The scFv16 fractions were pooled, concentrated, and flash frozen.

#### Formation and purification of the μOR–$G_i$-scFv16 complex.

500uM ligands (PZM21, FH210) was added to purified μOR while 1% L-MNG was added to purified $G_i$. Both mixtures were incubated on ice for 1 h. After that, ligand-bound μOR was mixed with a 1.5 molar excess of $G_i$ heterotrimer and extra TCEP was added to maintain 100uM TCEP concentration. The coupling reaction was allowed to proceed for another 1 h on ice, followed by addition of apyrase to catalyze GDP hydrolysis to obtain nucleotide free complex. After 30min, a 2 molar excess of scFv16 was also added. The reaction mixture was left on ice overnight to allow stable complex formation. After that, the complexing mixture was purified by M1 anti-Flag affinity chromatography and eluted in 20 mM Hepes pH 7.5, 100 mM NaCl, 0.003% L-MNG, 0.001% glyco-diosgenin (GDN), 0.0004% CHS, 10 uM bitopic ligand, 5 mM EDTA and Flag peptide[5]. After elution, 100uM TCEP was added to provide a reducing environment. Finally, the μOR–Gi-scFv16 complex was purified by size exclusion chromatography on a Superdex 200 10/300 gel filtration column in 20 mM Hepes pH 7.5, 100 mM NaCl, 5 uM ligand, 0.003% L-MNG and 0.001% GDN with 0.0004% CHS total. Peak fractions were concentrated to ~10 mg/ml for electron microscopy studies.

**Cryo-EM sample preparation and image acquisition**

For cryo-EM, 3 μL sample was directly applied to glow-discharged 300 mesh gold grids (Quantifoil R1.2/1.3) and vitrified using a FEI Vitrobot Mark IV (Thermo Fisher Scientific). For PZM21 bound μOR–Gi-scFv16 complex, movies were collected on a Titan Krios (SLAC/Stanford) operated at 300 keV using a Gatan K2 Summit direct electron detector in counting mode, with 1.06 Å pixel size. A total of 3959 movies were obtained. Each stack movie was recorded for a total of 10 s with 0.2 s per frame. The dose rate was 1.335 electrons/Å$^2$/subframe, resulting in an accumulated dose of 66.75 electrons per Å$^2$ (Figure S1). For FH210 bound μOR–Gi-scFv16 complex, movies were collected on a Titan Krios (SLAC/Stanford) operated at 300 keV using a Gatan K3 direct electron detector in super resolution mode, with 0.4338 Å pixel size. A total of 5207 movies were obtained. Each stack movie was recorded for a total of 2.5 s with 0.05 s per frame. The dose rate was 1.37 electrons/Å$^2$/subframe, resulting in an accumulated dose of 68.5 electrons per Å$^2$. Both datasets were collected using SerialEM[6].

**Cryo-EM data processing**

Dose-fractionated movies were subjected to beam-induced motion correction using RELION3[7]. For the PZM21 bound μOR–Gi-scFv16 dataset, we used unbinned movie while for the FH210 guano bound μOR–Gi-scFv16 dataset, the movies were binned by 2. CTF parameters for each micrograph were determined by CTFFIND-4.1[8]. Particle autopicking, 2D and 3D classification, and 3D auto-refine were performed in RELION3. Basically, the autopicked particles were frist subjected to 2D classification (Figure S1), 2D classes that look like GPCR-G protein complex were selected for 3D classification (Figure S1). The cryo-EM map of DAMGO bound μOR-Gi-scFv16 complex was low-pass filtered to 60Å and used as reference for 3D classification[9]. After that, the best 3D class were selected and the particles are subjected to 3D auto-refine, during which the G$\alpha_i$ helical domain and micelle were maked out. Further CTF refinement and Bayesian polishing of these particles were performed in RELION3 as well. This generate a 2.9A resolution map for PZM21 bound μOR–Gi-scFv16 complex and a 3.0A resolution map for FH210 guano bound μOR–Gi-scFv16 complex (Figure S1,2). The maps are autosharpened in Phenix[10]. The directional resolution and density isotropy are quantified using 3DFSC[11] (Figure S2C).

**Model building and refinement**

ligand models were generated by eLBOW in Phenix[12]. Models were rigid-body docked into the corresponding cryo-EM density map in Chimera[13], followed by iterative manual adjustment in COOT[14], and real space refinement in Phenix. Ligand coordination were further optimized by GemSpot[15]. The refinement statistics were provided in Table S1.

**Molecular Docking**

Docking studies at the PZM21-OR-Gi complex were carried out utilizing AutoDock Vina version 1.1.2[1]. In order to improve docking performance, a water molecule was manually modeled into the orthosteric site using UCSF Chimera[13], mediating a hydrogen bond of the phenol to Y150$^{3.33}$. The receptor structure was prepared using AutoDock Tools 1.5.6[16]. The three-dimensional ligand structures were prepared with Avogadro 1.1.1[17] and AutoDock Tools 1.5.6. Docking was performed with an exhaustiveness value of 30. Ten ranked ligand binding poses were evaluated based upon comparison to the native PZM21 conformation observed in the cryo-EM complex.

**Unbiased Simulations of Receptor Ligand Complexes**

Simulations of µOR are based on the herein reported structures of the µOR in complex with PZM21 and FH210. Coordinates were prepared by removing the Gβ/γ subunit of the heterotrimeric G protein. As internal water molecules play an important role for the µOR, crystal water resolved in the previously reported active-state crystal structure of the µOR in complex with BU72 (PDB code: 5C1M)[18] were included in the simulation systems. This was achieved by structurally aligning the respective structures and transferring the coordinates of the water molecules. For the simulations of FH210 we only included the water molecules that did not overlap with the already present water molecules.

The Protein Preparation Wizard (Schrödinger Release 2020-4: Protein Preparation Wizard; Epik, Prime, Schrödinger, LLC, New York, NY, 2020.)[19] was used to further prepare the structures. Missing side chains were modeled, hydrogen added, and the protein chain termini capped with the neutral acetyl and methylamide groups. Titratable residues were left in their dominant protonation state at pH 7.0 and PZM21 and FH210 were protonated at the secondary amine. The hydrogen bond networks, including the orientation of water molecules, were optimized. In the following, the structures were energy minimized using the OPLS4 force field in the presence of the Gα subunit. The Gα subunit was removed afterwards.

The protein structures were aligned to the Orientation of Proteins in Membranes (OPM)[20] structure of active µOR (PDB code: 5C1M). Each complex was inserted into a solvated and pre-equilibrated membrane of dioleoyl-phosphatidylcholine (DOPC) lipids using the GROMACS tool g_membed[21]. Subsequently, water molecules were replaced by sodium and chloride ions to give a neutral system with 0.15 M NaCl. The final box dimensions were approximately 80 x 80 x 100 $Å^3$.

Parameter topology and coordinate files were built up using the tleap module of AMBER18[22] and subsequently converted into GROMACS input files. For all simulations, the general AMBER force field (GAFF)[23] was used for PZM21 and FH210, the lipid14 force field[24] for the DOPC molecules and ff14SB[25] for the protein residues. The SPC/E water model[26] was applied. Parameters for PZM21 and FH210 were assigned using antechamber[22]. The structures of PZM21 and FH210 were optimized using Gaussian 16[27] at the B3LYP/6-31G(d) level of theory and charges calculated at HF/6-31G(d) level of theory. Subsequently, atom point charges were assigned according to the RESP procedure[26]. A formal charge of + 1 was assigned to PZM21 and FH210.

Simulations were performed using GROMACS 2018.4[28]. Each simulation system was energy minimized and equilibrated in the NVT ensemble at 310 K for 1 ns followed by the NPT ensemble for 1 ns with harmonic restraints of 10.0 $kcal \cdot mol^{-1}$ on protein and ligands. In the NVT ensemble, the V-rescale thermostat was used. In the NPT ensemble the Berendsen barostat, a surface tension of 22 $dyn \cdot cm^{-1}$, and a compressibility of $4.5 \times 10^{-5}$ $bar^{-1}$ was applied. The system was further equilibrated for 25 ns with restraints on protein backbone and ligand atoms. Here, the restraints were reduced every 5 ns in a stepwise fashion to be 10.0, 5.0, 1.0, 0.5 and 0.1 $kcal \cdot mol^{-1}$, respectively. Productive simulations were performed using periodic boundary conditions and a time step of 2 fs with bonds involving hydrogen constrained using LINCS[29]. Long-range electrostatic interactions were computed using the particle mesh Ewald (PME)[30] method with interpolation of order 4 and fast Fourier transform (FFT) grid spacing of 1.6 Å. Non-bonded interactions were cut off at 12.0 Å.

During production simulations, all residues within 5 Å of the G protein interface were restrained to the initial structure using 5.0 kcal $mol^{-1}$ $Å^{-2}$ harmonic restraints applied to non-hydrogen atoms. Using such restraints instead of the intracellular binding partner reduces the overall system size, enabling faster simulation, while ensuring that the receptor maintains an active conformation throughout the simulation

Analysis of the trajectories was performed using Visual Molecular Dynamics (VMD)[31], CPPTRAJ[32] and GetContacts (https://getcontacts.github.io/). Visualization was performed using the PyMOL Molecular Graphics System, Version 2.1.1 (Schrödinger, LLC). Plots were created using Matplotlib 3.0.2[33].

**Radioligand binding assay**

Binding affinities towards the human μOR, δOR, and κOR were determined as described previously.[34] In brief, membranes were prepared from HEK293T cells transiently transfected with the cDNA for μOR (gift from the Ernest Gallo Clinic and Research Center, UCSF, CA), δOR or κOR (cDNA Resource Center, www.cdna.org) and incubated with the radioligand [$^3$H]diprenorphine (specific activity 31 Ci/mmol; PerkinElmer, Rodgau, Germany) at concentrations of 0.25 nM for μOR, 0.35 nM for δOR, and 0.30 nM for κOR, respectively.[35] Membranes expressing μOR at a $B_{max}$ of 2600 ± 330 fmol/mg protein, a $K_d$ of 0.13 ± 0.03 nM and an amount of protein of 2-10 μg/well, δOR at $B_{max}$= 2200 ± 500 fmol/mg protein, $K_d$= 0.23 ± 0.04 nM and protein= 6-20 μg/well, or κOR at $B_{max}$= 3400 ± 840 fmol/mg protein, $K_d$= 0.11 ± 0.02 nM and protein= 2-6 μg/well, respectively in buffer A (50 mM Tris at pH 7.4) were incubated with radioligand and varying concentrations of test compound (in the range of 1 pM - 100 μM) for 60 min and filtered on glas fiber mats presoaked with 0.3% PEI solution. Trapped radioactivity was measured with a microplate reader (Microbeta Trilux, Perkin Elmer) by scintillation counting. Unspecific binding was determined in the presence of 10 μM of naloxone.

Screening of binding affinities to related GPCRs was performed according to the procedure described above.[34b, 36] Affinities to the dopamine receptors $D_1$ (cDNA from cDNA Resource Center, $B_{max}$= 3000 fmol/mg protein, $K_d$= 0.65 nM, protein= 4 μg/well) and $D_5$ (cDNA Resource Center, $B_{max}$= 3700 fmol/mg protein, $K_d$= 0.55 nM, protein= 3 μg/well) were tested with [$^3$H]SCH23390 (spec. act. 80 Ci/mmol; Biotrend, Cologne, Germany) at 0.4 nM ($D_1$) and 0.5 nM ($D_5$) in buffer B (50 mM Tris, 5 mM $MgCl_2$, 0.1 mM EDTA, 5 μg/mL bacitracin and 5 μg/mL soybean trypsin inhibitor at pH 7.4). Membranes from CHO cells stably expressing the subtype receptors $D_{2long}$ [37] ($B_{max}$= 2000 fmol/mg protein, $K_d$= 0.1 nM, protein= 3 μg/well), $D_3$ [38] ($B_{max}$= 7900 fmol/mg protein, $K_d$= 0.25 nM, protein= 2 μg/well) and $D_{4.4}$ [39] ($B_{max}$= 2100 fmol/mg protein, $K_d$= 0.35 nM, protein= 4 μg/well) were used together with [$^3$H]spiperone (spec. act. 57 Ci/mmol; Biotrend) at concentrations of 0.2 nM ($D_{2long}$), 0.3 nM ($D_3$), and 0.35 nM ($D_{4.4}$), respectively in buffer B.[34b] Unspecific binding to all dopamine receptor subtypes was determined in presence of 10 μM haloperidol. Adrenoceptor binding was determined with membranes from HEK293T cells transiently expressing $\alpha_{1A}$ (cDNA Resource Center, $B_{max}$= 1000 fmol/mg protein, $K_d$= 0.35 nM, protein= 6 μg/well), $\alpha_{2A}$ (gift from Davide Calebiro, University of Birmingham, Birmingham, UK, $B_{max}$= 4000 fmol/mg protein, $K_d$= 3.5 nM, protein= 4 μg/well), $\alpha_{2B}$ (cDNA Resource Center, $B_{max}$= 3500 fmol/mg protein, $K_d$= 1.2 nM, protein= 4 μg/well), $\beta_1$ (gift from Roger Sunahara, UCSD, San Diego, CA, $B_{max}$= 700 fmol/mg protein, $K_d$= 0.07 nM, protein= 10 μg/well), and $\beta_2$ (cDNA Resource Center, $B_{max}$= 1500 fmol/mg protein, $K_d$= 0.12 nM, protein= 8 μg/well) in buffer A ($\alpha_{2A}$, $\alpha_{2B}$), buffer B ($\alpha_{1A}$), or buffer C (25 mM HEPES, 5 mM $MgCl_2$, 1 mM EDTA, 0.006% bovine serum albumin at pH 7.4) ($\beta_1$, $\beta_2$), respectively. [$^3$H]Prazosin (spec. act. 85 Ci/mmol; PerkinElmer) ($\alpha_{1A}$ at 0.3 nM), [$^3$H]RX821002 (spec. act. 52 Ci/mmol; Novandi, Södertälje, Sweden) ($\alpha_{2A}$ at 0.7 nM, $\alpha_{2B}$ at 1.5 nM), and [$^3$H]CGP12177 (spec. act. 52 Ci/mmol; PerkinElmer) ($\beta_1$ at 0.2 nM, $\beta_2$ at 0.3 nM), respectively were used as radioligands. Unspecific binding was determined at 10 μM of the non-labelled radioligand. Affinities to the serotonin receptors 5-$HT_{1A}$ ($B_{max}$= 5000 fmol/mg protein, $K_d$= 0.09 nM, protein= 2 μg/well), 5-$HT_{2A}$ ($B_{max}$= 3000 fmol/mg protein, $K_d$= 0.45 nM, protein= 4 μg/well), and 5-$HT_6$ ($B_{max}$= 2900 fmol/mg protein, $K_d$= 2.1 nM, protein= 6 μg/well) (all cDNA purchased from the cDNA Resource Center) were performed in buffer B together with the radioligands [$^3$H]WAY600135 (spec. act. 80 Ci/mmol; Biotrend) (5-$HT_{2A}$), [$^3$H]ketanserin (spec. act. 47 Ci/mmol; Biotrend) (5-$HT_{2A}$), and [$^3$H]N-methyl-LSD (spec. act. 81 Ci/mmol; Biotrend) (5-$HT_6$) at concentrations of 0.2 nM, 0.4 nM, and 0.75 nM, respectively. Unspecific binding was measured at 10 μM of non-labelled radioligand (5-$HT_{1A,}$ 5-$HT_{2A}$) or serotonin (5-$HT_6$). Muscarinic receptor binding was done with the $M_1$ ($B_{max}$= 4300 fmol/mg protein, $K_d$= 0.15 nM, protein= 2 μg/well), $M_2$ ($B_{max}$= 700 fmol/mg protein, $K_d$= 0.25 nM, protein= 10 μg/well), and $M_3$ ($B_{max}$= 4100 fmol/mg protein, $K_d$= 0.25 nM, protein= 2 μg/well) subtype (cDNA Resource Center) and the radioligand [$^3$H]N-methyl-scopolamin (spec. act. 75 Ci/mmol; Novandi) at 0.3 nM for $M_1$ and $M_2$, and 0.25 nM for $M_3$ in

buffer B. Unspecific binding was determined at 10 µM of atropine. Binding to the neurotensin receptors $NTS_1$ ($B_{max}$= 850 fmol/mg protein, $K_d$= 1.0 nM, protein= 10 µg/well) and $NTS_2$ ($B_{max}$= 880 fmol/mg protein, $K_d$= 1.6 nM, protein= 12 µg/well) was performed with cDNAs purchased from the cDNA Resource Center and the radioligand [$^3$H]NT(8-13) (spec. act. 130 Ci/mmol; custom synthesis by Novandi) at 0.7 nM and 0.8 nM for $NTS_1$ and $NTS_2$, respectively in buffer D (50 mM Tris, 0.1 mM EDTA, 0.1% bovine serum albumin at pH 7.4). Unspecific binding was measured at 10 µM of non-labelled NT(8-13). Binding affinities to the orexin receptor subtypes $OX_1$ (cDNA Resource Center, $B_{max}$= 3000 fmol/mg protein, $K_d$= 0.7 nM, protein= 5 µg/well) and $OX_2$ (cDNA from Genescript, Piscataway Township, NJ; $B_{max}$= 8000 fmol/mg protein, $K_d$= 1.1 nM, protein= 2 µg/well) were performed in buffer B together with the radioligands [$^3$H]SB674042 (conc: 0.7 nM; spec. act. 43 Ci/mmol; Novandi) for $OX_1$ and [$^3$H]EMPA (conc: 0.75 nM; spec. act. 77 Ci/mmol; Novandi) for $OX_2$, respectively.[36] Unspecific binding was determined in presence of 10 µM of non-labelled radioligand. Protein concentrations were determined employing the method of Lowry with bovine serum albumin as standard.[40]
The resulting competition curves of the receptor binding experiments were analyzed by nonlinear regression using the algorithms in PRISM 8.0 (GraphPad Software, San Diego, USA). The data were initially fit using a sigmoid model to provide $IC_{50}$ values which were subsequently transformed to $K_i$ values according to the equation of Cheng and Prusoff.[41]

**Tests on functional activity. G-protein IP-One assay and β-arrestin-2 recruitment assay**
The determination of receptor mediated G-protein signaling by µOR activation was performed applying an IP accumulation assay (IP-One HTRF®, Cisbio, Codolet, France) according to the manufacturer's protocol and in analogy to previously described protocols.[42] In brief, HEK 293T cells were co-transfected with the cDNA for µOR and the hybrid G-protein $G\alpha_{qi}$ ($G\alpha_q$ protein with the last five amino acids at the C-terminus replaced by the corresponding sequence of $G\alpha_i$ (gift from The J. David Gladstone Institutes, San Francisco, CA), respectively and transferred into 384 well micro plates. Cells were incubated with test compound for 120 min and accumulation of second messenger was stopped by adding detection reagents (IP1-d2 conjugate and Anti-IP1cryptate TB conjugate). After 60 min TR-FRET was measured with a Clariostar plate reader. FRET-signals were normalized to vehicle (0%) and the maximum effect of the reference fentanyl (100%). Three to twelve repeats in duplicate were analyzed applying the algorithms for four parameter non-linear regression implemented in Prism 8.0 (GraphPad, San Diego, CA) to get dose-response curves representing $EC_{50}$ and $E_{max}$ values.
Determination of µOR stimulated β-arrestin-2 recruitment was performed applying the PathHunter assay (DiscoverX, Birmingham, U.K.) as described.[43] In detail, based on the measurement of fragment complementation of β-galactosidase HEK293T cells stably expressing the enzyme acceptor (EA) tagged β-arrestin-2 were transfected with the cDNA for µOR fused to the ProLink-PK1 fragment for enzyme complementation alone or were co-transfected with the cDNA of the receptor and GRK2 (cDNA Resource Center) and subsequently transferred into 384 well micro plates. The assay started by incubating the cells with test compound for 90 min and was stopped by adding detection regent. Chemoluminescence data was normalized relative to basal activity (0%) and the maximum effect of fentanyl (100%). Three to ten repeats in duplicate were analyzed to get dose-response curves representing $EC_{50}$ and $E_{max}$ values.

**Bioluminescence resonance energy transfer (BRET) assays**
BRET assays were performed and analyzed as previously described[44] with the following modifications: HEK-293S cells grown in FreeStyle 293 suspension media (Thermo Fisher) were transfected at a density of 1 million cells/mL in 2 mL volume using 1200 ng total DNA at 1:1:1:1 ratio of receptor:Gα:Gβ:Gγ and a DNA:PEI ratio of 1:5, and incubated in a 24 deep well plate at 220 rpm, 37°C for 48 hours. Cells were harvested by centrifugation, washed with Hank's Balanced Salt Solution (HBSS) without Calcium/Magnesium (Gibco), and resuspended in assay buffer (HBSS with 20 mM HEPES pH 7.45) with 5 µg/mL freshly prepared coelenterazine 400a (GoldBio). Cells were then placed in white-walled, white-bottom

96 well plates (Costar) in a volume of 60 µl/well and 60,000 cells/well. Drug dilutions were prepared in drug buffer (assay buffer with 0.1% BSA, 6 mM CaCl2, 6 mM MgCl2), of which 30 µl were immediately added to plated cells. Ten minutes after the addition of ligand, plates were read using a SpectraMax iD5 plate reader using 410 nm and 515 nm emission filters with a one second integration time per well. The computed BRET ratios (GFP2/RLuc8 emission) were normalized to ligand-free control (Net BRET) prior to further analysis. Eleven-point dose-response curves in technical duplicate were analyzed by simultaneous curve-fitting of at least 3 biological replicates (minimum 66 data points/curve) using a log(dose) vs. response model in Prism 9.1.0 (GraphPad Software). All 95% confidence intervals for EC50 and Emax were asymmetrically calculated. Any additional details of analysis are described in figure and table legends.

**Chemical synthesis**

Enantiopure acryl amides were synthesized starting from amino acid precursors and acrylic/acetylenic acids, which were commercially available. Phenol analogs were obtained from *L*-tyrosine amide adapting a convergent synthesis route described in the literature[45]. *L*-Tyrosine amide was dimethylated in a reductive amination using formaldehyde and sodium triacetoxyborohydride. Borane reduction of the resulting amide FH184 to a primary amine gave the central intermediate FH185. Acrylic/acetylenic amides were synthesized from FH185 and acrylic/acetylenic acids under amide coupling conditions using (benzotriazol-1-yloxy)tris(dimethylamino)phosphonium hexafluorophosphate (BOP) or (benzotriazol-1-yloxy)tripyrrolidinophosphonium hexafluorophosphate (PyBOP) and triethyl amine (TEA). This final step led to the phenolic target structures FH163, FH172, FH210, FH217, FH218, FH219, FH223, FH273, FH299, FH315 and FH321.

A 4-carboxamido analog was prepared starting from Fmoc-protected 4-carboxamido *L*-phenylalanine. Mild reduction of the carboxylic acid with borane-THF complex at 0°C led to the primary alcohol FH282. In a two-step procedure, the alcohol was first oxidized to the aldehyde FH298 using Dess-Martin periodinane (DMP) and subsequently converted to the primary amine FH305 in a reductive amination applying titanium(IV) tetraisopropoxide as an additive. In a Henry reaction, thiophene-3-carbaldehyde was reacted with nitroethane to yield the nitropropene intermediate FH95, which was reduced with lithium alumniniumhydride to the racemic amine FH98. Chiral resolution with a tartaric acid derivative led to the enantiopure building block (*S*)-FH98, which was coupled with 4-nitrophenyl chloroformate to afford the activated carbamate (*S*)-FH99.

The two molecular fragments FH305 and (*S*)-FH99 were fused in an amide coupling reaction using the PyBOP reagent. In a one-pot fashion, the Fmoc-protecting group was readily cleaved by the addition of piperidine, leading to FH308. The primary amine was dimethylated in a reductive amination to yield the tertiary amine FH310.

### Supporting Scheme

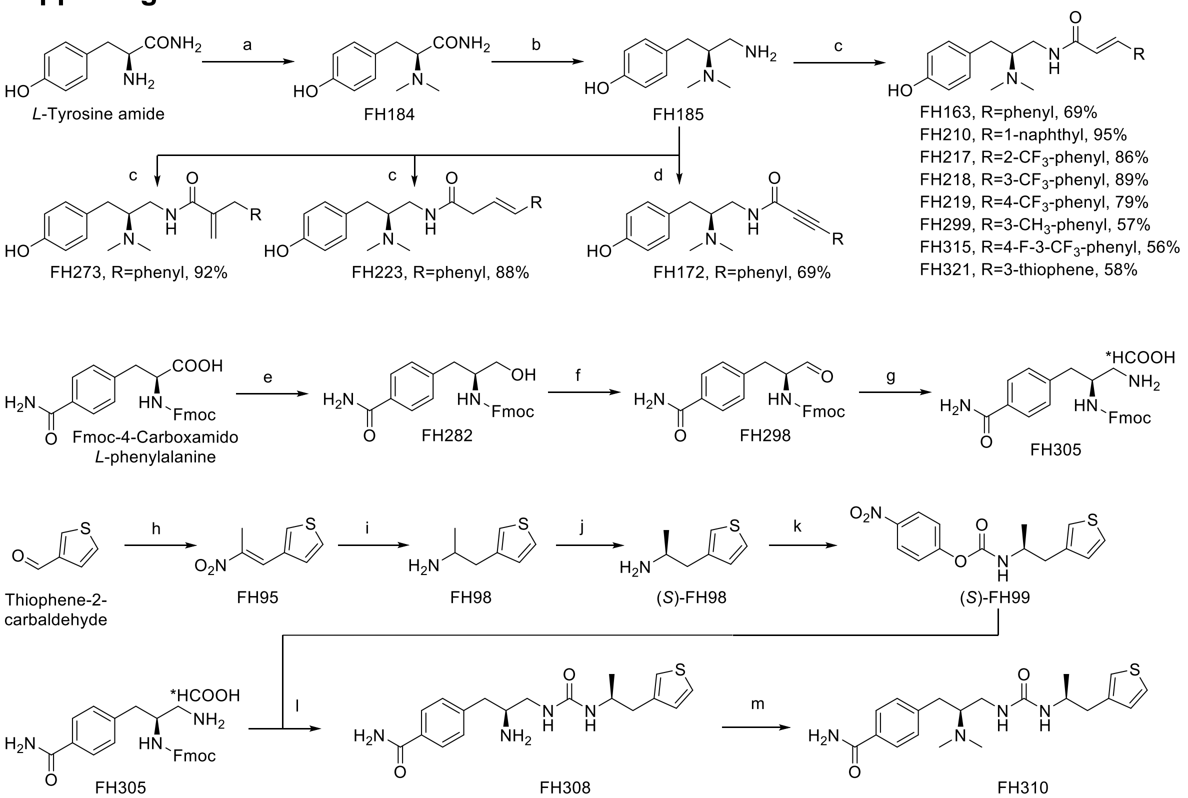


**Scheme S1**: Synthetic route for novel PZM21 analogs. Reagents and conditions: a: paraformaldehyde, $NaBH(OAc)_3$, ACN / $H_2O$, -10 °C, 10 min, 100 %, b: $BH_3$•THF, THF, argon, 0 °C – reflux, 6 h, 43 %, c: for FH163: (*E*)-cinnamic acid, BOP, TEA, DMF, r.t. 72 h, for FH210: (*E*)-3-(1-naphthyl) acrylic acid, PyBOP, TEA, DMF, r.t., 1 h, for FH217, FH218, FH219: *ortho*-, *meta*- or *para*-substituted (*E*)-(trifluormethyl) cinnamic acid, PyBOP, TEA, DMF, r.t., 30 min, for FH299: (*E*)-*m*-methyl cinnamic acid, PyBOP, TEA, DMF, r.t., 30 min, for FH315: (*E*)-4-fluoro-3-(trifluormethyl) cinnamic acid, PyBOP, TEA, DMF, r.t., 30 min, for FH321: (*E*)-3-(thiophen-3-yl) acrylic acid, PyBOP, TEA, DMF, r.t., 1 h, for FH273: 2-benzyl acrylic acid, PyBOP, TEA, DMF, r.t., 30 min; for FH223: (*E*)-4-phenylbut-3-enoic acid, PyBOP, TEA, DMF, r.t., 30 min, d: 3-phenyl propiolic acid, BOP, TEA, DMF, r.t. 23 h; e: $BH_3$•THF, THF, 0 °C, 24 DMP, THF, 0 °C, 6 h, 80 %, g: 1.) $Ti(OiPr)_4$, molecular sieves, THF, argon, - 50 °C, 15 min, 2.) $NH_4COOCF_3$, $NaBH_3CN$, THF, argon, - 50 °C, 2 h, - 20 °C, 3 h, 51 %; h: nitroethane, HCOOH, ethanolamine, 0 °C – 90 °C, 7 h, 83 %; i: $LiAlH_4$, THF, 0 °C – reflux, 30 min, 27 %; j: di-*para*-anisoyl-*D*-tartaric acid, recrystallization; k: *para*-nitrochloroformate, TEA, THF, 0 °C – r.t., 6 h, 79 %; l: 1.) TEA, DMF, r.t., 1 h, 2.) piperidine, DMF, r.t., 1 h, 74 %; m: 1.) paraformaldehyde, $NaBH(OAc)_3$, ACN / $H_2O$, - 10 °C – r.t., 1 h, 2.) 2M NaOH, ACN, r.t. 1 h, 35 %.

### Chemical synthesis material and methods

#### (*E*)-3-(2-Nitroprop-1-en-1-yl)thiophene[45] (FH95)

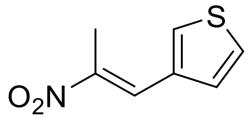

**FH95** was synthesized according to a procedure described in the literature[45]. Thiophene-3-carbaldehyde (5.47 mL, 62 mmol, 1 eq.) and nitroethane (17.85 mL, 250 mmol, 4 eq.) were added to an ice-cooled mixture of formic acid (10.6 mL, 281 mmol, 4.5 eq.) and ethanolamine (11.33 mL, 187 mmol, 3 eq.). The reaction mixture was heated at 85 − 90°C for 7 h. The resulting solution was poured into cold water (300 mL), and the slurry was filtered. The precipitated product was washed with water (3 × 50 mL) yielding a yellow solid. The crude product was recrystallized from ethanol/water (4:1 v/v) to give an orange crystalline solid (8.78 g, 52 mmol, 83 %). ESI (*m/z*) 266.95 $[M+H]^+$, $^1H$ NMR (400 MHz, 298K, Chloroform-*d*, TMS) δ (ppm): 8.09 (s, 1H), 7.60 (dt, *J* = 3.2, 0.9 Hz, 1H), 7.44 (dd, *J* = 5.1, 2.9 Hz, 1H), 7.28 (dd, *J* = 5.1, 1.3 Hz, 1H), 2.51 (d, *J* = 0.9 Hz, 3H).

**1-(Thiophen-3-yl)propan-2-amine hydrochloride[45] (FH98)**

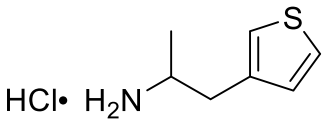

**FH98** was synthesized according to a procedure described in the literature[45]. To a solution of **FH95** (8.78 g, 52 mmol, 1 eq.) in 40 mL of dry THF was slowly added a solution of lithium aluminium hydride (64.9 mL, 260 mmol, 4 M in THF). After completion of addition, the reaction was refluxed for 30 min. The mixture was cooled to 0 °C where first 10 mL water, then 10 mL of a 15 wt. % solution of sodium hydroxide and magnesium sulfate (1 g) was added and allowed to stir for additional 15 min. The suspension was filtered and washed with ethyl acetate (500 mL). The combined organic phases were evaporated. The residue was diluted with MTBE (20 mL) and washed with 1 N HCl (three times 20 mL). The combined aqueous phases were basified with ammonia (25 %, aq.) and extracted with MTBE (three times 20 mL). The combined organic phases were dried over sodium sulfate and evaporated. The yellow oily residue was diluted with 20 mL of MTBE and HCl (52 mL, 52 mmol, 1 M in diethyl ether, 1 eq.) was added. The brown precipitate was filtered and recrystallized from 10 mL acetonitrile to give the product as a white solid (2.50 g, 14 mmol, 27 %). Analytical data matched those reported in the literature[45]. ESI (*m/z*) 141.93 $[M+H]^+$, $^1H$ NMR (600 MHz, 298K, DMSO-$d_6$, TMS) δ (ppm): 8.06 (s, 3H), 7.53 (dd, *J* = 4.9, 3.0 Hz, 1H), 7.32 (dd, *J* = 3.0, 1.2 Hz, 1H), 7.04 (dd, *J* = 4.9, 1.2 Hz, 1H), 3.42 (m, 1H), 2.98 (dd, *J* = 14.0, 5.1 Hz, 1H), 2.75 (dd, *J* = 14.0, 9.0 Hz, 1H), 1.12 (d, *J* = 6.5 Hz, 3H).

**(*S*)-1-(Thiophen-3-yl)propan-2-amine hydrochloride ((*S*)-FH98) (Chiral resolution)**

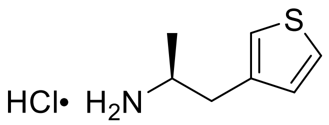

Compound **(*S*)-FH98** was synthesized by adapting a procedure described in the literature[45]. Racemic **FH98** (120 mg, 0.68 mmol) was dissolved in 3 mL of dist. water, basified with ammonia (25 % aq.) and extracted with chloroform (three times 3 mL). The combined organic phases were dried over sodium sulfate and evaporated to give a light yellow liquid (94 mg, 0.67 mmol). The residue was dissolved in ethanol (1 mL) and added to a hot solution of di-*para*-anisoyl-*D*-tartaric acid (278 mg, 0.67 mmol) in hot acetonitrile (2 mL). The white suspension was added dist. water (1 mL) and subsequently ethanol at boiling temperature until a clear solution was obtained. Slow cooling resulted in the formation of crystals, which were filtered off and washed with acetonitrile. Recrystallization from a mixture of water : ethanol (3 : 1) was repeated three times. The product (107 mg, 191 µmol) was dissolved in 3 mL of saturated sodium bicarbonate solution and extracted three times with 3 mL of dichloromethane. The combined organic phases were dried over sodium sulfate and evaporated to dryness to give the product as a yellow oil. To do so, the hydrochloride salt was obtained by the addition of hydrogen chloride (1M solution in diethyl ether, 0.29 mL, 0.29 mmol, 1.5 eq.) in a 1 mL diethyl ether solution. A white suspension formed and the solvent was evaporated in vacuum to give a white solid (27 mg, 0.19 mmol, 29 %). Analytical data matched those reported in the literature[45]. Specific optical rotation was measured and compared to the literature reference[45]. $[\alpha]_D^{25}$: + 15.5° (c = 0.40, $H_2O$), Ref.[45]: $[\alpha]_D^{25}$:+ 15.5° (c = 1.25, $H_2O$).

**($S$)-4-Nitrophenyl (1-(thiophen-3-yl)propan-2-yl)carbamate[45] ($S$-FH99)**

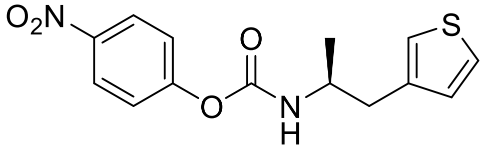

Compound **($S$)-FH99** was synthesized according to a procedure described in the literature[45]. *para*-Nitrochloroformate (79 mg, 0.39 mmol, 1 eq.) in 1 mL dry THF was added to an ice-cooled solution of 70 mg (0.39 mmol, 1 eq.) of **($S$)-9** and TEA (60 µL, 0.43 mmol, 1.1 eq.) in 1 mL dry THF. The reaction was stirred at room temperature for 6 h and then diluted with 2 mL of dichloromethane. The suspension was filtered and the organic phase washed with sat. sodium bicarbonate solution (5 mL), brine (5 mL), dried over sodium sulfate and evaporated under vacuum. The crude was purified by flash column chromatography (eluent: DCM : MeOH, $R_f$=0.6) to yield the title compound as a white solid (95 mg, 0.31 mmol, 79 %). Analytical data matched those reported in the literature[45]. $[\alpha]_D^{22}$: - 44.2° (c = 0.5, $CHCl_3$), Ref.[45]: $[\alpha]_D^{22}$: - 44.9° (c = 0.5, $CHCl_3$). $^1$H NMR (400 MHz, 298K, Chloroform-*d*, TMS) δ (ppm): 8.25 – 8.19 (m, 2H), 7.30 (dd, $J$ = 4.9, 3.0 Hz, 1H), 7.28 – 7.24 (m, 2H), 7.05 – 7.02 (m, 1H), 6.97 (dd, $J$ = 4.9, 1.3 Hz, 1H), 4.94 (d, $J$ = 8.4 Hz, 1H), 4.12 – 3.99 (m, 1H), 2.88 (d, $J$ = 6.3 Hz, 2H), 1.24 (d, $J$ = 6.6 Hz, 3H).

**(*S*)-*N*-(2-(Dimethylamino)-3-(4-hydroxyphenyl)propyl)cinnamamide (FH163)**

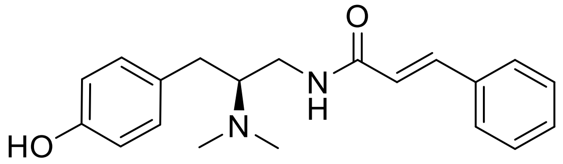

To a solution of **FH185** (13 mg, 67 µmol, 1 eq.) in 0.5 mL dry DMF was (*E*)-cinnamic acid (12 mg, 80 µmol, 1.2 eq.), BOP (35 mg, 80 µmol, 1.2 eq.) and triethylamine (65 µL, 0.47 mmol, 7 eq.) in 0.5 mL dry DMF. After stirring for 72 h at room temperature the solvent was removed under vacuum. The crude oil was dissolved in 5 mL of dichloromethane and extracted three times with 5 mL of saturated sodium bicarbonate solution. The organic phase was dried over sodium sulfate and evaporated to dryness. The crude was purified by column chromatography ($SiO_2$, solvent system: DCM : MeOH + 0.1 % ammonia (25 % aq.), 20 : 1, $R_f$=0.24). The product was obtained as a white solid (15 mg, 46 µmol, 69 %). Mp: 164 °C, $[\alpha]_D^{26}$: + 92.0° (c = 0.486, $CHCl_3$), ESI (*m/z*): 325.21 $[M+H]^+$, HRMS (*m/z*): $[M+H]^+$ calculated for $C_{20}H_{24}N_2O_2$: 325.1911, found: 325.1908. IR (NaCl): *υ* = 1629 (vs), 1514 (m) (C=O), 1660 (m) (C=C). $^1H$ NMR: (400 MHz, 298K, $CDCl_3$, TMS) δ 7.59 (d, *J* = 15.6 Hz, 1H), 7.52 – 7.46 (m, 2H), 7.38 – 7.32 (m, 3H), 7.00 – 6.96 (m, 2H), 6.82 – 6.78 (m, 2H), 6.65 – 6.60 (m, 1H), 6.40 (d, *J* = 15.6 Hz, 1H), 3.60 (ddd, *J* = 14.0, 6.7, 4.5 Hz, 1H), 3.14 – 3.00 (m, 1H), 2.93 (dd, *J* = 13.3, 3.3 Hz, 1H), 2.81 – 2.71 (m, 1H), 2.38 (s, 6H), 2.29 (dd, *J* = 13.3, 10.5 Hz, 1H). $^{13}C$ NMR: (151 MHz, 298K, $CDCl_3$, TMS) δ 166.3, 155.4, 141.2, 134.8, 129.9, 129.6, 129.4, 128.7, 127.9, 120.5, 115.7, 65.0, 40.2, 39.7, 30.5.

**(*S*)-N-(2-(Dimethylamino)-3-(4-hydroxyphenyl)propyl)-3-phenylpropiolamide (FH172)**

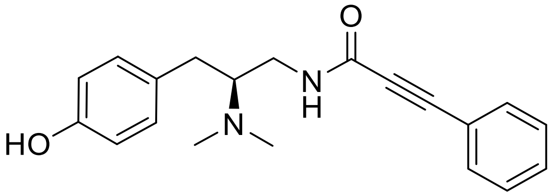


To a solution of **FH185** dihydrochloride (18 mg, 67 µmol) in 0.5 mL dry DMF was added 3-phenylpropiolic acid (12 mg, 81 µmol, 1.2 eq.), BOP (36 mg, 81 µmol, 1.2 eq.) and triethylamine (66 µL, 0.47 mmol, 7 eq.) in 0.5 mL dry DMF. After stirring for 23 h at room temperature the solvent was removed under vacuum. The crude oil was dissolved in 5 mL of dichloromethane and extracted three times with 5 mL of saturated sodium bicarbonate solution. The organic phase was dried over sodium sulfate and evaporated to dryness.The crude was first passed through a silica plug ($SiO_2$, ethyl acetate + 0.1 % TEA) and then purified by column chromatography (solvent system: DCM : MeOH + 0.1 % ammonia (25 % aq.), 20 : 1, $R_f$=0.13). The product was obtained as a white solid (16 mg, 50 µmol, 74 %). Mp:152 - 153 °C, $[\alpha]_D^{26}$: + 10.5° (c = 0.523, $CHCl_3$), ESI (m/z) 323.16 $[M+H]^+$, HRMS (*m/z*): $[M+H]^+$ calculated for $C_{20}H_{22}N_2O_2$: 323.1754, found: 323.1753. IR (NaCl): *υ* = 1624 (vs), 1452 (m) (C=O), 2213 (m) (alkyne). $^1H$ NMR (400 MHz, 298K, $CDCl_3$, TMS) δ (ppm): 7.55 – 7.51 (m, 2H), 7.43 – 7.32 (m, 3H), 7.01 – 6.96 (m, 2H), 6.79 – 6.74 (m, 3H), 3.51 (ddd, *J* = 13.9, 6.8, 4.6 Hz, 1H), 3.05 – 2.97 (m, 1H), 2.93 (dd, *J* = 13.4, 3.5 Hz, 1H), 2.74 (m, 1H), 2.37 (s, 6H), 2.27 (dd, *J* = 13.4, 10.3 Hz, 1H). $^{13}C$ NMR (101 MHz, 298K, $CDCl_3$, TMS) δ (ppm): 154.7, 153.5, 132.5, 130.1, 130.1, 130.0, 128.5, 120.3, 115.6, 84.7, 83.1, 64.6, 40.1, 39.9, 30.5.

**(*S*)-2-(Dimethylamino)-3-(4-hydroxyphenyl)propanamide (FH184)[46]**

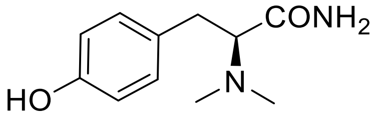


**FH184** has been described in the literature.[46] The compound was prepared following a previously reported procedure[45], solely the conversion to the hydrochloride salt was omitted. In a sealed microwave tube paraformaldehyde (4.30 g, 143 mmol, 10 eq.) was refluxed for two hours in an oil bath to prepare a 35 wt. % solution in water. *L*-Tyrosine amide (**1**) (2.58 g, 14.3 mmol, 1 eq.) was dissolved in a mixture of 60 mL acetonitrile and 12 mL distilled water and cooled to -10 °C in an acetone/ice bath. First, sodium triacetoxyborohydride (15.2 g, 71.5 mmol, 5 eq.), second, the formaldehyde solution was added. The reaction was quenched after 10 min by the addition of 40 mL saturated sodium bicarbonate and sodium carbonate solution to adjust to pH = 9. The slurry was diluted with 40 mL of dist. water. The aqueous phase was extracted five times with 100 mL of an organic solvent mixture of isopropanol : ethyl acetate 1 : 4. The combined organic phases were dried over sodium sulfate and evaporated to dryness. No further purification was necessary. The product was obtained as a white solid (2.98 g, 14.3 mmol, 100 %). Mp:105 °C, $[\alpha]_D^{24}$: + 35.6° (c = 0.44, MeOH), ESI (*m/z*)**:** 209 (40) $[M+H]^+$, 164 (60) $[M-CONH_2]^+$, IR (NaCl): $\upsilon$ = 1671 (vs) ((C=O)$NH_2$). $^1H$ NMR: (400 MHz, 298K, DMSO-$d_6$, TMS) δ 7.15 (d, *J* = 2.6 Hz, 1H), 6.99 – 6.94 (m, 2H), 6.84 (d, *J* = 2.6 Hz, 1H), 6.65 – 6.60 (m, 2H), 4.55 (s, 1H), 3.07 (dd, *J* = 9.1, 5.3 Hz, 1H), 2.80 (dd, *J* = 13.4, 9.1 Hz, 1H), 2.60 (dd, *J* = 13.4, 5.3 Hz, 1H), 2.23 (s, 6H). $^{13}C$ NMR: (151 MHz, 298K, DMSO-$d_6$, TMS) δ 171.8, 155.4, 129.7, 129.2, 114.7, 69.0, 41.5, 33.3.

**(*S*)-4-(3-Amino-2-(dimethylamino)propyl)phenol (FH185)**

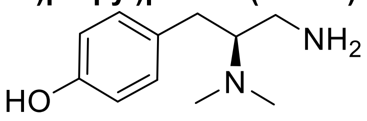

To an ice-cooled solution of **FH184** (3.00 g, 14.4 mmol, 1 eq.) in 60 mL of dry THF, borane-THF complex (86.4 ml, 86.4 mmol, 1M in THF, 6 eq.) was added dropwise at 0°C. After addition, the mixture was refluxed for six hours. The reaction was quenched by the slow addition of 20 mL dry methanol at reflux temperature in order to cleave the borane-amine complex. The mixture was refluxed for additional 30 minutes. The methanol addition-evaporation sequence was repeated twice. The crude product was subjected to column chromatography ($SiO_2$, eluent: DCM : MeOH + 0.1 % ammonia (25 % aq.), 10 : 1 to 4 : 1, $R_f$=0.2) and obtained as a white solid (1.20 g, 6.18 mmol, 43 %). Mp: 179 °C, $[\alpha]_D^{25}$: + 15.5° (c = 0.50, MeOH), ESI (*m/z*): 195 (40) $[M+H]^+$, 178 (60) $[M-NH_2]^+$, HRMS (*m/z*): $[M+H]^+$ calculated for $C_{11}H_{19}N_2O$: 195.1492, found: 195.1494. IR (NaCl): *υ* = 3333 (m) ($NH_2$). $^1H$ NMR: (400 MHz, 298K, DMSO-*d*$_6$, TMS) δ 6.97 – 6.91 (m, 2H), 6.68 – 6.62 (m, 2H), 2.67 (dd, *J* = 13.3, 3.8 Hz, 1H), 2.50 – 2.36 (m, 2H), 2.33 (dd, *J* = 11.4, 3.2 Hz, 1H), 2.23 (s, 6H), 2.16 (dd, *J* = 13.3, 8.7 Hz, 1H). $^{13}C$ NMR: (151 MHz, 298K, DMSO, TMS) δ 155.1, 130.5, 129.7, 114.9, 68.2.0, 40.1, 30.3.

**(*S*,*E*)-N-(2-(Dimethylamino)-3-(4-hydroxyphenyl)propyl)-3-(naphthalen-1-yl)acrylamide (FH210)**

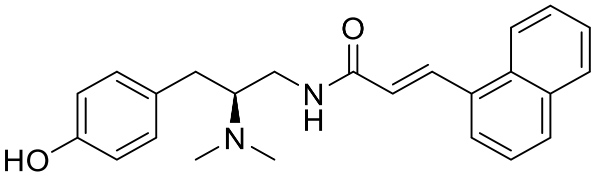

To a solution of **FH185** (60 mg, 0.31 mmol, 1 eq.) in 1 mL dry DMF was added (*E*)-3-(1-naphtyl) acrylic acid (61 mg, 0.31 mmol, 1 eq.), PyBOP (161 mg, 0.31 mmol, 1 eq.) and triethylamine (86 µL, 0.62 mmol, 2 eq.) in 1 mL dry DMF. The reaction flask was protected from light using aluminium foil. After stirring for 1 h at room temperature, the solvent was removed under high vacuum. The crude was purified by column chromatography ($SiO_2$, solvent system: DCM : ethyl acetate : MeOH + 0.1 % ammonia (25 % aq.), 90 : 5 : 1 to 90 : 5 : 5, $R_f$=0.15). The product was obtained as a colorless crystalline solid (110 mg, 0.29 mmol, 95 %). The product is light sensitive and should be handled in the dark. Mp: 90 °C, $[\alpha]_D^{25}$: + 30.0° (c = 0.36, MeOH), ESI (*m/z*): 375.08 $[M+H]^+$, HRMS (*m/z*): $[M+H]^+$ calculated for $C_{24}H_{26}N_2O_2$: 375.2067, found: 375.2066. IR (NaCl): *υ* = 1655 (m) (C=O), 1744 (m) (C=C). $^1$H NMR: (400 MHz, 298K, $CDCl_3$, TMS) δ (ppm): 8.44 (d, *J* = 15.3 Hz, 1H), 8.25 – 8.20 (m, 1H), 7.87 – 7.83 (m, 2H), 7.69 (dt, *J* = 7.2, 0.9 Hz, 1H), 7.58 – 7.48 (m, 2H), 7.47 – 7.42 (m, 1H), 7.04 – 7.00 (m, 2H), 6.86 – 6.81 (m, 2H), 6.66 – 6.59 (d, *J* = 5.2 Hz, 1H), 6.47 (d, *J* = 15.3 Hz, 1H), 3.66 (ddd, *J* = 14.0, 6.8, 4.5 Hz, 1H), 3.11 (ddd, *J* = 14.0, 10.4, 4.5 Hz, 1H), 2.95 (dd, *J* = 13.3, 3.3 Hz, 1H), 2.81 – 2.73 (m, 1H), 2.38 (s, 6H), 2.31 (dd, *J* = 13.3, 10.5 Hz, 1H). $^{13}$C NMR (151 MHz, 298K, $CDCl_3$, TMS) δ (ppm): 166.2, 155.4, 138.4, 133.6, 132.5, 131.5, 130.0, 129.9, 129.7, 128.5, 126.7, 126.1, 125.3, 124.6, 123.8, 123.4, 115.7, 65.1, 40.2, 39.8, 30.5.

**(*S*,*E*)-N-(2-(Dimethylamino)-3-(4-hydroxyphenyl)propyl)-3-(2-(trifluoromethyl)phenyl)acrylamide (FH217)**

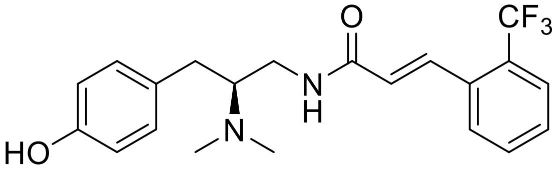


To a solution of **FH185** (15 mg, 77 µmol, 1 eq.) in 0.5 mL dry DMF was added (*E*)-*o*-(trifluoromethyl) cinnamic acid (17 mg, 77 µmol, 1 eq.), PyBOP (40 mg, 77 µmol, 1 eq.) and triethylamine (22 µL, 0.15 mmol, 2 eq.) in 0.5 mL dry DMF. After stirring for 30 min at room temperature the solvent was removed under vacuum. The crude was passed through a silica plug ($SiO_2$, ethyl acetate + 0.1 % TEA) and then purified by column chromatography ($SiO_2$, DCM : MeOH + 0.1 % ammonia (25 % aq.), 97 : 3, $R_f$=0.2). The product was obtained as a colorless crystalline solid (26 mg, 66 µmol, 86 %). Mp: 57 -59 °C, $[\alpha]_D^{25}$: + 39.3° (c = 0.603, MeOH), ESI (m/z) 393.01 $[M+H]^+$, HRMS (*m/z*) $[M+H]^+$ calculated for $C_{21}H_{23}F_3N_2O_2$: 393.1784, found: 393.1783. IR (NaCl): *υ* = 1666 (m) (C=O). $^1H$ NMR (400 MHz, 298K, Methanol-$d_4$, TMS) δ (ppm): 7.92 (dd, *J* = 15.5, 2.4 Hz, 1H), 7.81 (d, *J* = 7.8 Hz, 1H), 7.74 (d, *J* = 7.8 Hz, 1H), 7.65 (*pseudo*-t, *J* = 7.6 Hz, 1H), 7.55 (*pseudo*-t, *J* = 7.6 Hz, 1H), 7.15 – 7.09 (m, 2H), 6.80 – 6.72 (m, 2H), 6.57 (d, *J* = 15.5 Hz, 1H), 3.61 (dd, *J* = 15.1, 8.7 Hz, 1H), 3.42 – 3.34 (m, 2H), 3.05 (dd, *J* = 13.8, 4.1 Hz, 1H), 2.75 (s, 6H), 2.63 (dd, *J* = 13.7, 9.9 Hz, 1H). $^{13}C$ NMR (101 MHz, 298K, Methanol-$d_4$, TMS) δ (ppm): 168.6, 157.6, 137.7, 135.0, 133.7, 131.3, 130.8, 130.8 , 129.5 (q, *J* = 30.17 Hz, 129.1, 127.2 (q, *J* = 5.7 Hz), 125.7, 125.6 (q, *J* = 272.7 Hz, <u>C</u>$F_3$), 116.7, 68.2, 40.7, 40.1, 32.5.

**(*S*,*E*)-N-(2-(Dimethylamino)-3-(4-hydroxyphenyl)propyl)-3-(3-(trifluoromethyl)phenyl)acrylamide formate (FH218)**

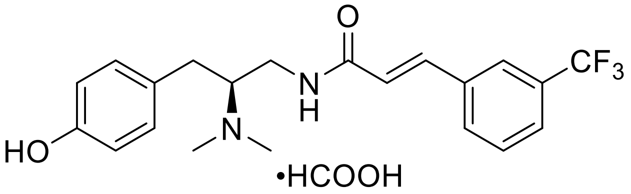


To a solution of **FH185** (15 mg, 77 µmol, 1 eq.) in 0.5 mL dry DMF was added (*E*)-*m*-(trifluoromethyl) cinnamic acid (17 mg, 77 µmol, 1 eq.), PyBOP (40 mg, 77 µmol, 1 eq.) and triethylamine (22 µL, 0.15 mmol, 2 eq.) in 0.5 mL dry DMF. After stirring for 30 min at room temperature the solvent was removed under vacuum. The crude was passed through a silica plug (eluent: ethyl acetate + 0.1 % TEA) and then purified by preparative HPLC (RP-C8, flow: 10 mL/min, solvent system: ACN/$H_2O$ + 0.1 % formic acid, 10 % ACN for 3 min, 10 % to 65 % ACN in 9 min, 65 % to 100 % ACN in 1 min, $t_R$ = 10.4 min). The product was obtained as a white solid (30 mg, 68 µmol, 89 %). Mp: 67 -69 °C, $[\alpha]_D^{24}$: + 13.1° (c = 0.865, MeOH), ESI (*m/z*)**:** 393.00 $[M+H]^+$, HRMS (*m/z*) $[M+H]^+$ calculated for $C_{21}H_{23}F_3N_2O_2$: 393.1784, found: 393.1785. IR (NaCl): *υ* = 1666 (m) (C=O). $^1$H NMR (600 MHz, 298K, Methanol-$d_4$, TMS) δ (ppm): 8.49 (s, 1H), 7.83 (s, 1H), 7.81 (d, *J* = 7.9 Hz, 1H), 7.67 (d, *J* = 7.7 Hz, 1H), 7.60 (*pseudo*-t, *J* = 7.8 Hz, 1H), 7.56 (d, *J* = 15.8 Hz, 1H), 7.14 – 7.10 (m, 2H), 6.79 – 6.75 (m, 2H), 6.67 (d, *J* = 15.8 Hz, 1H), 3.60 (dd, *J* = 14.7, 8.1 Hz, 1H), 3.44 – 3.35 (m, 2H), 3.06 (dd, *J* = 13.8, 4.2 Hz, 1H), 2.75 (s, 6H), 2.65 (dd, *J* = 13.8, 9.8 Hz, 1H). $^{13}$C NMR (151 MHz, 298K, Methanol-$d_4$, TMS) δ (ppm): 169.6, 169.0, 157.7, 140.5, 137.3, 132.4 (q, *J* = 32.4, <u>C</u>$CF_3$), 132.4, 131.3, 131.0, 128.9, 127.3 (q, *J* = 3.6 Hz, phenyl-C4), 125.4 (q, *J* = 3.8 Hz, phenyl-C2), 125.5 (q, *J* = 271.5 Hz, <u>C</u>$F_3$), 123.5, 116.8, 68.2, 40.8, 40.0, 32.7.

**(*S*,*E*)-N-(2-(Dimethylamino)-3-(4-hydroxyphenyl)propyl)-3-(4-(trifluoromethyl)phenyl)acrylamide (FH219)**

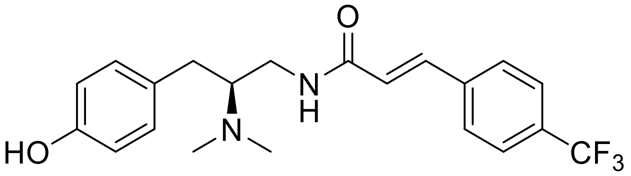

To a solution of **FH185** (15 mg, 77 µmol, 1 eq.) in 0.5 mL dry DMF was added (*E*)-*p*-(trifluoromethyl) cinnamic acid (17 mg, 77 µmol, 1 eq.), PyBOP (40 mg, 77 µmol, 1 eq.) and triethylamine (22 µL, 0.15 mmol, 2 eq.) in 0.5 mL dry DMF. After stirring for 30 min at room temperature the solvent was removed under vacuum. The crude was passed through a silica plug ($SiO_2$, ethyl acetate + 0.1 % TEA) and then purified by column chromatography ($SiO_2$, DCM : MeOH + 0.1 % ammonia (25 % aq.), 97 : 3, $R_f$=0.2). The product was obtained as a colorless crystalline solid (26 mg, 61 µmol, 79 %). Mp: 83 °C, $[\alpha]_D^{24}$: + 7.2° (c = 0.607, MeOH), ESI (*m/z*) 393.01 $[M+H]^+$, HRMS (*m/z*) $[M+H]^+$ calculated for $C_{21}H_{23}F_3N_2O_2$: 393.1784, found: 393.1784. IR (NaCl): *υ* = 1666 (m) (C=O). $^1$H NMR (600 MHz, 298K, Methanol-$d_4$, TMS) δ (ppm): 7.71 (d, *J* = 8.3 Hz, 2H), 7.67 (d, *J* = 8.3 Hz, 2H), 7.50 (d, *J* = 15.8 Hz, 1H), 7.07 – 7.05 (m, 2H), 6.74 – 6.71 (m, 2H), 6.68 (d, *J* = 15.8 Hz, 1H), 3.41 (dd, *J* = 14.3, 8.3 Hz, 1H), 3.37 (dd, *J* = 14.3, 5.2 Hz, 1H), 3.08 – 3.02 (m, 1H), 2.95 (dd, *J* = 13.7, 4.3 Hz, 1H), 2.52 – 2.44 (m, 7H). $^{13}$C NMR (151 MHz, 298K, Methanol-$d_4$, TMS) δ (ppm): 168., 157.1, 140.2, 139.9, 132.2 (q, *J* = 32.4 Hz, <u>C</u>CF$_3$), 131.1, 130.8, 129.3, 126.8 (q, *J* = 3.8 Hz, phenyl-C3,5), 125.5 (q, *J* = 271.0 Hz, <u>C</u>F$_3$), 124.7, 116.5, 67.1, 40.8, 40.6, 32.9.

**(*S*,*E*)-N-(2-(Dimethylamino)-3-(4-hydroxyphenyl)propyl)-4-phenylbut-3-enamide formiate (FH223)**

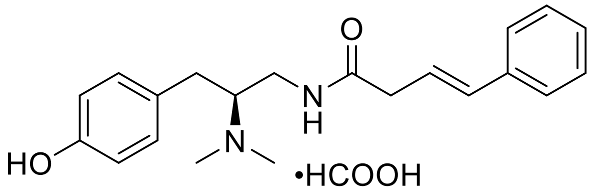

To a solution of **FH185** (15 mg, 77 µmol, 1 eq.) in 0.5 mL dry DMF was added (*E*)-4-phenylbut-3-enoic acid (13 mg, 77 µmol, 1 eq.), PyBOP (40 mg, 77 µmol, 1 eq.) and triethylamine (22 µL, 0.15 mmol, 2 eq.) in 0.5 mL dry DMF. After stirring for 30 min at room temperature the solvent was removed under vacuum. The crude was purified by preparative HPLC (RP-C8, flow = 7 mL/min, solvent system: ACN/$H_2O$ + 0.1 % formic acid, 5 % ACN for 3 min, 5 % to 40 % ACN in 17 min, 40 % to 95 % in 2 min, $t_R$ = 12.8 min). The product was obtained as a white solid (26 mg, 68 µmol, 88 %). Mp: 74 °C, $[\alpha]_D^{24}$: + 13.0° (c = 0.397, MeOH), ESI (*m/z*) 339.05 $[M+H]^+$, HRMS (*m/z*) $[M+H]^+$ calculated for $C_{21}H_{26}N_2O_2$: 339.2067, found: 339.2065. $^1$H NMR (400 MHz, 298K, Methanol-*d*$_4$, TMS) δ (ppm): 8.50 (s, 1H), 7.40 – 7.34 (m, 2H), 7.33 – 7.24 (m, 2H), 7.25 – 7.16 (m, 1H), 7.13 – 7.04 (m, 2H), 6.78 – 6.70 (m, 2H), 6.50 (dt, *J* = 15.8, 1.4 Hz, 1H), 6.26 (dt, *J* = 15.8, 7.2 Hz, 1H), 3.50 (dd, *J* = 13.9, 7.3 Hz, 1H), 3.36 – 3.32 (m, 1H), 3.30 – 3.25 (m, 1H), 3.09 (dd, *J* = 7.2, 1.4 Hz, 2H), 3.02 (dd, *J* = 13.9, 4.2 Hz, 1H), 2.72 (s, 6H), 2.61 (dd, *J* = 13.8, 9.5 Hz, 1H). $^{13}$C NMR (151 MHz, 298K, Methanol-*d*$_4$, TMS) δ (ppm): 175.0, 169.6, 157.6, 138.4, 135.1, 131.2, 129.6, 128.9, 128.6, 127.3, 123.5, 116.7, 67.9, 40.9, 40.8, 40.0, 32.8.

**(*S*)-2-Benzyl-*N*-(2-(dimethylamino)-3-(4-hydroxyphenyl)propyl)acrylamide (FH273)**

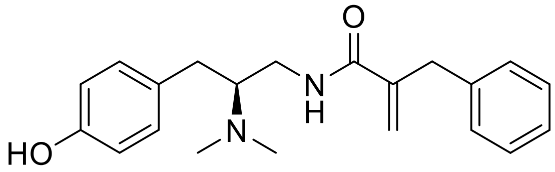


To a solution of **FH185** (20 mg, 0.10 mmol, 1 eq.) in 0.5 mL dry DMF was added 2-benzyl acrylic acid (17 mg, 0.10 mmol, 1 eq.), PyBOP (54 mg, 0.10 mmol, 1 eq.) and triethylamine (29 µL, 0.20 mmol, 2 eq.). After stirring for 30 min at room temperature the solvent was removed under vacuum. The white residue was dissolved in 5 mL of dichloromethane and extracted three times with 10 mL of 1 N HCl. The combined aqueous phases were basified with ammonia (25 % aq.) and a white precipitate formed. The suspension was filtered. The solid was washed with distilled water and dried under high vacuum. The product was obtained as a white solid and no further purification was necessary (32 mg, 95 µmol, 92 %). Mp: 170 °C, $[\alpha]_D^{25}$: + 32.8° (c = 0.50, MeOH), ESI (*m/z*) 339.17 $[M+H]^+$, HRMS (*m/z*) $[M+H]^+$ calculated for $C_{21}H_{26}N_2O_2$: 339.2067, found: 339.2066. IR (NaCl): *υ* = 1739 (vs) (C=O). $^1H$ NMR (400 MHz, 298K, Methanol-*d*$_4$, TMS) δ (ppm): (ppm): 7.28 – 7.23 (m, 2H), 7.20 – 7.14 (m, 3H), 6.97 – 6.92 (m, 2H), 6.70 – 6.66 (m, 2H), 5.65 (q, *J* = 0.8 Hz, 1H), 5.25 (q, *J* = 1.3 Hz, 1H), 3.58 (s, 2H), 3.19 (d, *J* = 6.5 Hz, 2H), 2.84 – 2.75 (m, 2H), 2.34 – 2.29 (m, 1H), 2.27 (s, 6H). $^{13}C$ NMR (151 MHz, 298K, Methanol-*d*$_4$, TMS) δ (ppm): 170.6, 156.9, 145.6, 139.8, 131.5, 131.0, 129.9, 129.5, 127.5, 120.7, 116.4, 66.2, 40.8, 40.6, 39.5, 33.4.

**(9*H*-Fluoren-9-yl)methyl (*S*)-(1-(4-carbamoylphenyl)-3-hydroxypropan-2-yl)carbamate (FH282)**

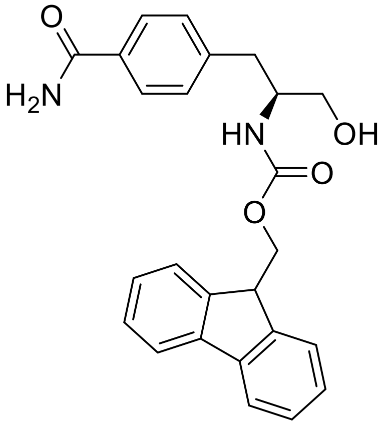

*N*-Fmoc-*L*-4-Carbamoylphenylalanine (1.00 g, 2.32 mmol, 1 eq.) was suspended in 20 mL of dry THF under nitrogen atmosphere. Borane-THF complex (1 M, 23.23 mL, 23.23 mmol) was slowly added at 0°C. The white suspension turns into a clear solution upon completion of addition. After stirring at 0°C for 24 h, 3 mL of methanol are slowly added in order to quench the reaction. Celite (1 g) was added to the solution and the solvent was evaporated. The crude was purified on silica gel, eluting with a gradient of DCM : MeOH 100 : 0 to 10 : 1, $R_f$=0.5. Remaining starting material could be recovered (63 mg, 0.15 mmol). The product was obtained as a white solid (550 mg, 1.32 mmol, respective yield 61 %.). Mp: 142 - 144 °C, $[\alpha]_D^{25}$, - 22.0° (c = 1.025, DMSO), ESI (*m/z*) 417.15 $[M+H]^+$, HRMS (*m/z*) $[M+Na]^+$ calculated for $C_{25}H_{24}N_2O_4$: 439.1628, found: 439.1622. IR (NaCl): *υ* = 3674 (m) (OH). NMR spectra were measured at 298 K and 318 K. At 298 K, two sets of signals were observed in the $^1H$ and $^{13}C$ NMR spectra. At 318 K, one single signal set was observed. This indicates that at 298 K the molecule adopts two distinct conformations. $^1H$ NMR (600 MHz, , T = 318 K, DMSO-$d_6$, TMS) δ (ppm): 7.87 (dt, *J* = 7.5, 1.0 Hz, 2H), 7.83 (dt, *J* = 7.5, 0.9 Hz, 2H), 7.79 – 7.77 (m, 3H), 7.41 (td, *J* = 7.4, 1.1 Hz, 2H), 7.34 (td, *J* = 7.4, 1.1 Hz, 2H), 7.27 (d, *J* = 8.1 Hz, 2H), 7.12 (s, 1H), 6.26 (s, 2H), 3.41 – 3.37 (m, 1H), 3.28 (dd, *J* = 10.4, 4.9 Hz, 1H), 3.19 (dd, *J* = 10.4, 6.4 Hz, 1H), 2.92 – 2.86 (m, 1H), 2.73 (dd, *J* = 13.3, 5.5 Hz, 1H), 2.49 – 2.45 (m, 1H), 1.46 – 1.42 (m, 1H). The signals for C<u>H</u> and C<u>H</u>$_2$ of the 9*H*-fluorenyl methyl group can be observed at 298 K but not at 318 K. $^1H$ NMR (600 MHz, T = 298 K, DMSO-$d_6$, TMS) δ (ppm): 4.81 (t, *J* = 5.6 Hz, 0.7H, 9*H*-fluorenyl methyl-C<u>H</u>CH$_2$), 4.37 (t, *J* = 5.2 Hz, 0.3H, 9*H*-fluorenyl methyl-C<u>H</u>CH$_2$), 4.26 – 4.14 (m, 2H, 9*H*-fluorenyl methyl-CHC<u>H</u>$_2$). $^{13}C$ NMR (151 MHz, T = 318 K, DMSO, TMS) δ (ppm): 167.8, 143.4, 142.6, 139.4, 137.4, 131.9, 128.9, 128.8, 127.2, 127.2, 121.2, 119.9, 109.4, 65.9, 60.7, 54.2, 40.0, 29.2.

**(9*H*-Fluoren-9-yl)methyl (*S*)-(1-(4-carbamoylphenyl)-3-oxopropan-2-yl)carbamate (FH298)**

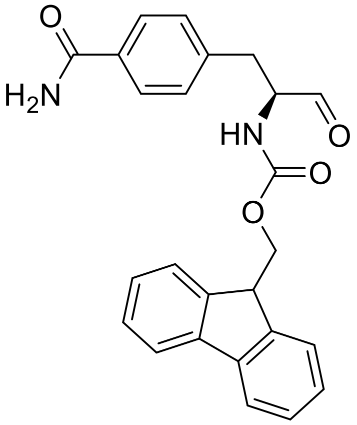


**FH282** (403 mg, 0.97 mmol, 1 eq.) was dissolved in 100 mL dry THF. The clear solution was cooled with an acetone/ice bath and Dess-Martin periodinane (616 mg, 1.45 mmol, 1.5 eq.) was added in small portions. The white suspension was stirred at 0 °C for 6 h. The solvent was evaporated in a nitrogen stream to avoid heating. Celite and 10 mL of water were added for lyophilizing overnight. Purification via a Biotage system (10 g, silica column) eluting with a ratio of 100 : 0 to 95 : 5 (DCM : MeOH) gave the product as a white solid (320 mg, 0.77 mmol, 80 %). Due to instability, the substance was directly used for the next step without further characterization. ESI (*m/z*) 415.06 $[M+H]^+$, HRMS (*m/z*) $[2M+H]^+$ calculated for $C_{25}H_{22}N_2O_4$: 829.3232, found: 829.3242. IR (NaCl): $\upsilon$ = 3331 (s), 3200 (m) (aldehyde, hydrated).

**(*S*,*E*)-*N*-(2-(Dimethylamino)-3-(4-hydroxyphenyl)propyl)-3-(*m*-tolyl)acrylamide (FH299)**

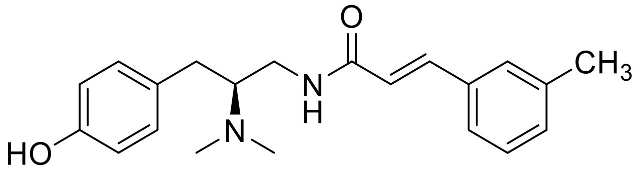

To a solution of **FH185** (15 mg, 77 µmol, 1 eq.) in 0.5 mL dry DMF was added (*E*)-*m*-methylcinnamic acid (13 mg, 77 µmol, 1 eq.), PyBOP (40 mg, 77 µmol, 1 eq.) and triethylamine (22 µL, 0.15 mmol, 2 eq.). After stirring for 30 min at room temperature the solvent was removed under vacuum. The crude was passed through a silica plug (eluent: ethyl acetate + 0.1 % TEA) and then purified by flash column chromatography (eluent: DCM : MeOH + 0.1 % ammonia (25 % aq.), 100 % : 0 % to 97 % :3 %, $R_f$=0.2). The product was obtained as a white solid (15 mg, 44 µmol, 57 %). Mp: 74 - 75 °C, $[\alpha]_D^{26}$: + 2.3° (c = 0.42, MeOH), ESI (*m/z*) 339.07 $[M+H]^+$, HRMS (*m/z*) $[M+H]^+$ calculated for $C_{21}H_{26}N_2O_2$: 339.2067, found: 339.2068. IR (NaCl): *υ* = 1614 (m), 1515 (m) (C=O), 1659 (m) (C=C). $^1$H NMR (600 MHz, 298K, Methanol-*d*$_4$, TMS) δ (ppm): 7.46 (d, *J* = 15.8 Hz, 1H), 7.35 (s, 1H), 7.33 (d, *J* = 7.8 Hz, 1H), 7.26 (t, *J* = 7.6 Hz, 1H), 7.18 (d, *J* = 7.6 Hz, 1H), 7.11 – 7.08 (m, 2H), 6.77 – 6.74 (m, 2H), 6.54 (d, *J* = 15.8 Hz, 1H), 3.52 (dd, *J* = 14.6, 8.4 Hz, 1H), 3.38 (dd, *J* = 14.6, 4.5 Hz, 1H), 3.28 – 3.22 (m, 1H), 3.01 (dd, *J* = 13.8, 4.4 Hz, 1H), 2.65 (s, 6H), 2.57 (dd, *J* = 13.8, 9.8 Hz, 1H), 2.34 (s, 3H). $^{13}$C NMR 151 MHz, 298K, Methanol-*d*$_4$, TMS) δ (ppm): 169.5, 157.4, 142.4, 139.8, 136.1, 131.7, 131.2, 129.9, 129.7, 129.5, 126.1, 121.2, 116.6, 67.8, 40.8, 40.2, 32.7, 21.3.

**(9*H*-Fluoren-9-yl)methyl (*S*)-(1-amino-3-(4-carbamoylphenyl)propan-2-yl)carbamate formiate (FH305)**

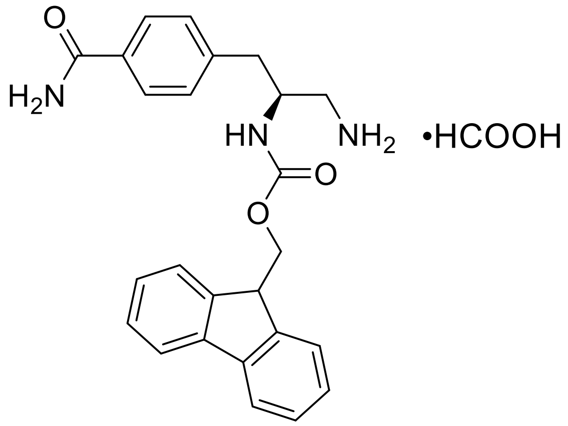


**FH298** (310 mg, 0.76 mmol, 1 eq.) was dissolved in 60 mL of dry THF in an argon atmosphere. Dry molecular sieves were added and the solution was cooled to – 50 °C. Titanium tetraisopropoxide (443 µL, 1.50 mmol, 2 eq.) was slowly added and the mixture was stirred for 15 min. Ammonium trifluoroacetate (980 mg, 1.48 mmol, 10 eq.) and sodium cyanoborohydride (141 mg, 2.24 mmol, 3 eq.) in 20 mL dry THF were slowly added. The mixture was stirred at – 50 °C for two hours and then at -20 °C for three hours. Molecular sieves were filtered off and washed with 20 mL of methanol. The combined organic solvent fractions were evaporated in a nitrogen stream. Purification was performed on a Biotage system (C18, 12 g column, eluent: acetonitrile : water + 0.1 % formic acid, 3 % to 100 % ACN in 30 column volumes). The product was obtained as a white solid (90 mg, 0.20 µmol, 37 %, 60 mg of the respective alcohol were recovered, overall yield: 51 %). Mp: 158 °C, ESI (*m/z*) 416.08 $[M+H]^+$, HRMS (*m/z*): $[M+H]^+$ calculated for $C_{25}H_{26}N_3O_3$: 416.1969, found: 416.1970. IR (NaCl): $\upsilon$ = 3354 (m) ($NH_2$). $^1H$ NMR (600 MHz, 298K, DMSO-$d_6$, TMS) δ (ppm): 8.24 (s, 1H), 7.90 – 7.82 (m, 4H), 7.81 – 7.79 (m, 2H), 7.63 – 7.60 (m, 2H), 7.44 – 7.39 (m, 2H), 7.37 – 7.28 (m, 4H), 7.26 (d, *J* = 8.0 Hz, 2H), 6.28 (s, 1H), 4.32 (dd, *J* = 9.9, 6.3 Hz, 1H), 4.22 – 4.14 (m, 2H), 3.87 – 3.80 (m, 1H), 2.88 – 2.80 (m, 2H), 2.78 – 2.69 (m, 2H). $^{13}C$ NMR (151 MHz, 298K, DMSO, TMS) δ (ppm): 167.5, 163.8, 155.7, 143.7, 142.5, 141.4, 140.6, 139.3, 137.3, 132.2, 128.9, 128.8, 127.5, 127.4, 127.2, 126.9, 125.0, 124.9, 121.3, 120.0, 109.6, 65.3, 51.4, 46.6, 42.7, 37.3. NMR signals appear as two sets of signals (ratio 4:1). This could result from an intramolecular hydrogen bond between the primary amine and the carbamate group (compare **FH282**). Only the major set of signals is reported.

**4-((*S*)-2-Amino-3-(3-((*S*)-1-(thiophen-3-yl)propan-2-yl)ureido)propyl)benzamide (FH308)**

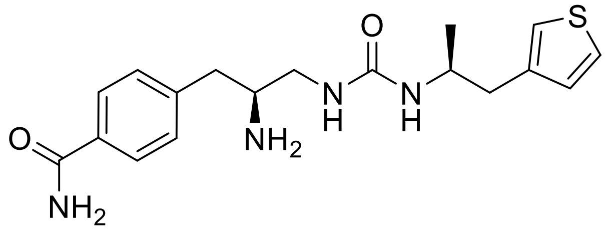

**FH305** (40 mg, 87 µmol) was dissolved in 1.5 mL dry DMF. The activated carbamate **(*S*)-FH99** (27 mg, 87 µmol, 1.0 eq.) in 1.5 mL of dry DMF was slowly added at 0 °C. TEA (12 µL, 87 µmol, 1 eq.) was added and the solution turned bright yellow. The mixture was stirred at r.t. for 1 h, after which piperidine (0.75 mL, 20 vol. % in DMF) was added. After stirring for 1 h at r.t., the solvent was evaporated under vacuum and the crude was purified by flash column chromatography (eluent: DCM : MeOH + 0.1 % ammonia (25 % aq.), 99 : 1 to 90 : 10, $R_f$=0.15). The product was obtained as a yellow sticky oil (23 mg, 64 µmol, 74 %). $[\alpha]_D^{26}$: + 12.8° (c = 0.63, MeOH), ESI (*m/z*) 361.01 $[M+H]^+$: HRMS (*m/z*) $[M+H]^+$ calculated for $C_{18}H_{24}N_4O_2S$: 361.1693, found: 361.1693. IR (NaCl): *υ* = 1661 (s) (C=O). $^1H$ NMR (600 MHz, 298K, DMSO-$d_6$, TMS) δ (ppm): 7.94 (s, 1H), 7.86 – 7.82 (m, 2H), 7.43 (dd, *J* = 4.9, 2.9 Hz, 1H), 7.35 – 7.29 (m, 3H), 7.16 (dd, *J* = 3.0, 1.2 Hz, 1H), 6.98 (dd, *J* = 4.9, 1.2 Hz, 1H), 6.18 (t, *J* = 5.9 Hz, 1H), 6.08 (d, *J* = 8.0 Hz, 1H), 3.85 – 3.78 (m, 1H), 3.29 – 3.22 (m, 1H), 3.17 – 3.11 (m, 1H), 3.06 – 2.98 (m, 1H), 2.79 (d, *J* = 6.9 Hz, 2H), 2.74 (dd, *J* = 13.9, 6.3 Hz, 1H), 2.63 (dd, *J* = 13.9, 6.9 Hz, 1H), 0.99 (d, *J* = 6.6 Hz, 3H). $^{13}C$ NMR (151 MHz, 298K, DMSO, TMS) δ (ppm): 167.5, 158.0, 140.5, 139.2, 132.5, 129.0, 128.8, 127.6, 125.4, 121.6, 52.7, 45.9, 42.1, 37.1, 36.9, 20.6.

**4-((*S*)-2-(Dimethylamino)-3-(3-((*S*)-1-(thiophen-3-yl)propan-2-yl)ureido)propyl)benzamide formiate (FH310)**

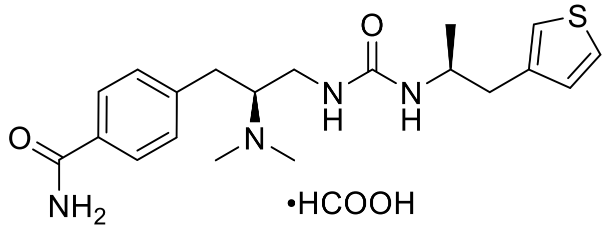


Formaldehyde was freshly prepared from paraformaldehyde (29 mg, 0.33 mmol, 10 eq., 35 % in water) by refluxing for 3 h in a sealed tube. Compound **FH308** (12 mg, 33 µmol, 1 eq.) was dissolved in a mixture of 1 mL water and 5 mL of acetonitrile. The solution was cooled in an acetone/ice bath. Sodium triacetoxyborohydride (35 mg, 0.17 mmol, 5 eq.) and subsequently formaldehyde was added. The reaction was allowed to warm to r.t. and then stirred for one hour. Saturated sodium bicarbonate solution (3 mL) and a spatula tip of sodium carbonate was added (pH = 9). The aqueous phase was extracted five times with 5 mL of ethyl acetate and evaporated to dryness. The crude was dissolved in 5 mL of acetonitrile and 1 mL of 2 M sodium hydroxide solution. The mixture was stirred at r.t. for 1 h, diluted with 5 mL of dist. water and extracted five times with 5 mL of ethyl acetate. The procedure was repeated two times until no more N-hydroxymethyl side product could be detected by LC-MS analysis. The crude was subjected to preparative HPLC (RP-C8, flow: 10 mL/min, solvent system: ACN/$H_2O$ + 0.1 % formic acid, 5 % ACN for 3 min, 5 % to 25 % ACN in 17 min, 25 % to 95 % in 2 min, $t_R$ = 16.2 min). The product was obtained as a white solid (5.0 mg, 12 µmol, 35 %). $[\alpha]_D^{26}$: + 32.0° (c = 0.04, MeOH), ESI (*m/z*) 389.02 $[M+H]^+$, HRMS (*m/z*) $[M+H]^+$ calculated for $C_{20}H_{28}N_4O_2S$: 389.2006, found: 389.2006. IR (NaCl): *υ* = 1658 (m) (C=O). $^1H$ NMR (400 MHz, 298K, DMSO-*d*$_6$, TMS) δ (ppm): 8.21 (s, 1H), 7.91 (s, 1H), 7.80 – 7.77 (m, 2H), 7.41 (dd, *J* = 4.9, 3.0 Hz, 1H), 7.29 – 7.24 (m, 3H), 7.11 (dd, *J* = 3.0, 1.2 Hz, 1H), 6.93 (dd, *J* = 4.9, 1.3 Hz, 1H), 6.01 (d, *J* = 7.9 Hz, 1H), 5.63 (dd, *J* = 7.4, 2.3 Hz, 1H), 3.72 (m, 1H), 3.04 (ddd, *J* = 12.6, 7.3, 4.7 Hz, 1H), 2.91 – 2.81 (m, 2H), 2.71 – 2.63 (m, 2H), 2.55 (dd, *J* = 14.3, 7.3 Hz, 1H), 2.43 – 2.33 (m, 1H), 2.26 (s, 6H), 0.93 (d, *J* = 6.5 Hz, 3H). $^{13}C$ NMR (151 MHz, 298K, DMSO, TMS) δ (ppm): 167.7, 163.5, 157.1, 143.9, 139.3, 131.8, 128.8, 128.7, 127.4, 125.3, 121.4, 65.0, 45.5, 40.0-39.0 (phenyl-$CH_2CH\underline{C}H_2$, signal under DMSO signal, proven by COSY-NMR), 39.9, 36.9, 31.2, 20.6.

**(*S*,*E*)-N-(2-Dimethylamino)-3-(4-hydroxyphenyl)propyl)-3-(4-fluoro-3-(trifluoromethyl)phenyl)acrylamide formiate (FH315)**

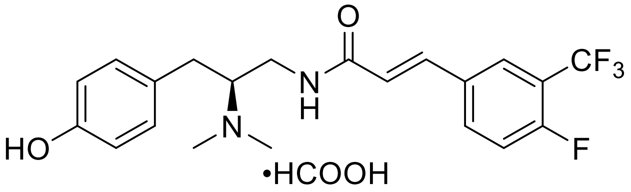


To a solution of **FH185** (35 mg, 0.18 mmol, 1 eq.) in 0.5 mL dry DMF was added (*E*)-4-fluoro-3-trifluormethyl cinnamic acid (42 mg, 0.18 mmol, 1 eq.), PyBOP (94 mg, 0.18 mmol, 1 eq.) and triethylamine (50 µL, 0.36 mmol, 2 eq.). After stirring for 30 min at room temperature the solvent was removed under vacuum. The crude was passed through a silica plug (eluent: ethyl acetate + 0.1 % TEA) and then purified by preparative HPLC (RP-C8, flow: 10 mL/min, solvent system: ACN/$H_2O$ + 0.1 % formic acid, 5 % ACN for 3 min, 5 % to 71 % ACN in 15 min, 71 % to 100 % in 3 min, $t_R$ = 13.0 min). The product was obtained as a white solid (46 mg, 0.10 mmol, 56 %). Mp: 106 - 108 °C, $[\alpha]_D^{25}$: - 12.5° (c = 2.05, MeOH), ESI (*m/z*) 411.08 $[M+H]^+$, HRMS (*m/z*) $[M+H]^+$ calculated for $C_{21}H_{22}F_4N_2O_2$: 411.1690, found: 411.1688. IR (NaCl): *υ* = 1663 (m) (C=O), 1613 (s) (C=C). $^1H$ NMR (600 MHz, 298K, DMSO-$d_6$, TMS) δ (ppm): 8.17 (s, 1H), 7.98 (dd, *J* = 7.0, 2.2 Hz, 1H), 7.94 (ddd, *J* = 8.4, 4.9, 2.2 Hz, 1H), 7.82 – 7.79 (m, 1H), 7.56 (dd, *J* = 10.6, 8.7 Hz, 1H), 7.43 (d, *J* = 15.8 Hz, 1H), 7.03 – 7.00 (m, 2H), 6.84 (d, *J* = 15.8 Hz, 2H), 6.70 – 6.67 (m, 2H), 3.22 (ddd, *J* = 14.0, 6.0, 4.8 Hz, 1H), 3.16 (ddd, *J* = 14.0, 8.3, 4.5 Hz, 1H), 2.83 – 2.75 (m, 2H), 2.36 – 2.30 (m, 7H). $^{13}C$ NMR (151 MHz, 298K, DMSO-$d_6$, TMS) δ (ppm): 164.3, 163.2, 158.9 (d, *J* = 256.0 Hz, phenyl-C4), 155.4, 135.6, 133.9 (d, *J* = 8.9 Hz), 132.3 (d, *J* = 3.6 Hz, phenyl-C1), 129.7 , 129.6, 125.9 (q, *J* = 4.3 Hz, phenyl-C2), 124.4 (d, *J* = 1.6 Hz, amide-<u>C</u>H=CH), 122.4 (q, *J* = 272.2 Hz, <u>C</u>$F_3$), 117.8 (d, *J* = 20.9 Hz, phenyl-C5), 117.0 (qd, *J* = 32.4, 12.7 Hz, phenyl-C3), 115.0, 64.9, 40.0, 38.4, 30.9.

**(*S*,*E*)-*N*-(2-(Dimethylamino)-3-(4-hydroxyphenyl)propyl)-3-(thiophen-3-yl)acrylamide (FH321)**

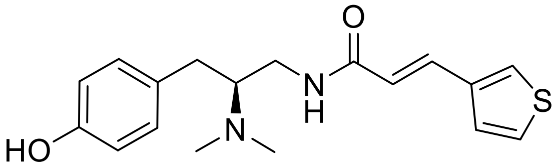

To a solution of **FH185** (20 mg, 120 µmol, 1 eq.) in 0.5 mL dry DMF was added (*E*)-3-(thiophen-3-yl)acrylic acid (19 mg, 120 µmol, 1.0 eq.), PyBOP (63 mg, 120 µmol, 1.0 eq.) and TEA (34 µL, 0.24 mmol, 2 eq.) in 0.5 mL dry DMF. After stirring for 1 h at room temperature the solvent was removed under vacuum. The crude oil was dissolved in 5 mL of dichloromethane and extracted three times with 5 mL of saturated sodium bicarbonate solution. The organic phase was extracted with four times 5 mL of 2 N HCl solution. The combined acidic phases were basified to pH = 10 with 6 N NaOH solution and extracted five times with 5 mL of DCM to remove side products. Afterwards, the basic phase was extracted four times with 5 mL of ethyl acetate to extract the product. The combined ethyl acetate phases were dried over sodium sulfate and evaporated to dryness. The product was obtained as a white solid (23 mg, 70 µmol, 58 %). Further purification was not necessary. Mp: 168 – 170 °C, $[\alpha]_D^{24}$: + 28.9 ° (c = 0.56, MeOH), ESI (*m/z*) 331.18 $[M+H]^+$, HRMS (*m/z*) $[M+H]^+$ calculated for $C_{18}H_{22}N_2O_2S$: 331.1475, found: 331.1476. $^1H$ NMR (600 MHz, 298K, $CDCl_3$, TMS) δ (ppm): 7.57 (d, *J* = 15.6 Hz, 1H), 7.43 (dd, *J* = 2.8, 1.2 Hz, 1H), 7.30 (dd, *J* = 5.1, 2.9 Hz, 1H), 7.28 – 7.25 (m, 1H), 7.02 – 6.96 (m, 2H), 6.81 – 6.76 (m, 2H), 6.27 (d, *J* = 15.5 Hz, 1H), 3.66 – 3.58 (m, 1H), 3.13 – 3.07 (m, 1H), 2.95 (dd, *J* = 13.5, 3.6 Hz, 1H), 2.89 – 2.83 (m, 1H), 2.43 (s, 6H), 2.40 – 2.32 (m, 1H). $^{13}C$ NMR (151 MHz, 298K, $CDCl_3$, TMS) δ (ppm): 166.6, 155.4, 137.8, 134.8, 130.0, 129.2, 127.2, 126.6, 125.2, 120.2, 115.8, 65.3, 40.2, 39.5, 30.7.

**$^{1}$H-NMR and $^{13}$C-NMR spectra**

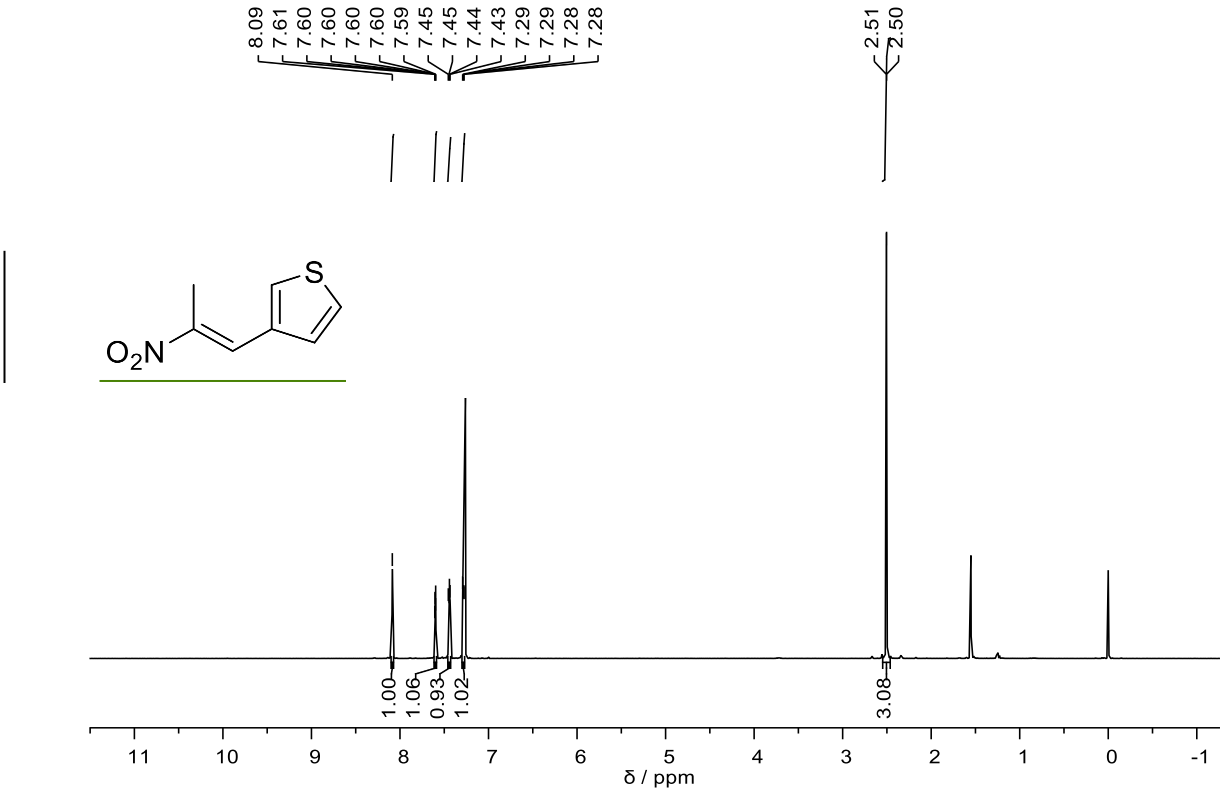


400MHz $^{1}$H NMR spectrum of **FH95** at 298K in $CDCl_3$.

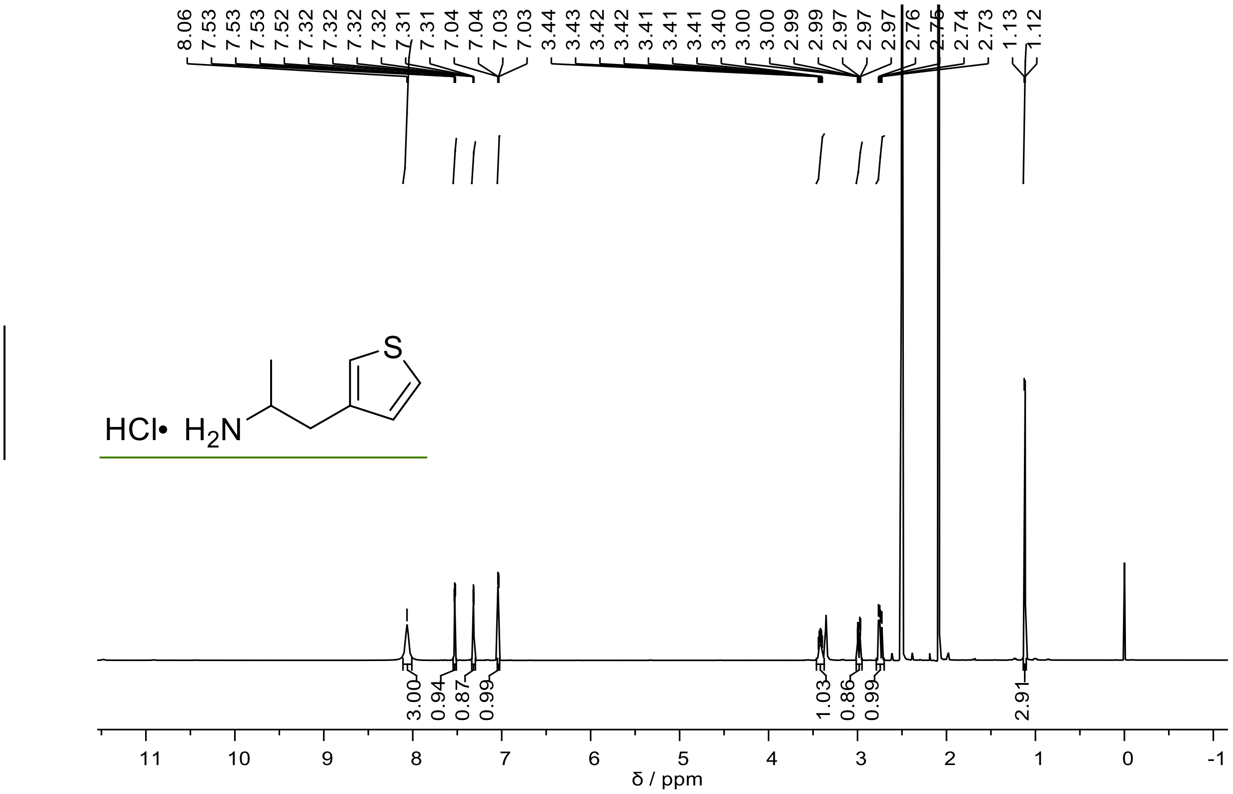


600MHz [1]H NMR spectrum of **FH98** at 298K in DMSO-$d_6$.

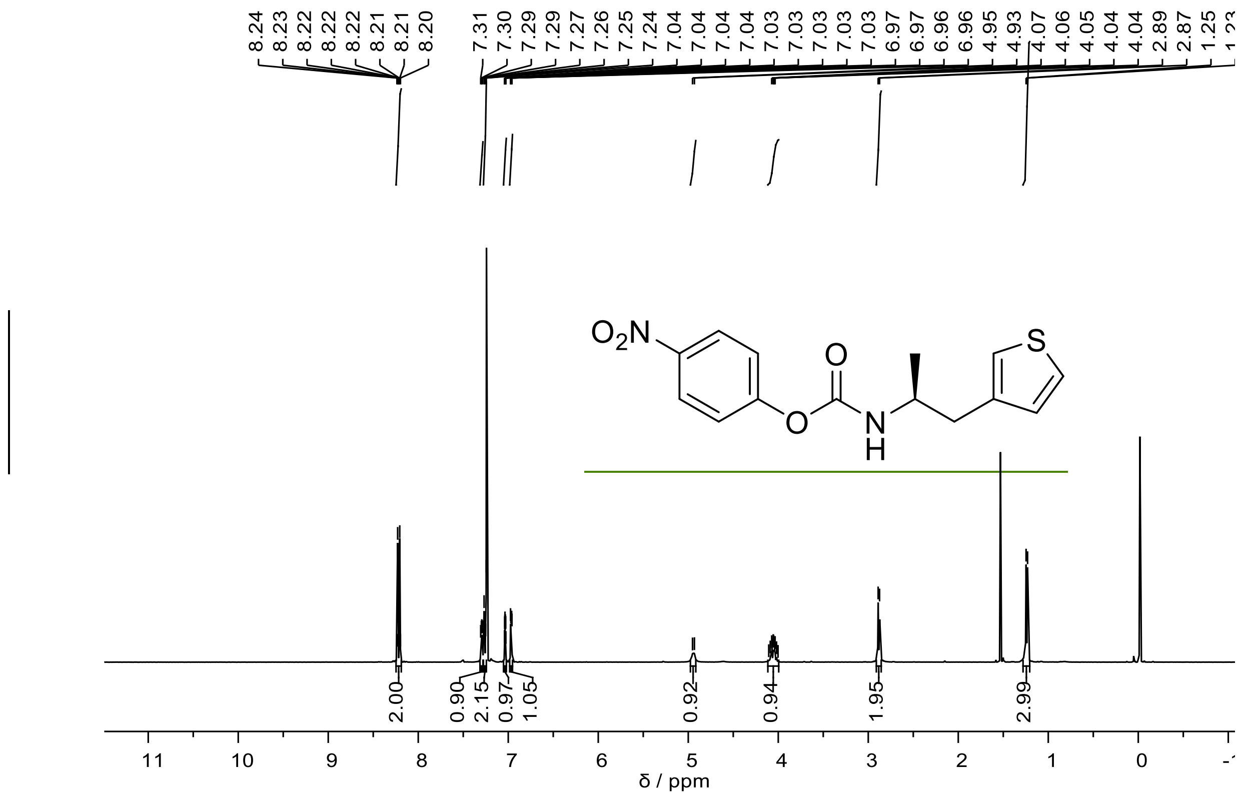

151MHz $^{1}$H NMR spectrum of **(*S*)-FH99** at 298K in $CDCl_3$.

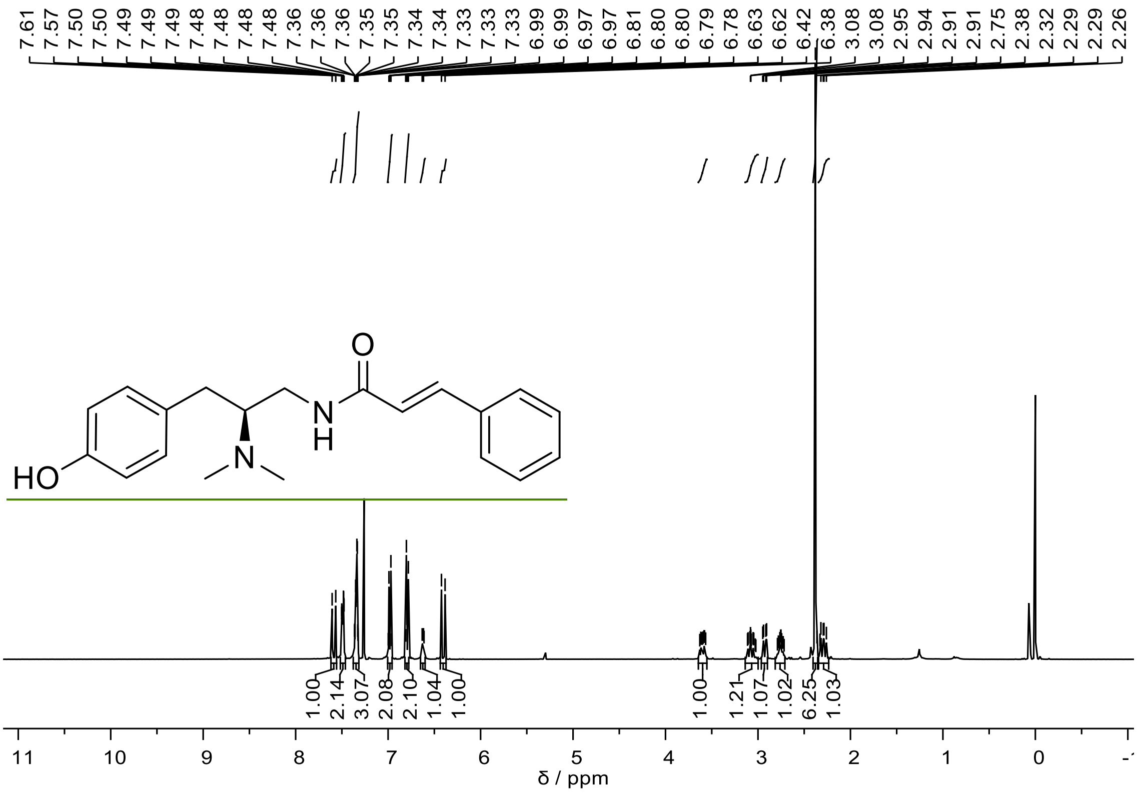


400MHz $^{1}$H NMR spectrum of **FH163** at 298K in $CDCl_3$.

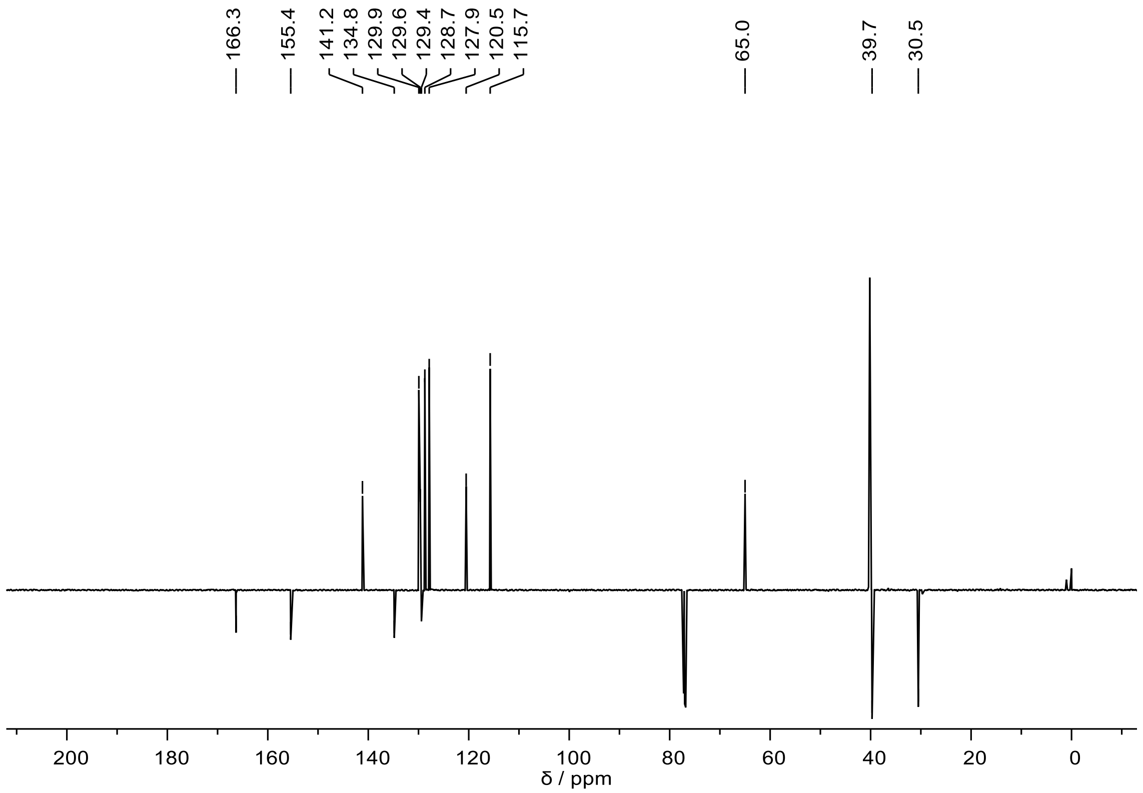


151MHz $^{13}$C NMR spectrum of **FH163** at 298K in $CDCl_3$.

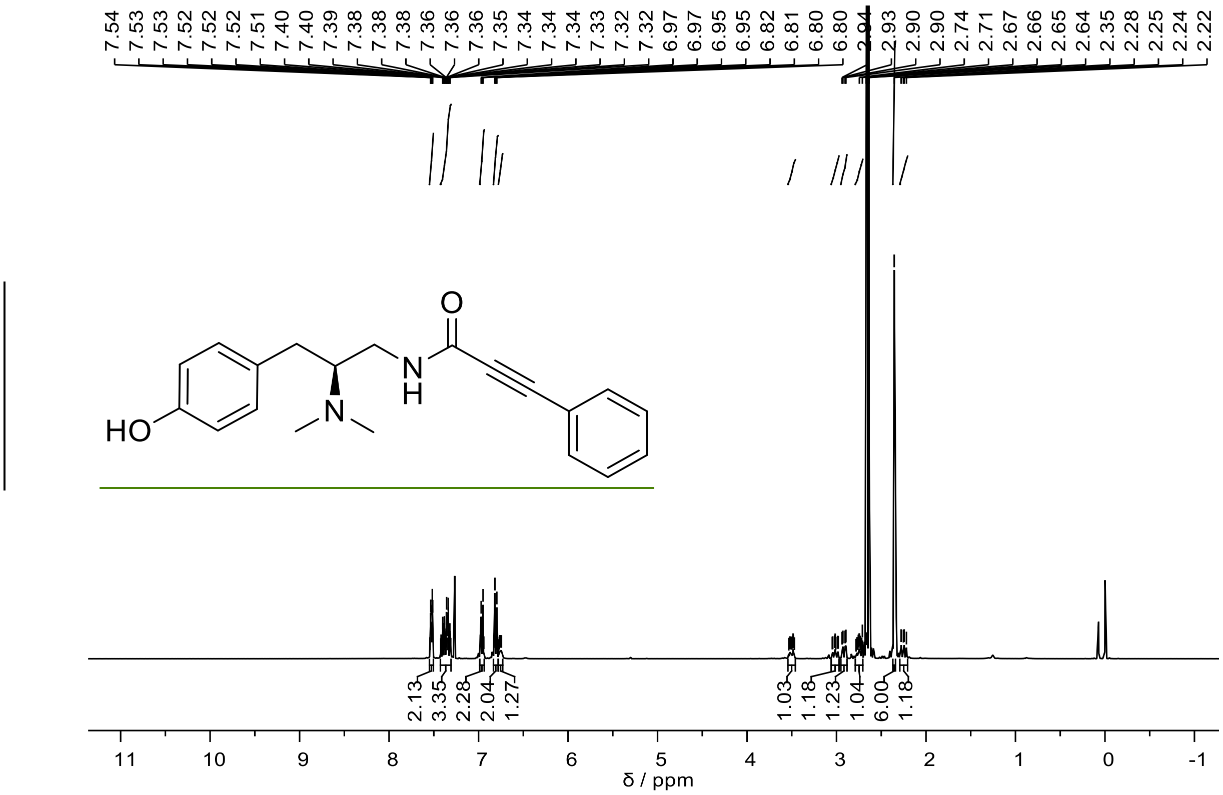


400MHz $^{1}$H NMR spectrum of **FH172** at 298K in $CDCl_3$.

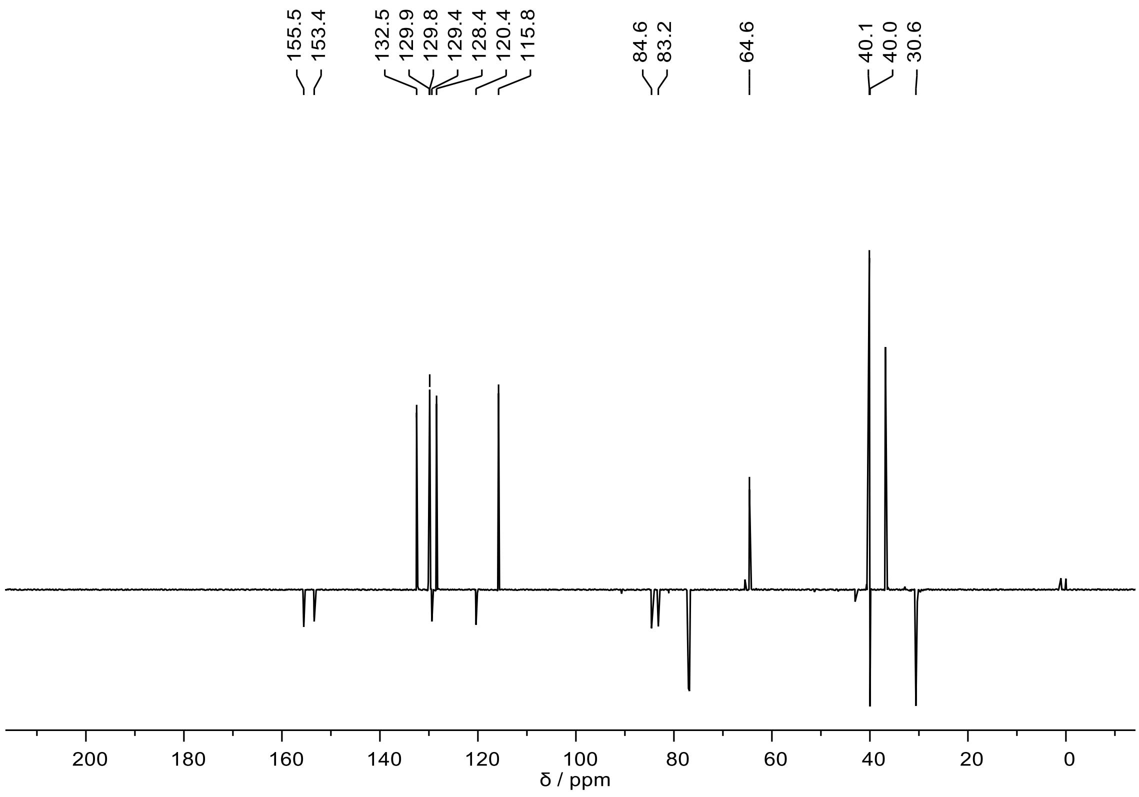


101MHz $^{13}$C NMR spectrum of **FH172** at 298K in $CDCl_3$.

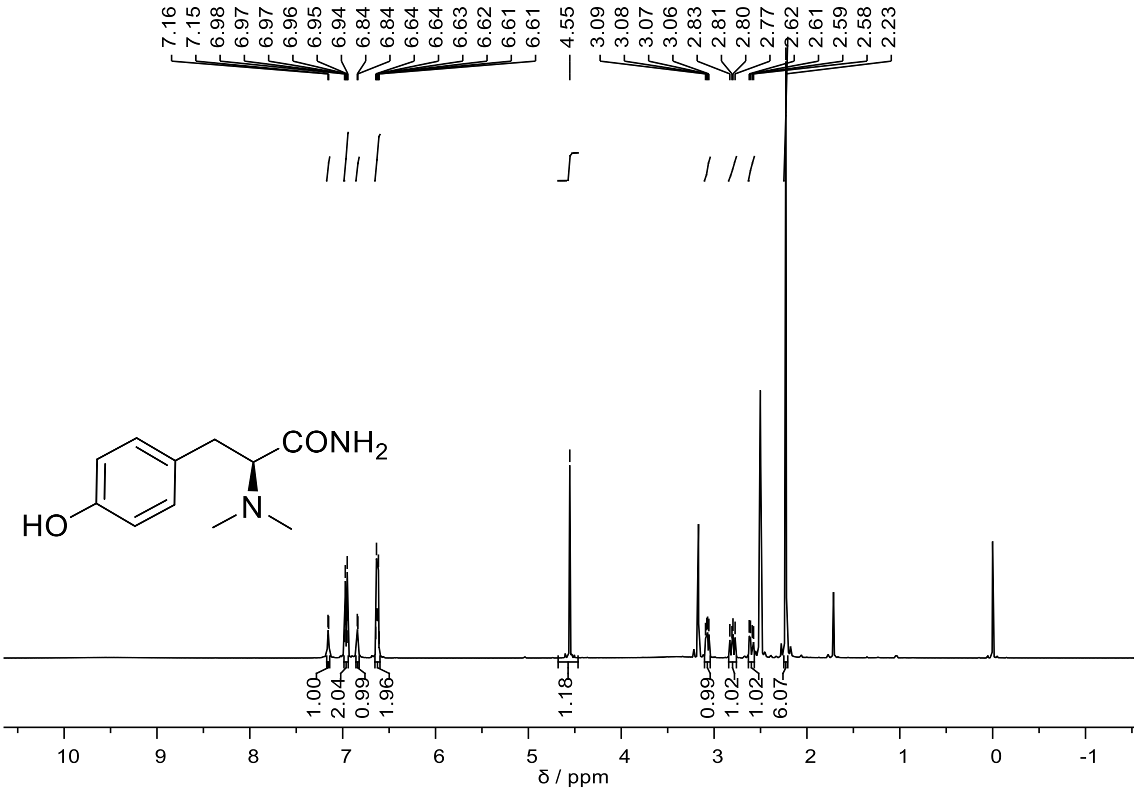


400MHz $^{1}$H NMR spectrum of **FH184** at 298K in DMSO-$d_6$.

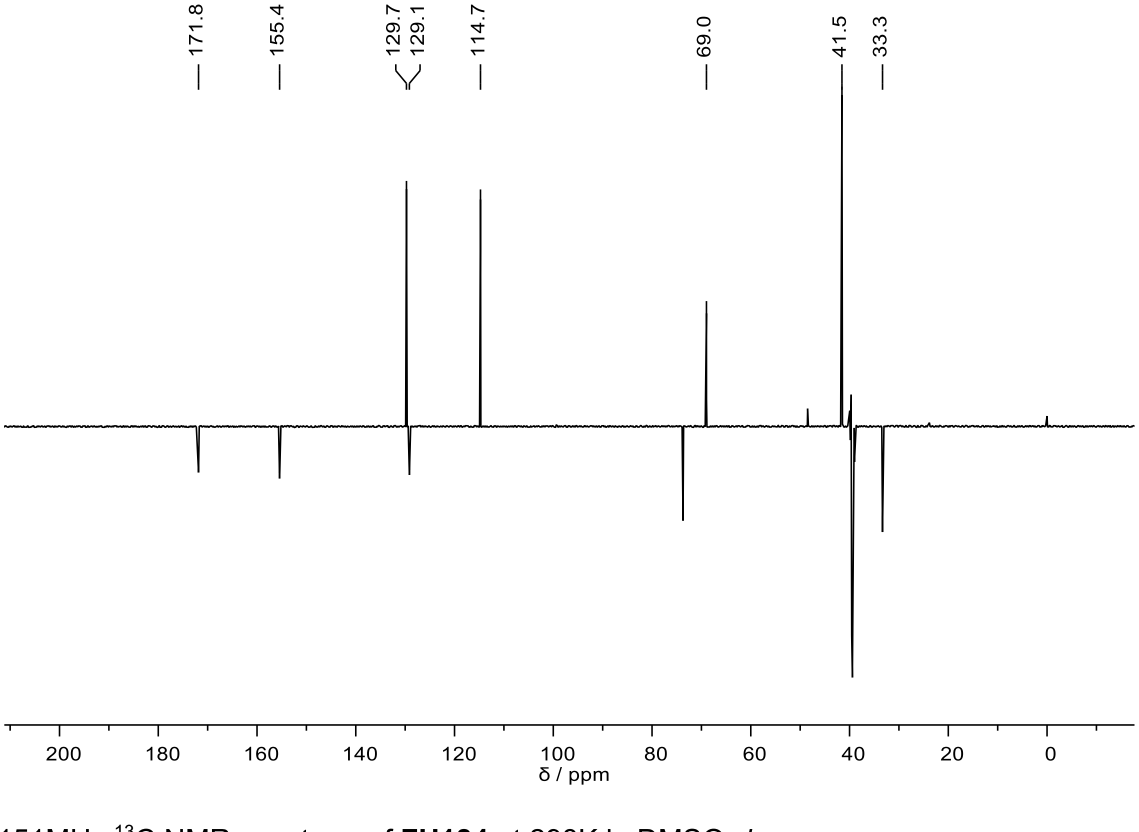


151MHz $^{13}$C NMR spectrum of **FH184** at 298K in DMSO-$d_6$.

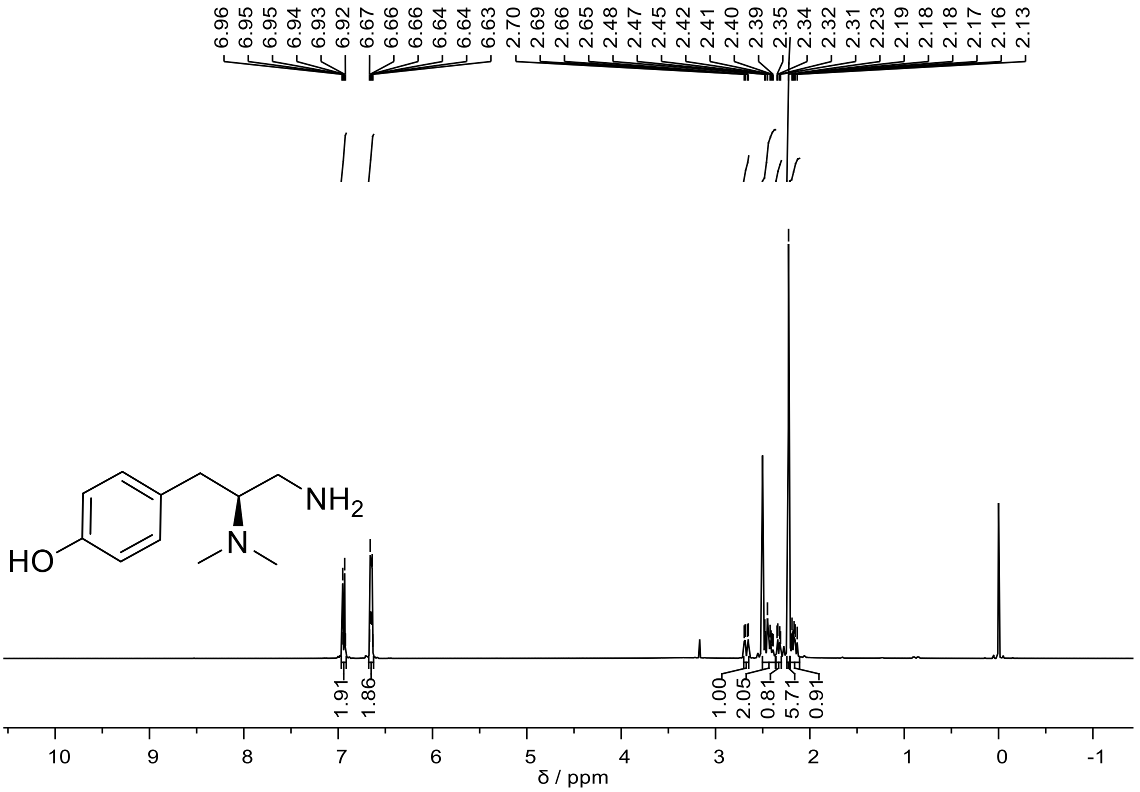


400MHz $^1$H NMR spectrum of **FH185** at 298K in DMSO-$d_6$.

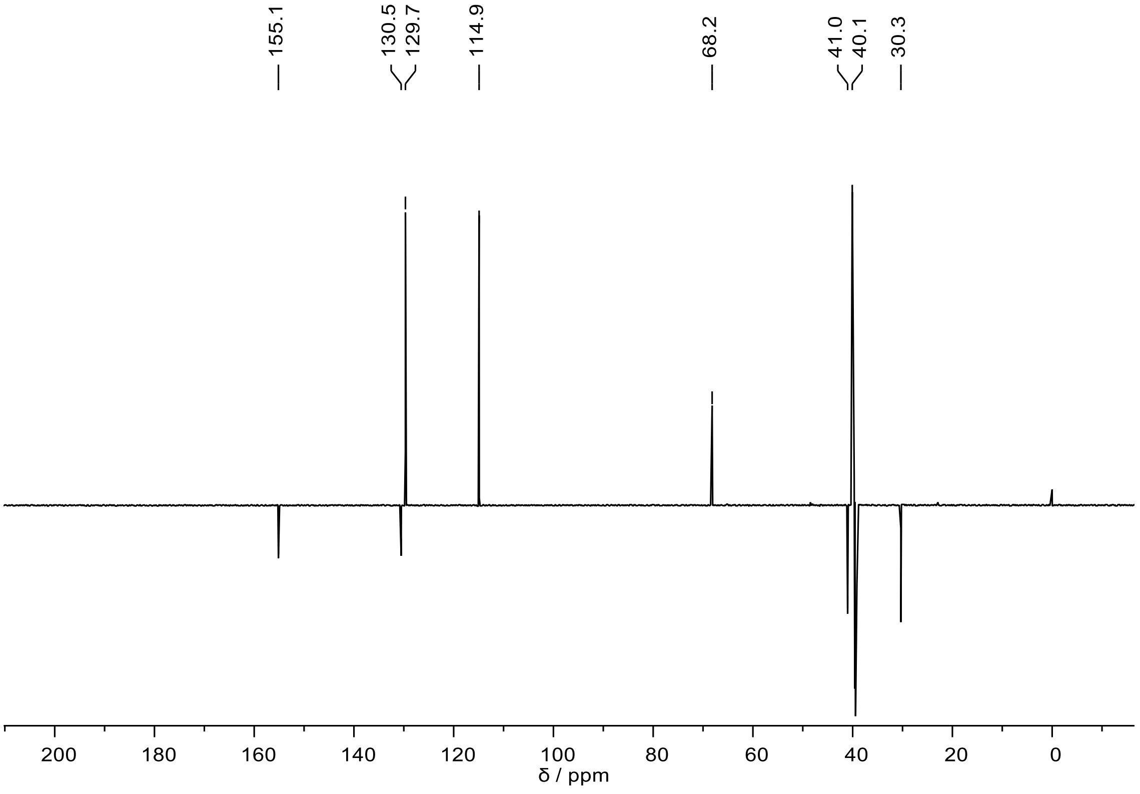


151MHz $^{13}$C NMR spectrum of **FH185** at 298K in DMSO-$d_6$.

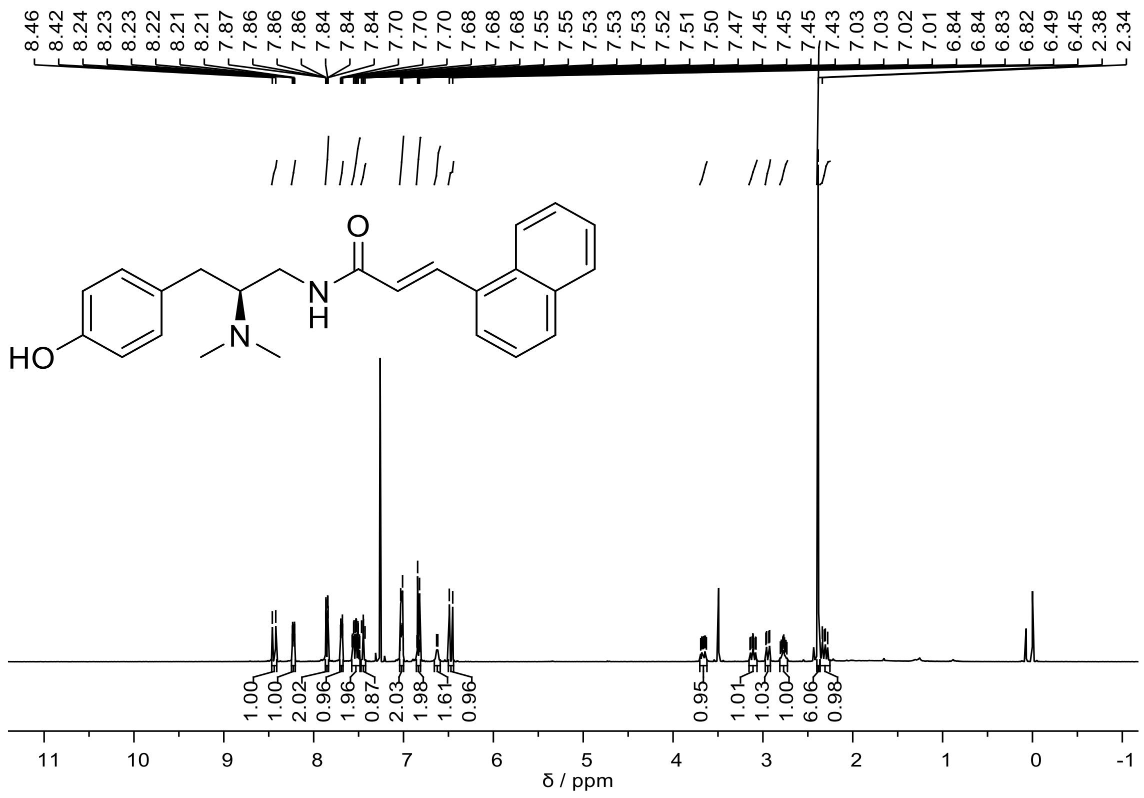


400MHz $^{1}$H NMR spectrum of **FH210** at 298K in $CDCl_3$.

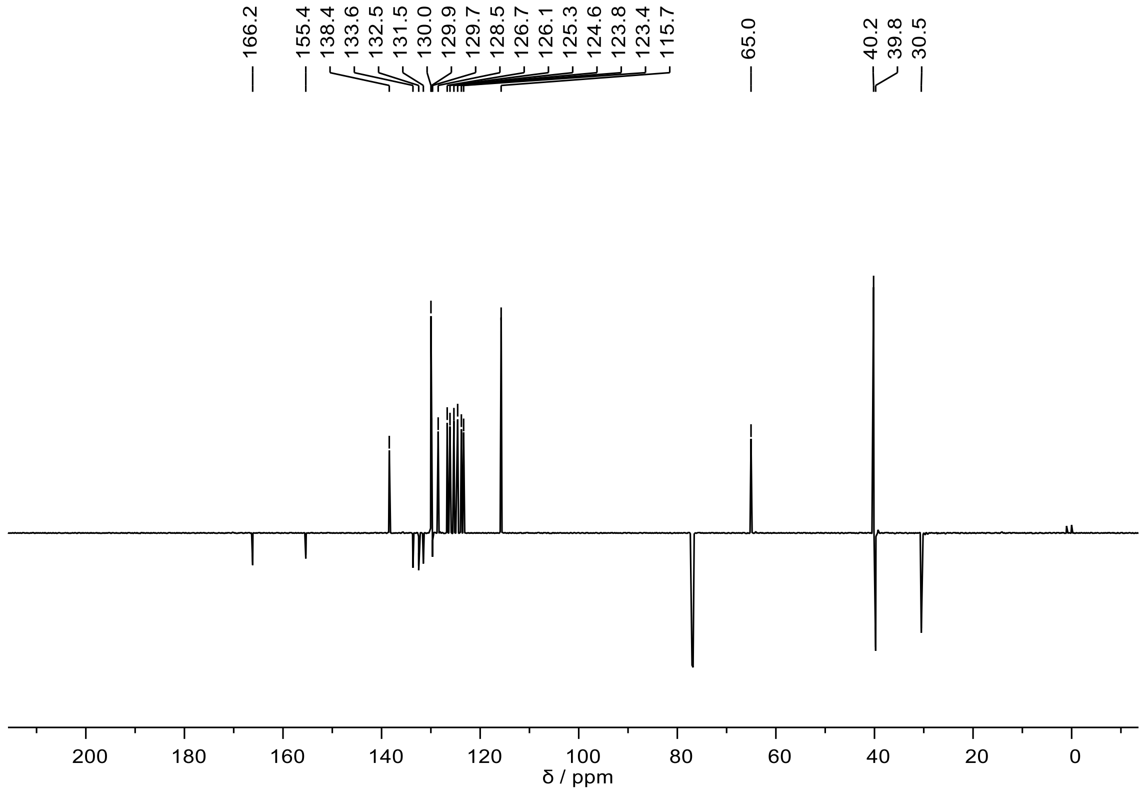


151MHz $^{13}$C NMR spectrum of **FH210** at 298K in $CDCl_3$.

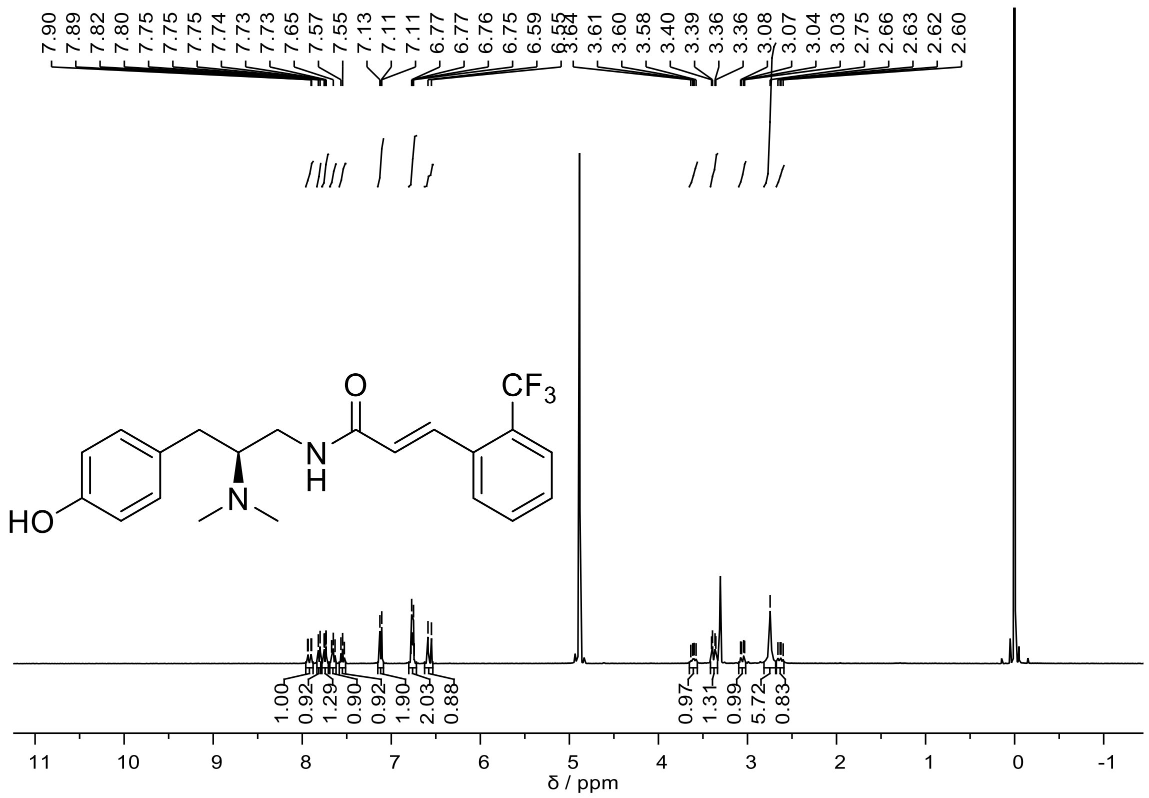

400MHz $^{1}H$ NMR spectrum of **FH217** at 298K in $CD_3OD$.

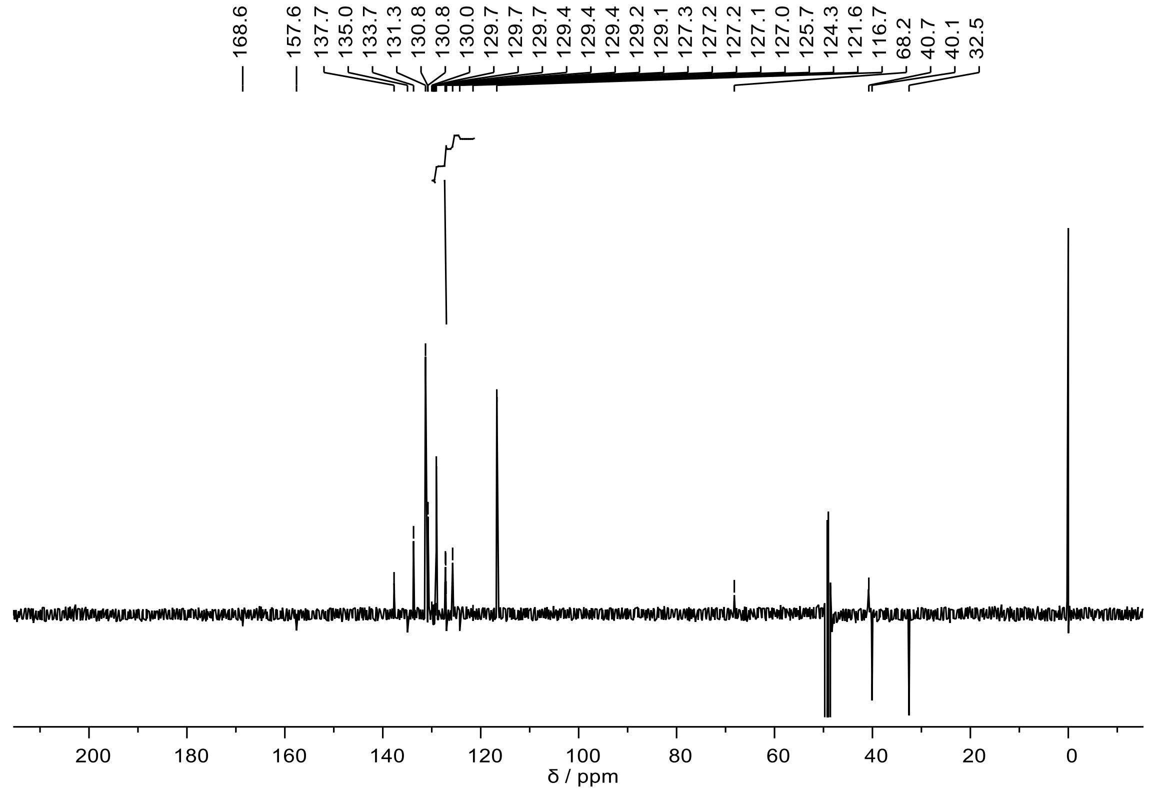

101MHz $^{13}C$ NMR spectrum of **FH217** at 298K in $CD_3OD$.

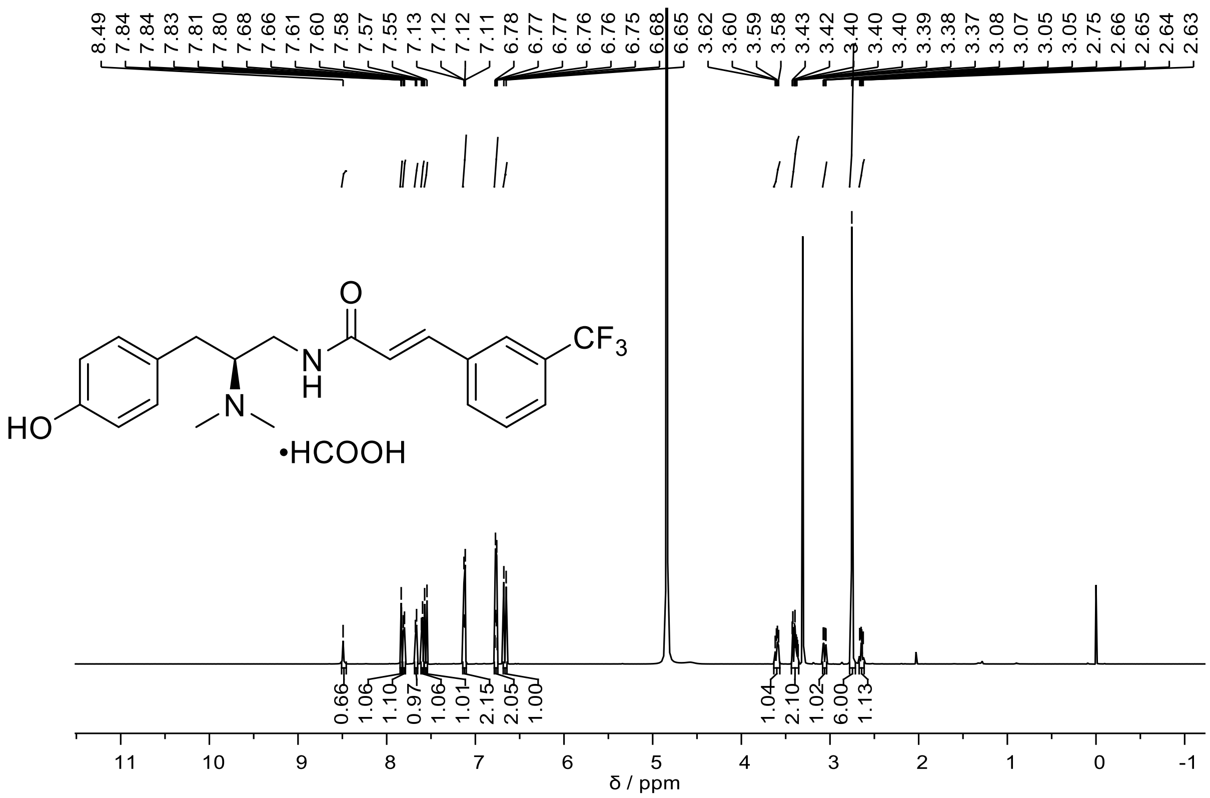


600MHz $^1$H NMR spectrum of **FH218** at 298K in $CD_3OD$.

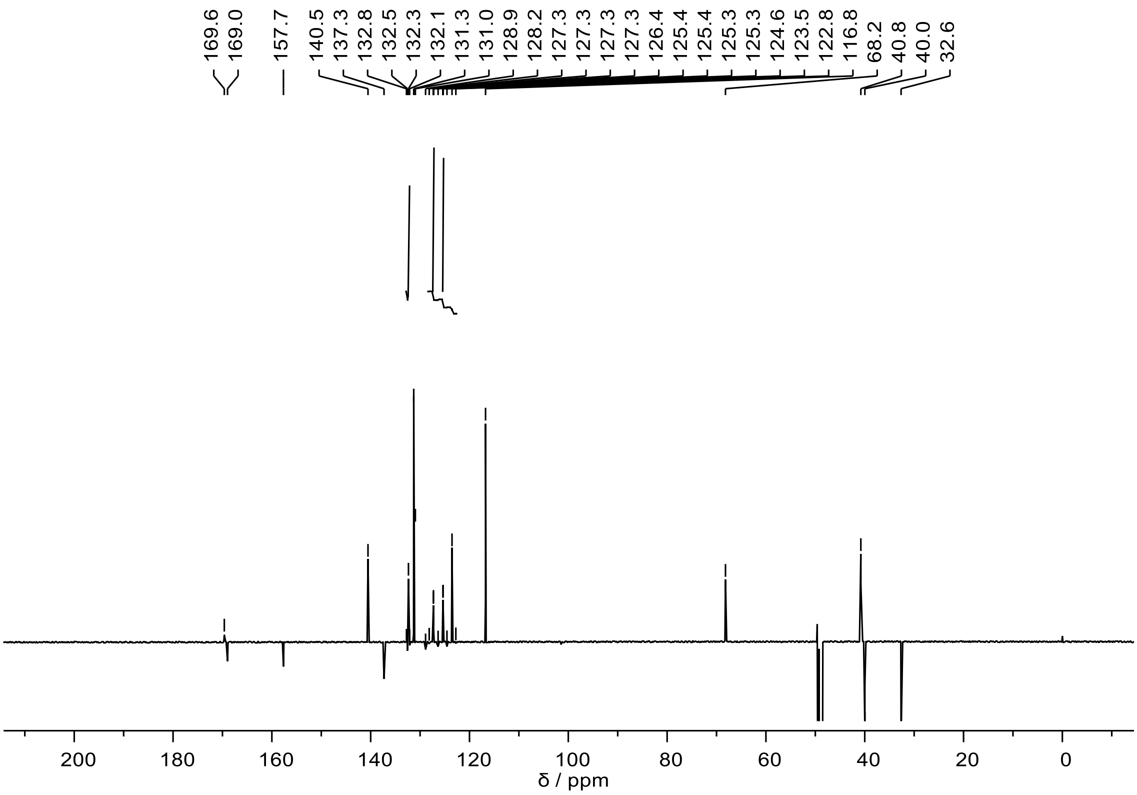


151MHz $^{13}$C NMR spectrum of **FH218** at 298K in $CD_3OD$.

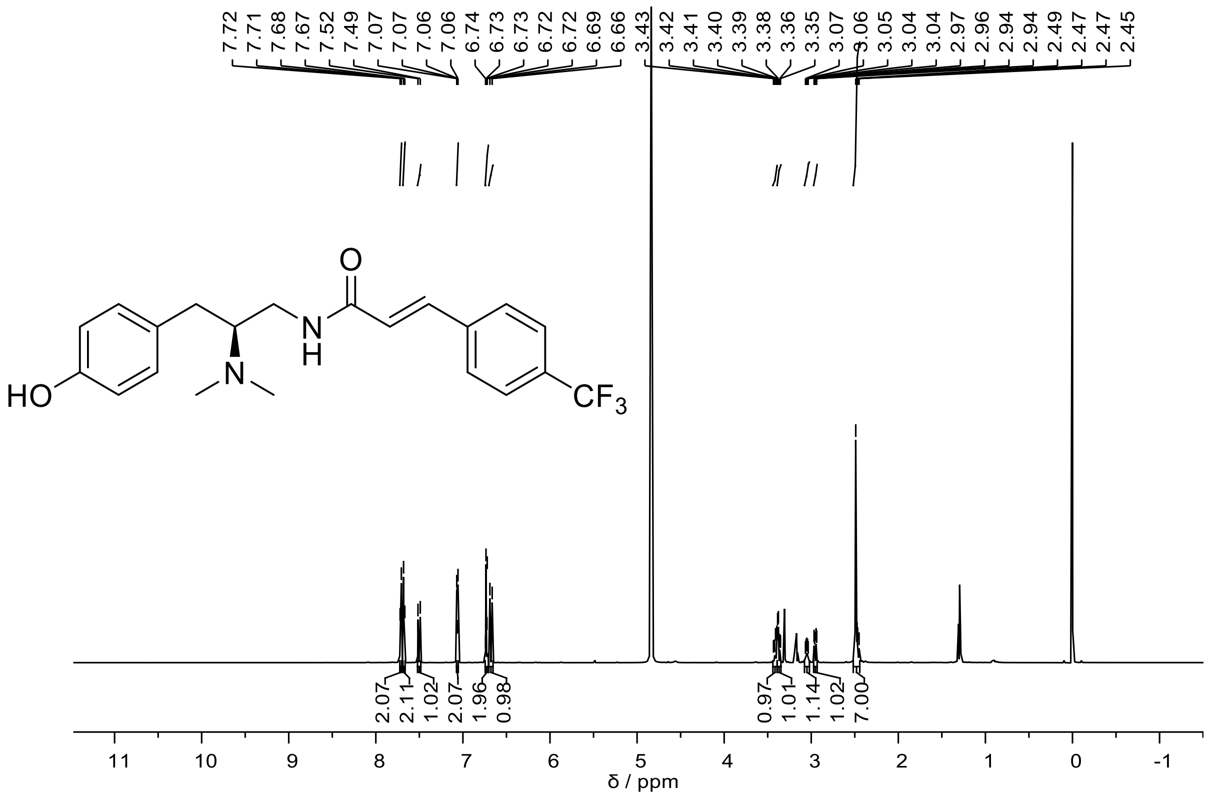


600MHz $^1$H NMR spectrum of **FH219** at 298K in $CD_3OD$.

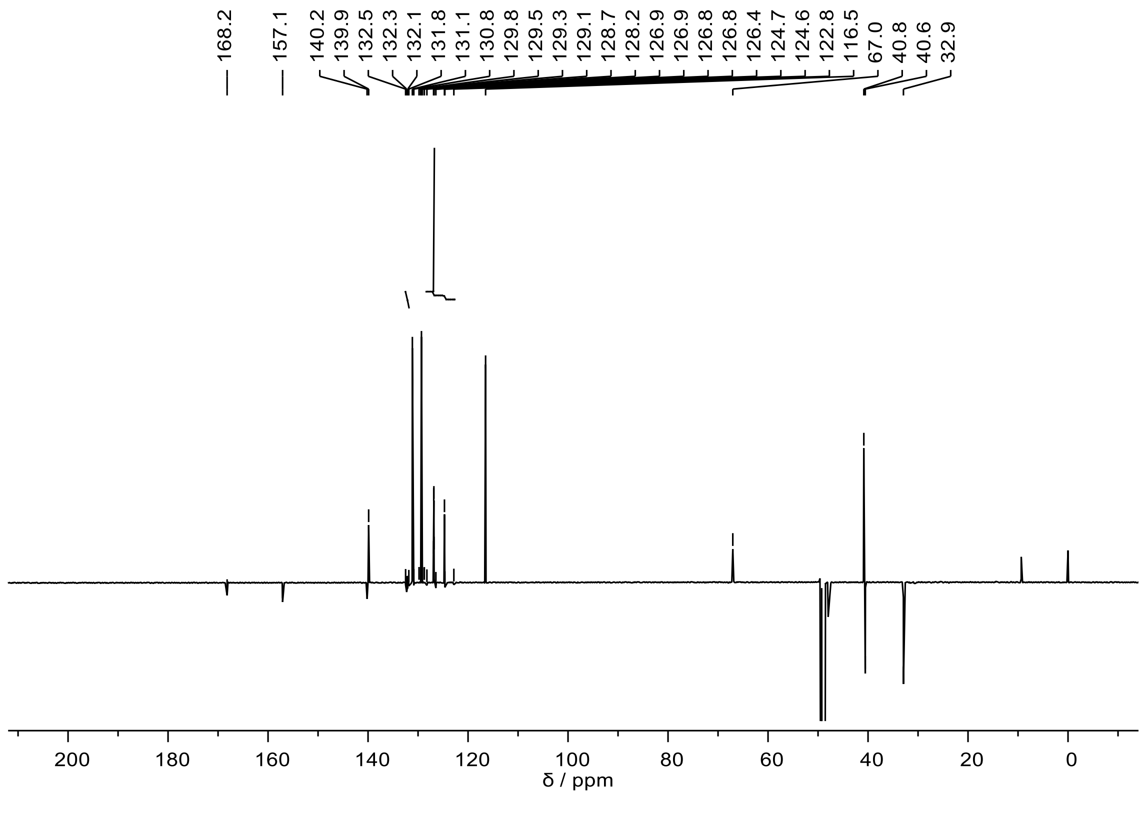


151MHz $^{13}$C NMR spectrum of **FH219** at 298K in $CD_3OD$.

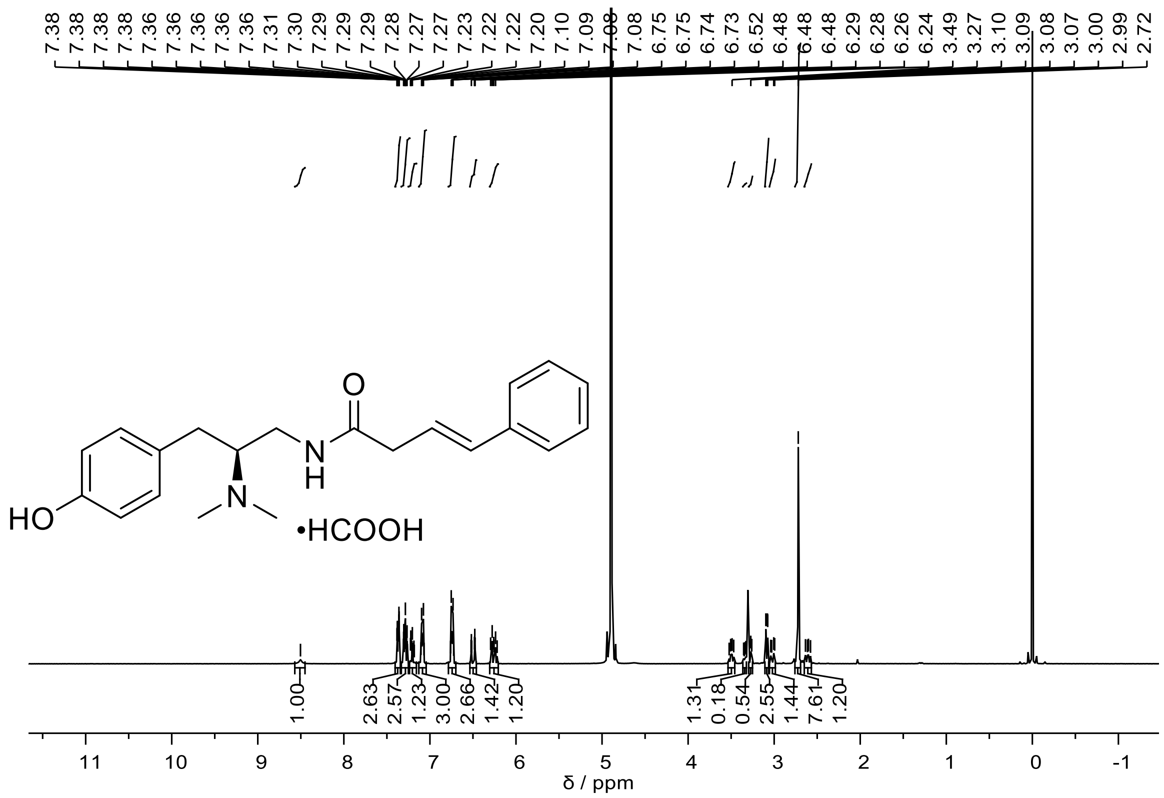


400MHz $^{1}H$ NMR spectrum of **FH223** at 298K in $CD_3OD$.

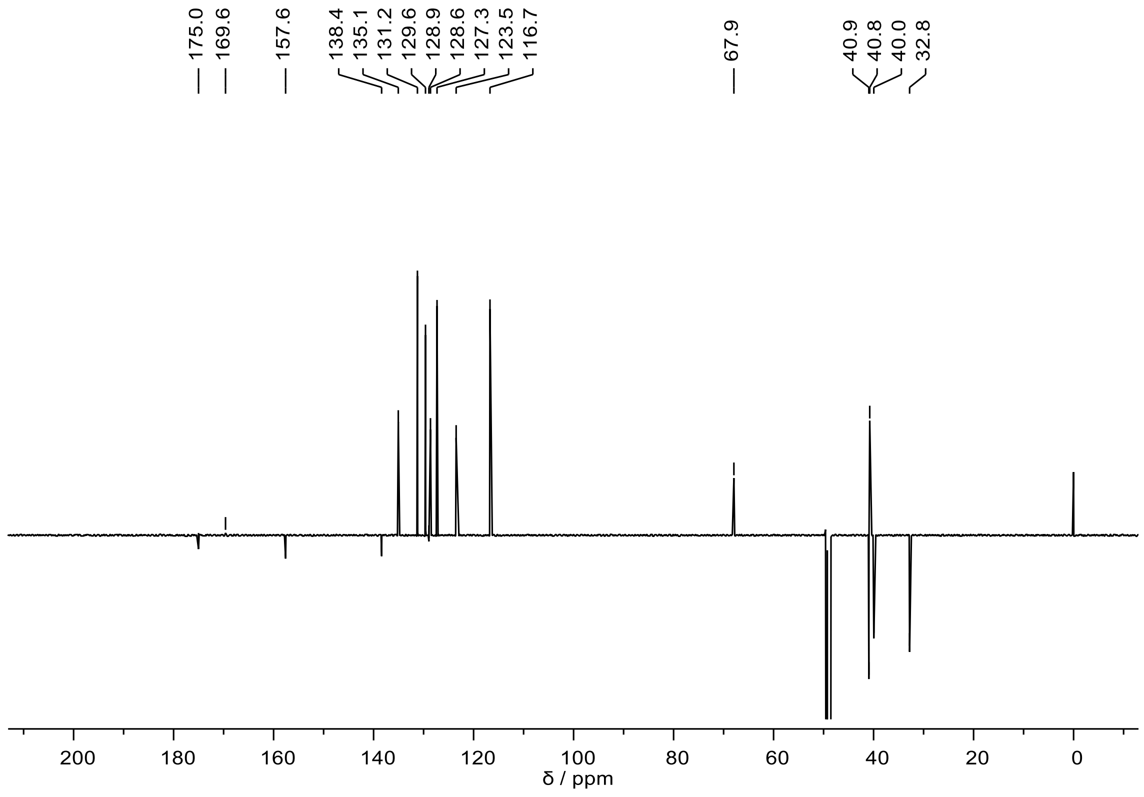


151MHz $^{13}C$ NMR spectrum of **FH223** at 298K in $CD_3OD$.

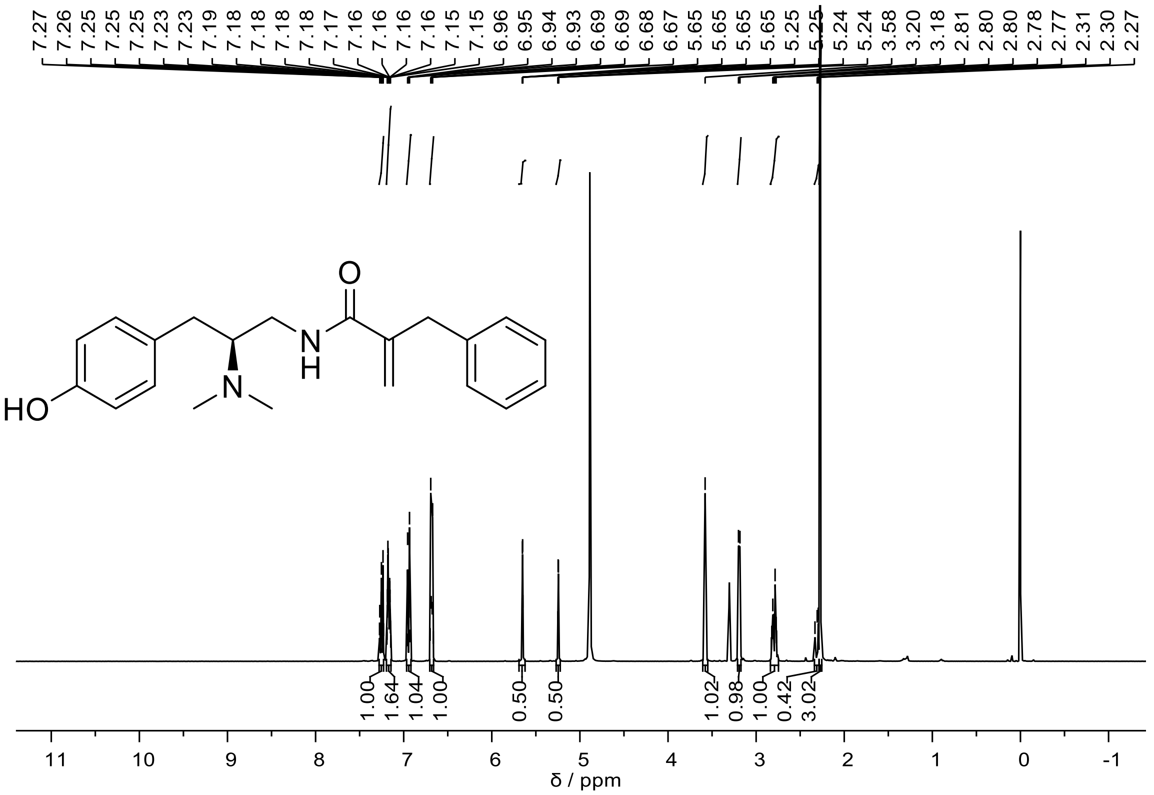


400MHz $^{1}$H NMR spectrum of **FH273** at 298K in $CD_3OD$.

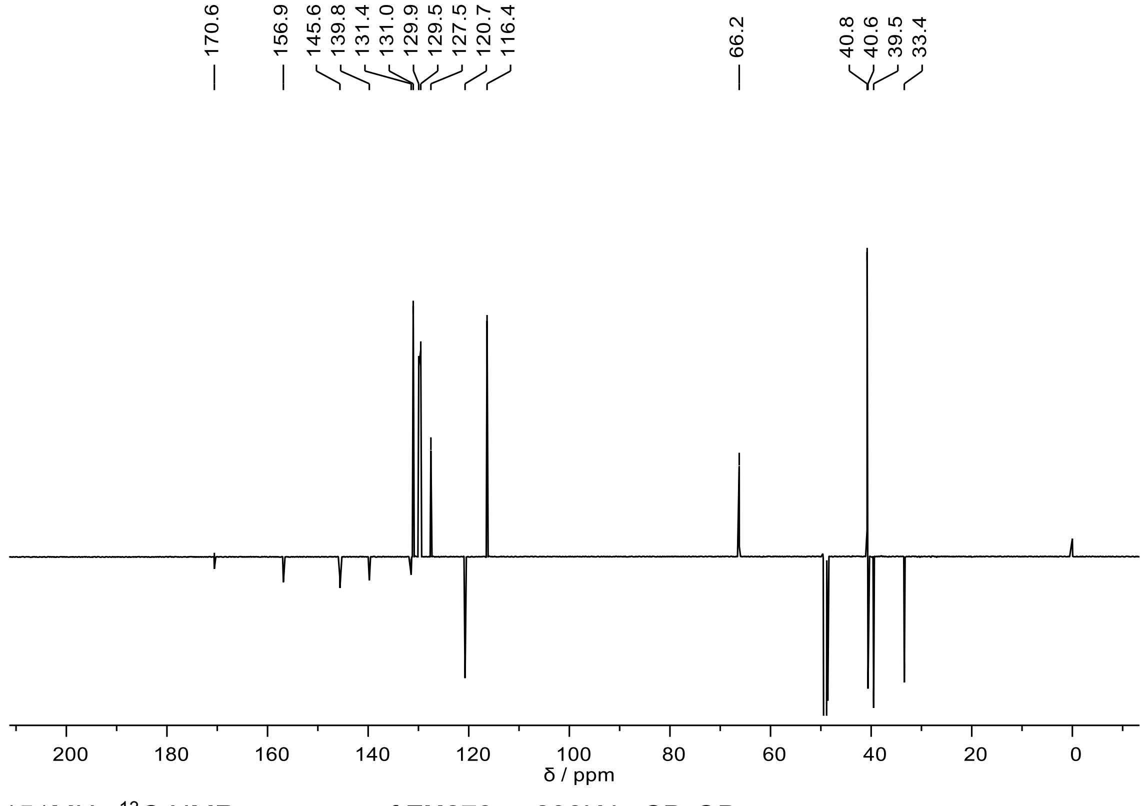


151MHz $^{13}$C NMR spectrum of **FH273** at 298K in $CD_3OD$.

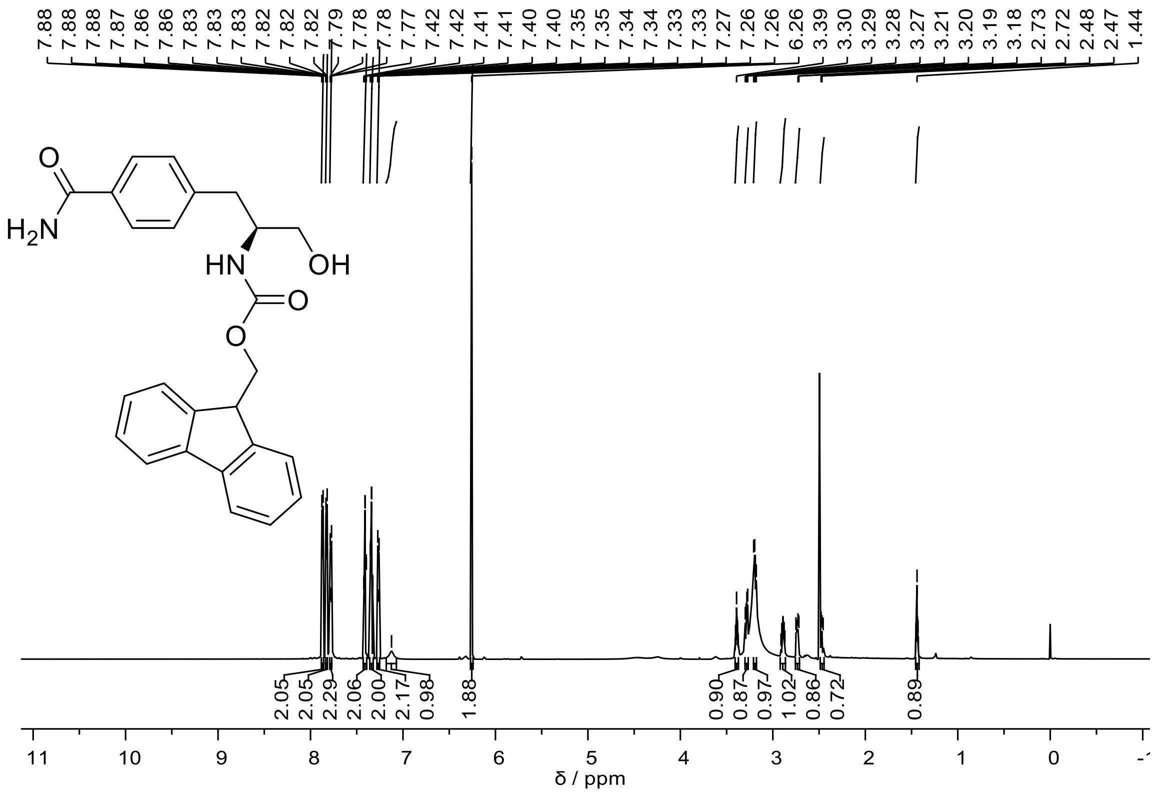


600MHz $^{1}$H NMR spectrum of **FH282** at 318K in DMSO-$d_6$.

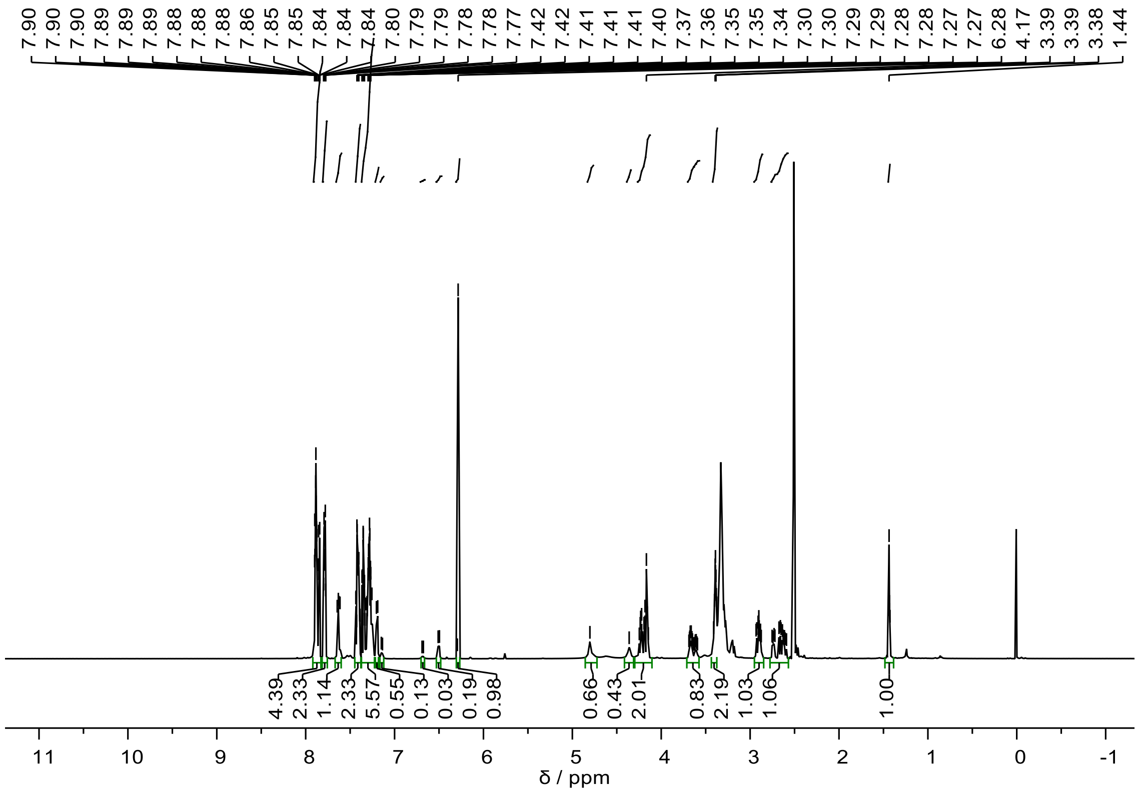


600MHz $^{1}$H NMR spectrum of **FH282** at 298K in DMSO-$d_6$.

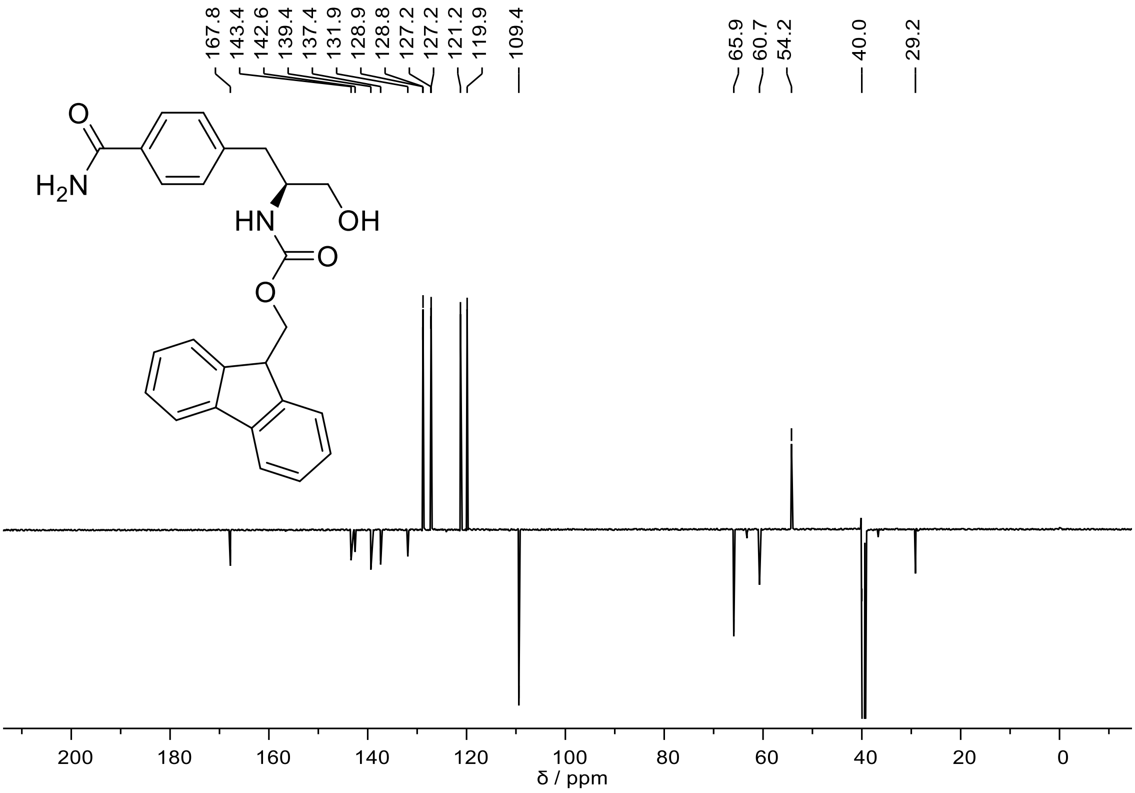


151MHz $^{13}$C NMR spectrum of **FH282** at 318K in DMSO-$d_6$.

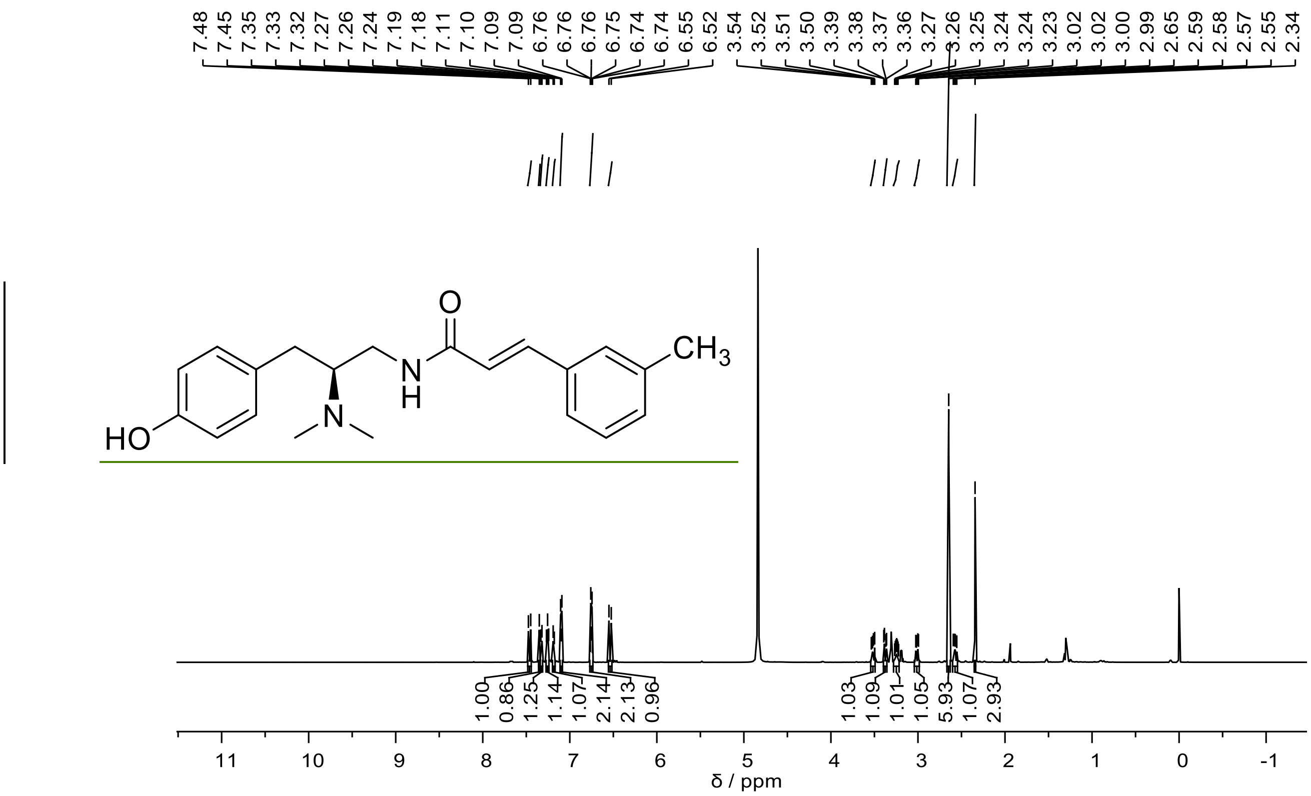


600MHz $^{1}$H NMR spectrum of **FH299** at 298K in $CD_3OD$.

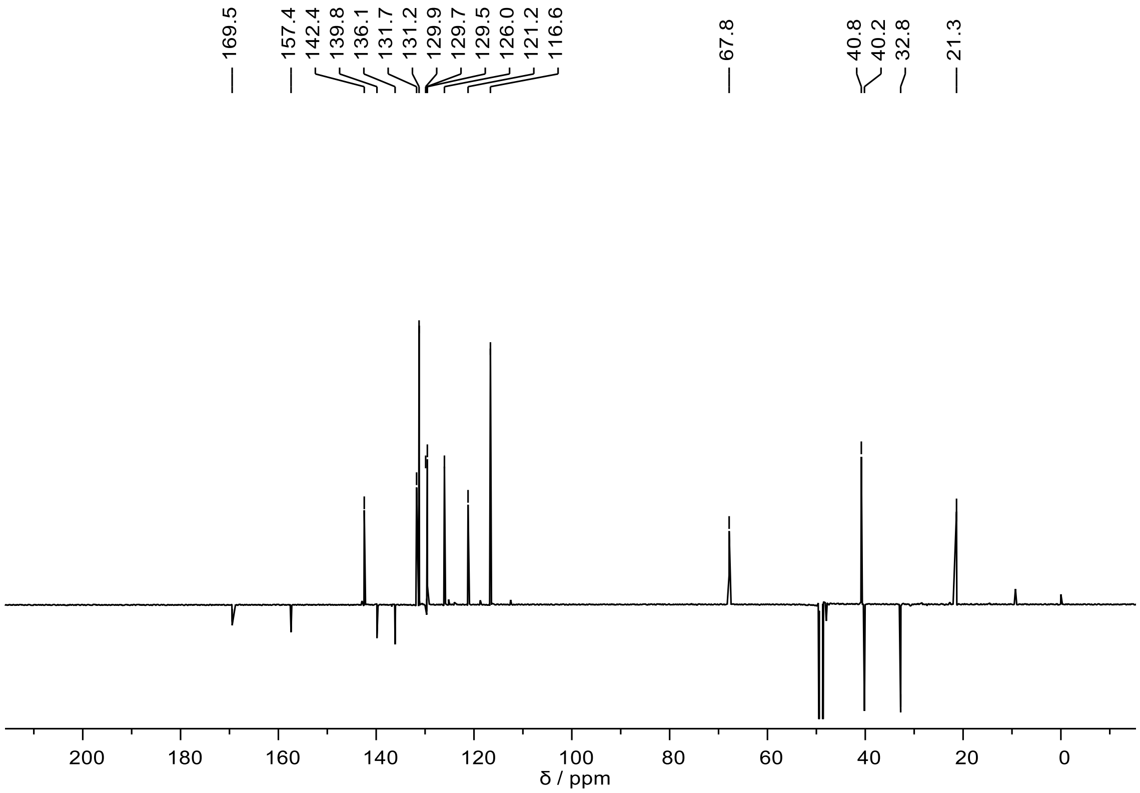


151MHz $^{13}$C NMR spectrum of **FH299** at 298K in $CD_3OD$.

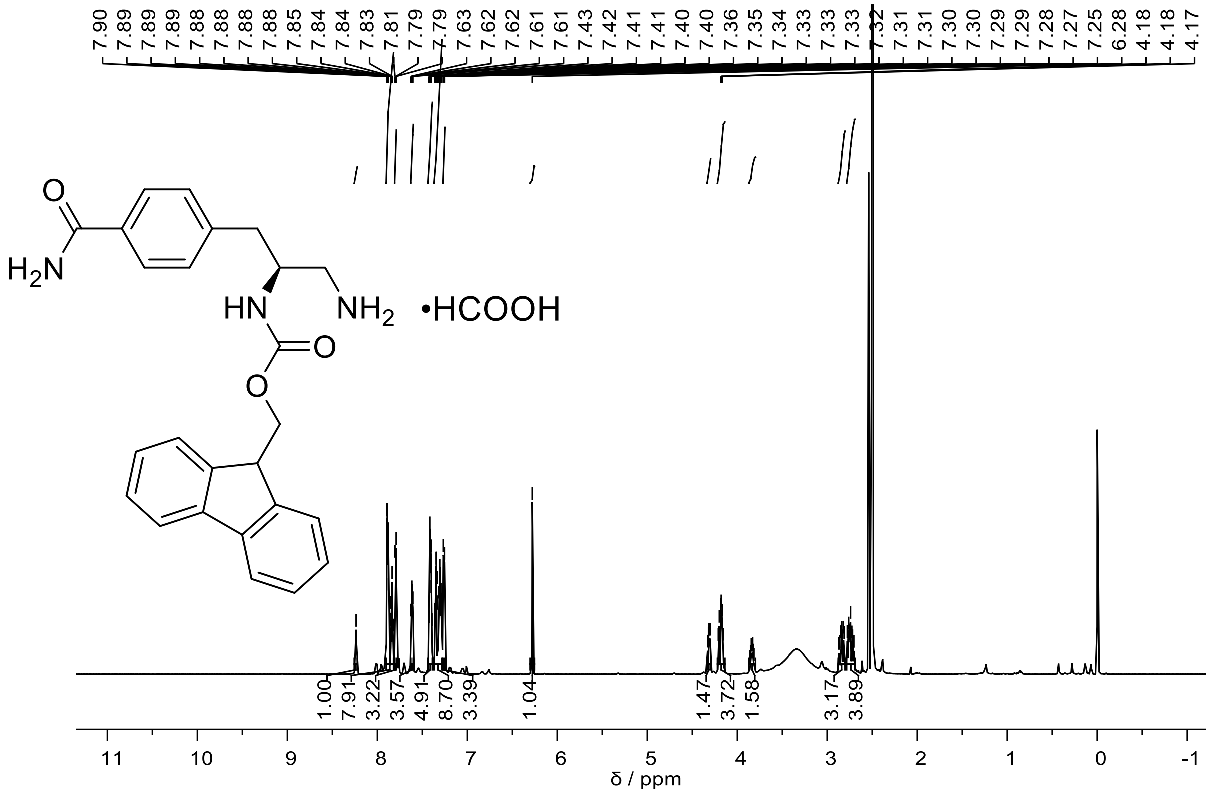


600MHz $^{1}$H NMR spectrum of **FH305** at 298K in DMSO-$d_6$.

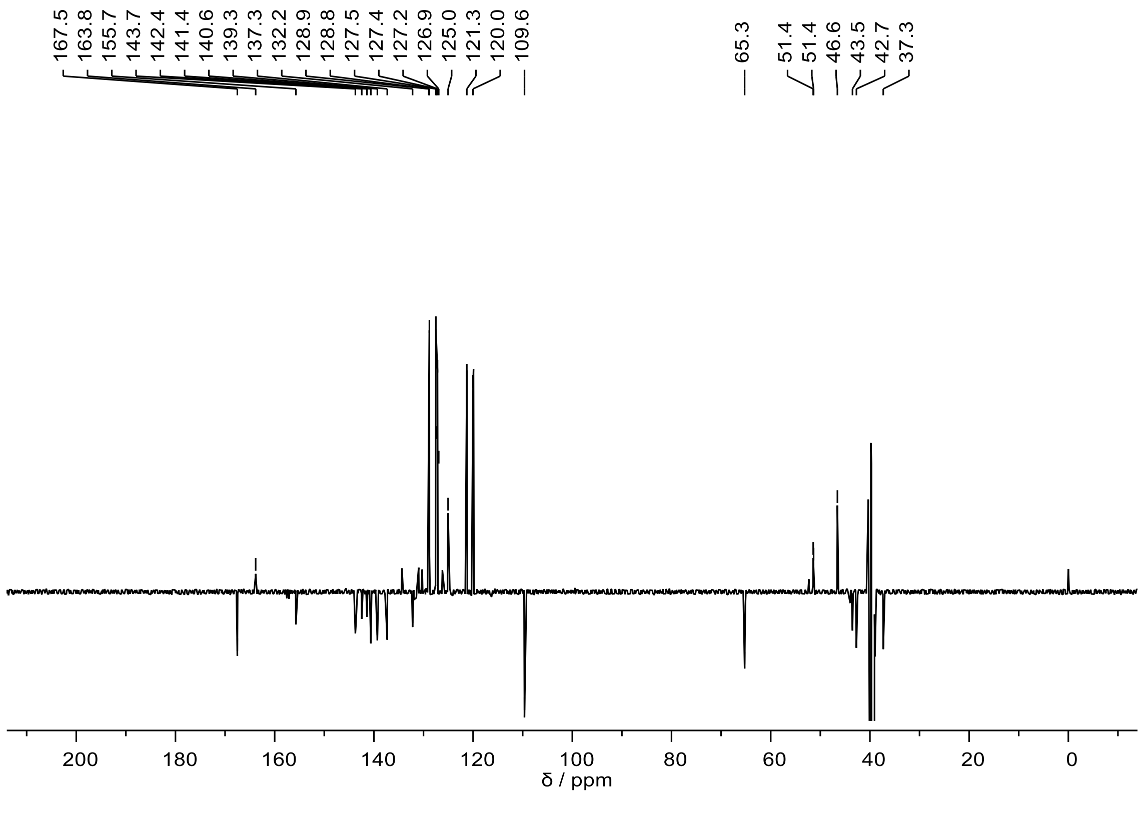

151MHz $^{13}C$ NMR spectrum of **FH305** at 298K in DMSO-$d_6$.

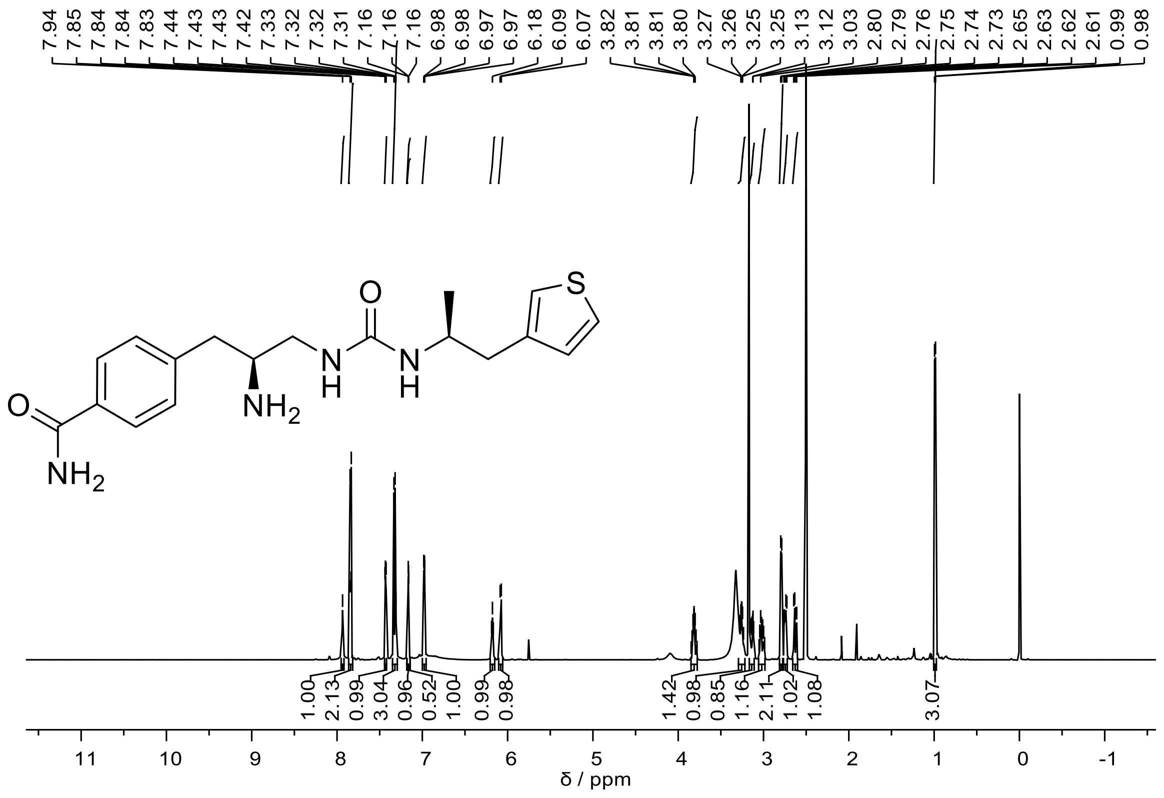

600MHz $^{1}$H NMR spectrum of **FH308** at 298K in DMSO-$d_6$.

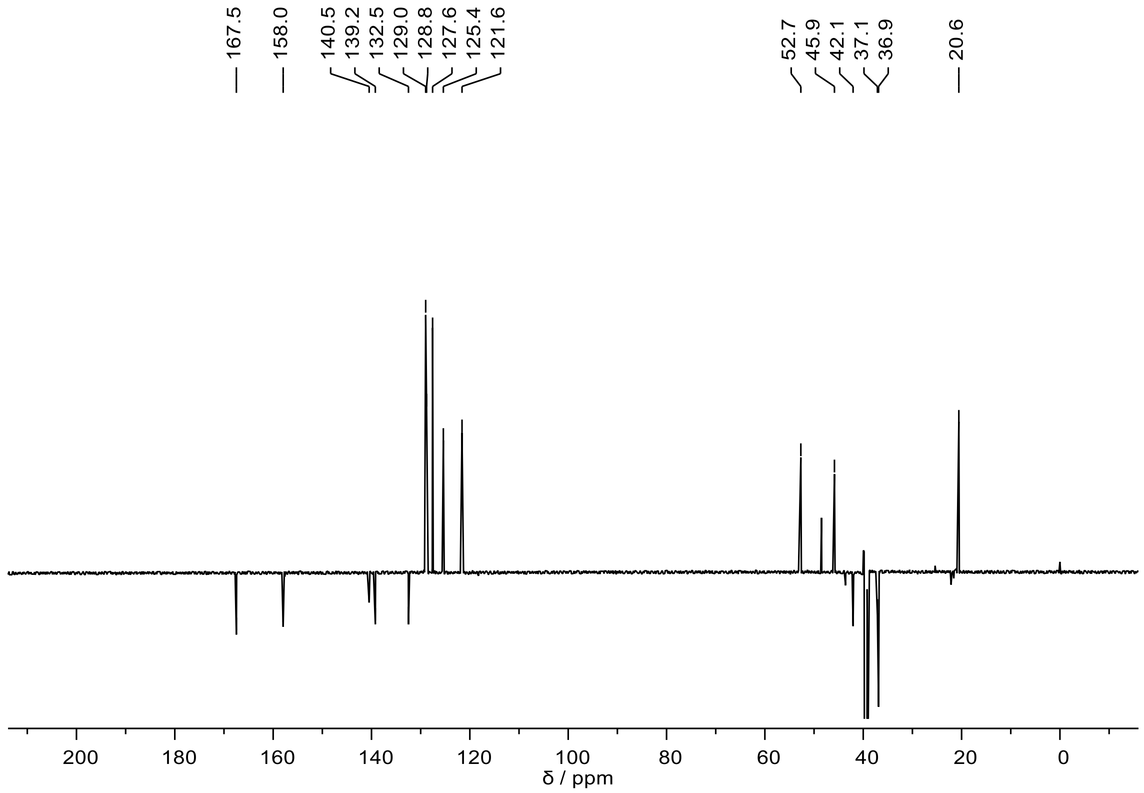

151MHz $^{13}$C NMR spectrum of **FH308** at 298K in DMSO-$d_6$.

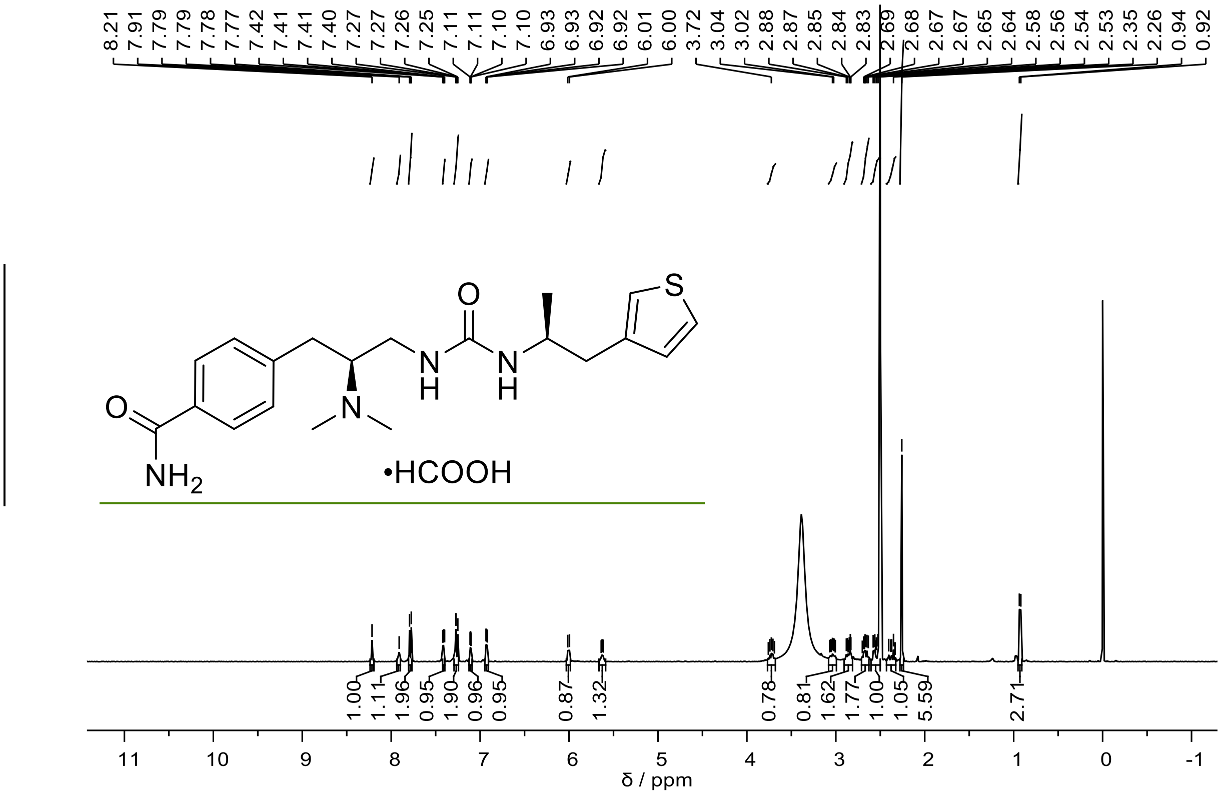


400MHz $^{1}H$ NMR spectrum of **FH310** at 298K in DMSO-$d_6$.

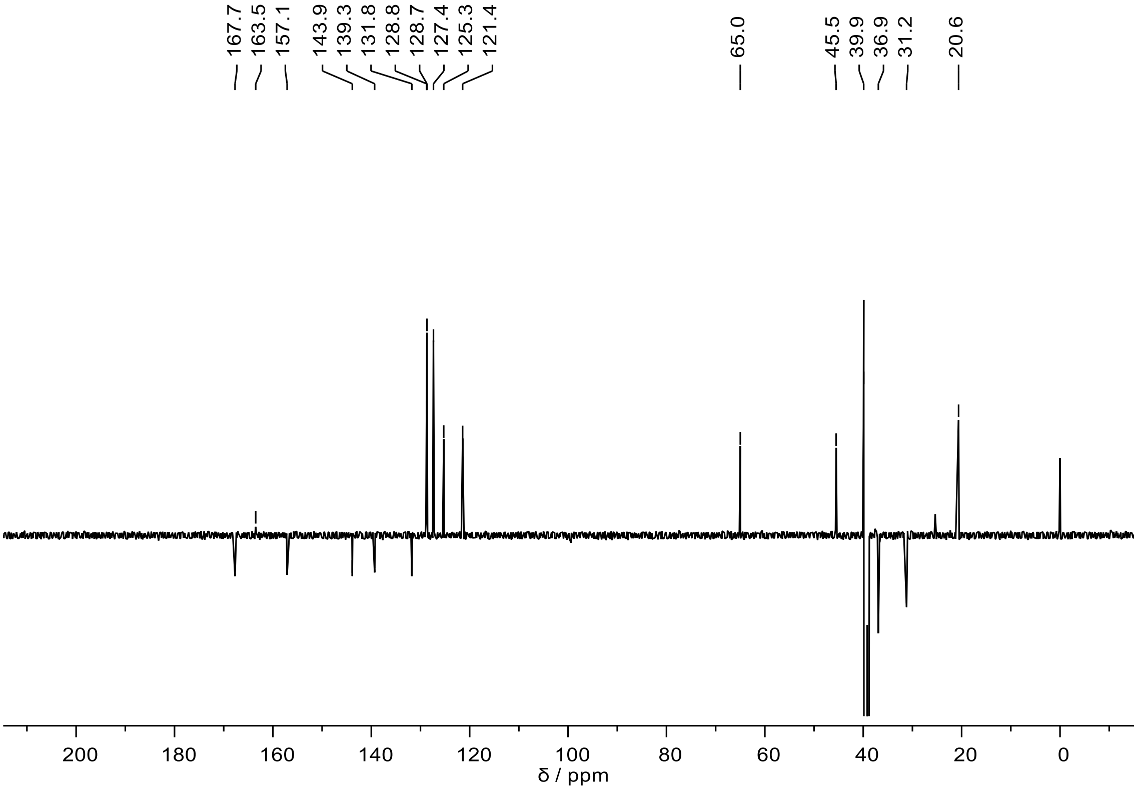


151MHz $^{13}C$ NMR spectrum of **FH310** at 298K in DMSO-$d_6$.

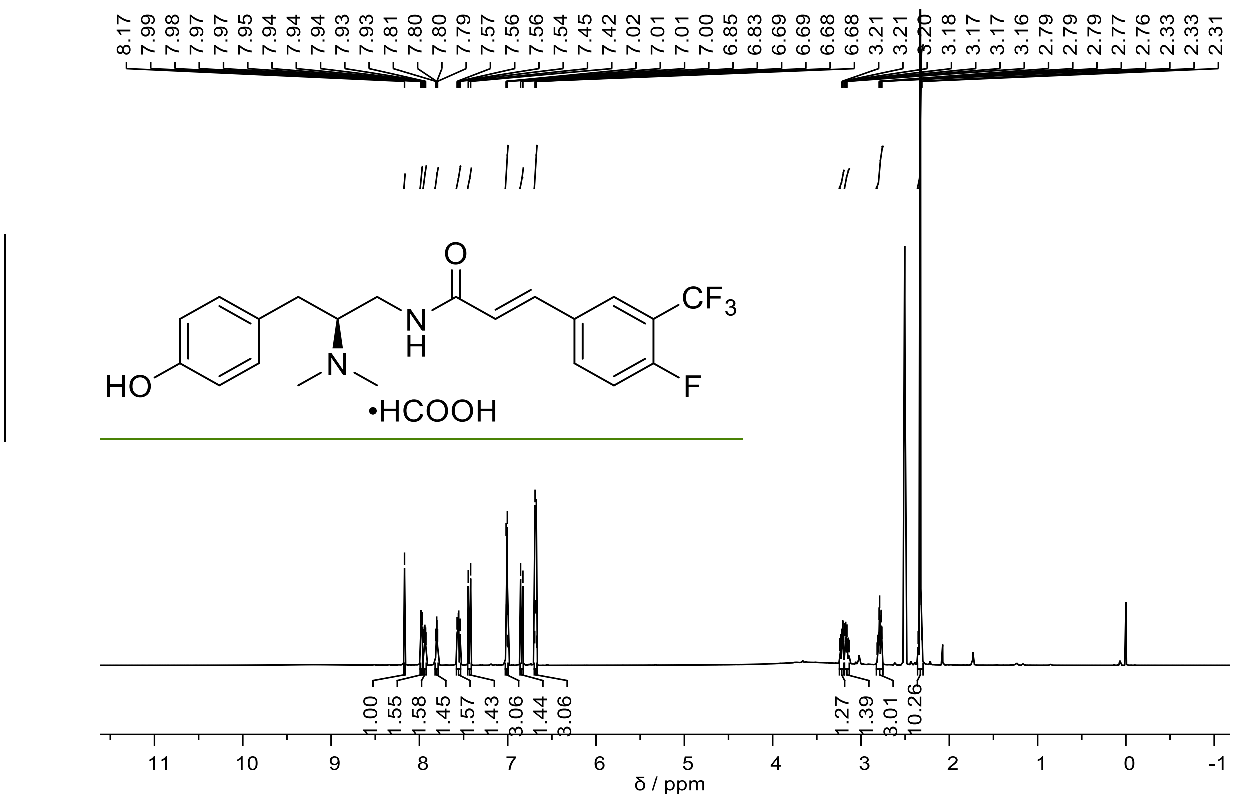


600MHz $^{1}$H NMR spectrum of **FH315** at 298K in DMSO-$d_6$.

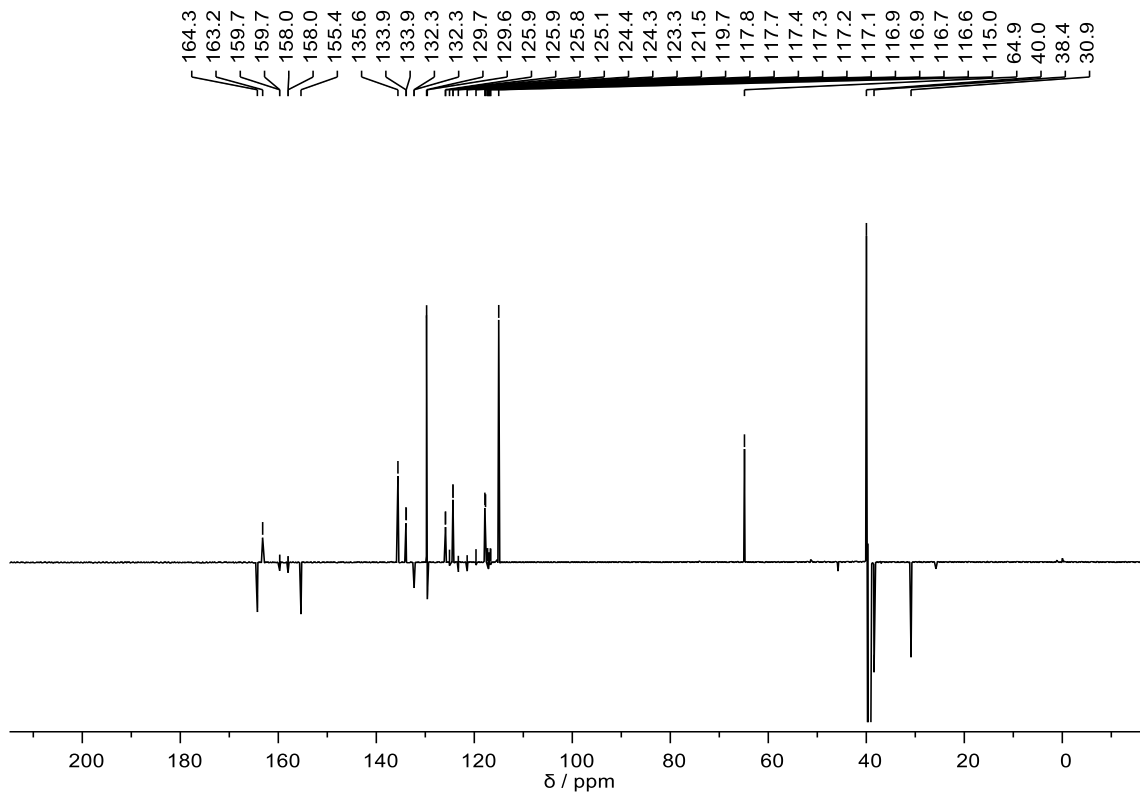


151MHz $^{13}$C NMR spectrum of **FH315** at 298K in DMSO-$d_6$.

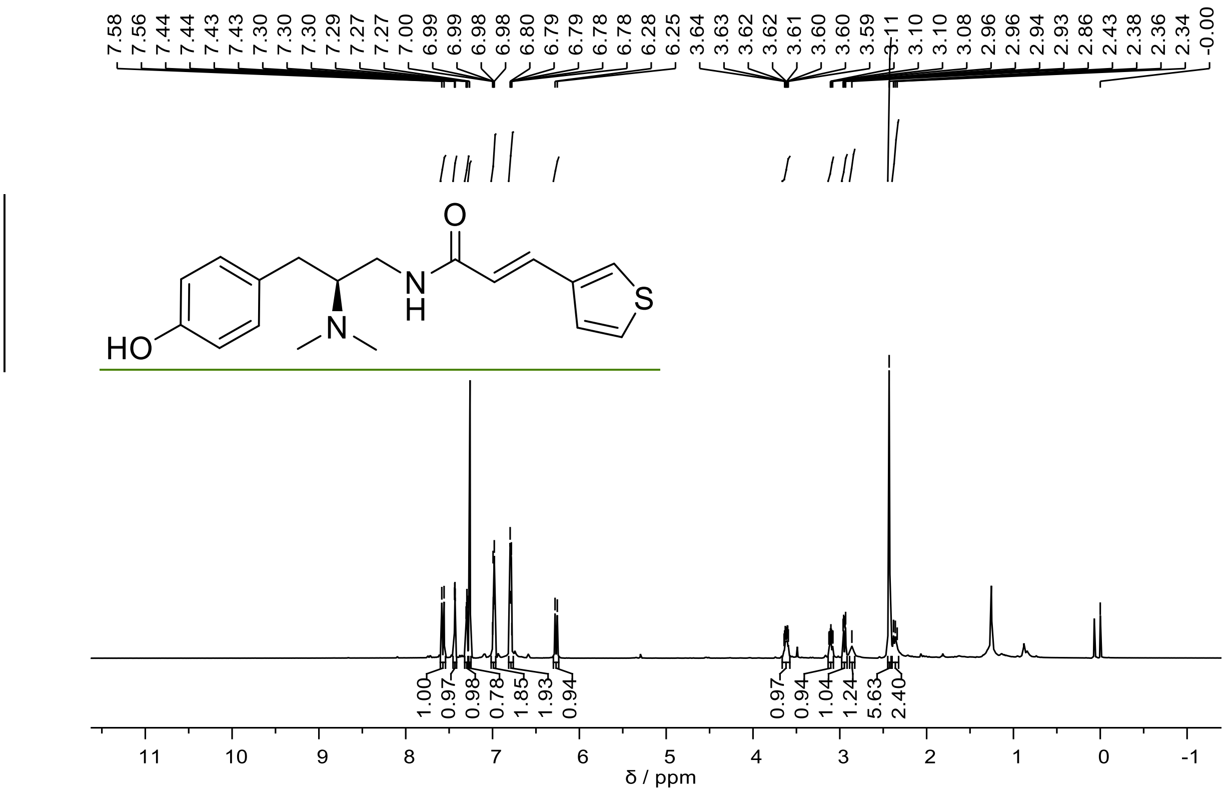

600MHz $^{1}$H NMR spectrum of **FH321** at 298K in DMSO-$d_6$.

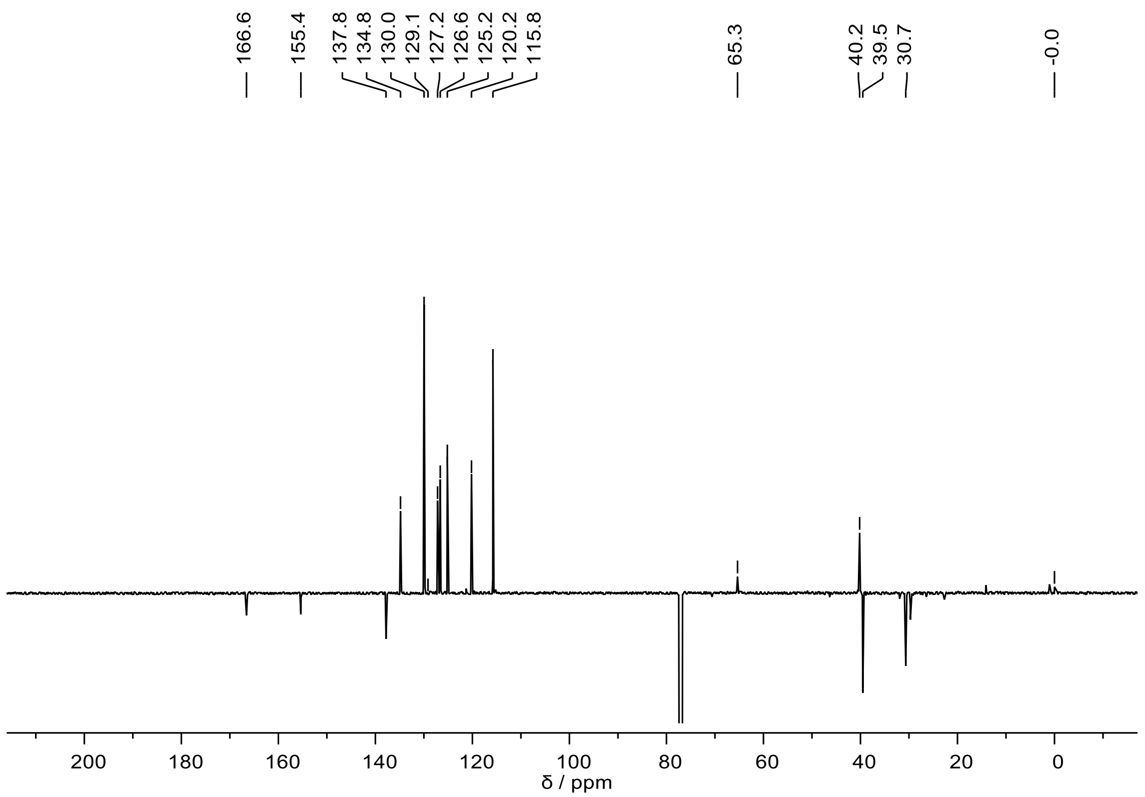

151MHz $^{13}$C NMR spectrum of **FH321** at 298K in DMSO-$d_6$.
